# The neuronal calcium sensor Synaptotagmin-1 and SNARE proteins cooperate to dilate fusion pores

**DOI:** 10.1101/623827

**Authors:** Zhenyong Wu, Nadiv Dharan, Zachary A. McDargh, Sathish Thiyagarajan, Ben O’Shaughnessy, Erdem Karatekin

## Abstract

All membrane fusion reactions proceed through an initial fusion pore, including calcium-triggered release of neurotransmitters and hormones. Expansion of this small pore to release cargo is energetically costly and regulated by cells, but the mechanisms are poorly understood. Here we show that the neuronal/exocytic calcium sensor Synaptotagmin-1 (Syt1) promotes expansion of fusion pores induced by SNARE proteins. Pore dilation relied on calcium-induced insertion of the tandem C2 domain hydrophobic loops of Syt1 into the membrane, previously shown to reorient the C2 domain. Mathematical modelling suggests that C2B reorientation rotates a bound SNARE complex so that it exerts force on the membranes in a mechanical lever action that increases the height of the fusion pore, provoking pore dilation to offset the bending energy penalty. We conclude that Syt1 exerts novel non-local calcium-dependent mechanical forces on fusion pores that dilate pores and assist neurotransmitter and hormone release.

**SIGNIFICANCE STATEMENT:** During neurotransmitter release, calcium-induced membrane insertion of the C2B domain of Synaptotagmin re-orients the bound SNARE complex which dilates the fusion pore in a mechanical lever action.

## INTRODUCTION

Release of neurotransmitters and hormones occurs through exocytosis in which neurotransmitter-filled synaptic vesicles or hormone-laden secretory vesicles fuse with the plasma membrane to release their cargo to the extracellular space (1). The initial merger of the vesicular and plasma membranes results in a narrow fusion pore only ~1 nm in diameter (2–5). Dynamics of this key intermediate determine release kinetics and the mode of vesicle recycling. The fusion pore can fluctuate in size, flicker open-closed multiple times and either reseal after partial release of contents or dilate for full cargo release. Because many endocrine cells co-package small and large cargoes, the pore can additionally act as a molecular sieve, controlling the type of cargo released. In pancreatic β-cells, fusion pores that fail to dilate release only small cargo such as ATP, but not insulin, a process that occurs more commonly in type 2 diabetes (6). Adrenal chromaffin cells release small catecholamines through flickering small pores, or release additional, larger cargo, in an activity-dependent manner (7). Fusion pore dynamics also affect release of neurotransmitters and the mode of endocytosis during synaptic vesicle fusion (5, 8–14).

Little is known about the molecular mechanisms that control pore dilation. SNARE proteins, a core component of the release machinery, are known to influence fusion pore dynamics (15–23). Formation of complexes between the vesicular v-SNARE VAMP2/Syb2 and plasma membrane t-SNAREs Syntaxin-1/SNAP25 is required for fusion (24). Insertion of flexible linkers between the SNARE domain and the transmembrane domain in VAMP2, or truncation of the last nine residues of SNAP25, retard fusion pore expansion in adrenal chromaffin cells (20, 21, 25). Mutations in SNARE TMDs also affect fusion pores (18). Increasing the number of SNAREs at the fusion site accelerated fusion pore expansion in neurons (15, 26), astrocytes (27) and chromaffin cells (28) and led to larger pores in nanodisc-based single-pore fusion assays (15, 17). This was interpreted as due to increased molecular crowding at the waist of the pore with increasing SNARE copy numbers (17).

Although they are best known for their role as calcium sensors for exocytosis at most synapses and endocrine cells, Synaptotagmins are another component of the release machinery known to affect fusion pore properties (29–39). They couple membrane fusion driven by neuronal/exocytic SNAREs to calcium influx (40, 41). Synaptotagmins are integral membrane proteins possessing two cytosolic C2 domains (C2A and C2B) which can bind Ca^2+^, acidic lipids, SNAREs, and other effectors, but affinities vary widely among the 17 mammalian isoforms (41–48). Synaptotagmin-1 (Syt1) is the major neuronal isoform that mediates fast, synchronous neurotransmitter release (41, 45, 49). It resides in synaptic vesicles in neurons and secretory granules in neuroendocrine cells and interacts with SNAREs, acidic phospholipids, and calcium (1, 41, 50, 51). How calcium binding to Syt1 leads to the opening of a fusion pore is an area of active research and debate (37, 50, 52–60). In addition to its role in triggering the opening of a fusion pore, Syt1 also affects the expansion of the fusion pore after it has formed (29–39), but mechanisms are even less clear.

Calcium-binding to Syt1 causes hydrophobic residues at the tips of the Ca^2+^-binding loops to insert into the membrane, generating curvature, which may be important for triggering fusion (37, 52, 53). Membrane bending has been proposed to facilitate opening of the initial fusion pore by helping to bring the two membranes into close proximity, reducing the repulsive hydration forces by reducing the contact area, and exposing the hydrophobic interior of the two membranes to initiate lipid exchange (61, 62). After fusion pore opening, Syt1 was suggested to contribute to fusion pore expansion through membrane curvature generation as well, based on the observation that in PC12 cells, membrane-insertion deficient mutants reduced exocytosis whereas mutants with enhanced insertion led to larger fusion pores (37). However, once the initial fusion pore is formed it is not clear whether and how much curvature generation by Syt1contributes to fusion pore expansion. First, in PC12 cells multiple Syt isoforms reside on the same secretory granule and potentially compete for fusion activity (63, 64). Disrupting Syt1 function may allow another isoform to dominate fusion pore dynamics. In adrenal chromaffin cells where Syt1 and Syt7 are sorted to distinct granule populations, fusion pores of Syt7 granules dilate more slowly (38). Second, compared to Syt1 C2 domains, the higher calcium-affinity Syt7 C2 domains penetrate more avidly and deeply into membranes (65, 66), which should lead to more efficient membrane bending (52, 53). This would appear to be inconsistent with the slower dilation of fusion pores by Syt7. Finally, most previous reconstitutions could not probe the role of Syt1 in fusion pore regulation, as they lacked the required sensitivity and time resolution to detect single pores.

Here we investigated the mechanism by which Syt1 contributes to fusion pore dynamics, using a single-pore conductance assay (16, 17). Compared to SNAREs alone, addition of Syt1 increased the mean pore conductance three-fold. This effect required binding of Syt1 to calcium, the acidic phospholipid PI(4,5)P_2_, and likely to the SNAREs. In addition, both pore opening and dilation are promoted by insertion of Syt1 C2AB top loops into the membrane in a Ca^2+^-dependent manner, but we propose that membrane curvature generation is not needed to explain fusion pore expansion by Syt1. Mathematical modeling suggests that pore dilation relies on regulation of the intermembrane distance by Syt1. Syt1 penetration into the target membrane upon calcium binding re-orients the C2AB domains and SNARE complexes, forcing the membranes apart in a lever-like action that concomitantly expands the pore.

## RESULTS

### Co-reconstitution of Synaptotagmin-1 and v-SNAREs into nanolipoprotein particles

Previously, using a nanodisc-cell fusion assay, we characterized single, SNARE-induced fusion pores connecting a nanodisc and an engineered cell expressing neuronal “flipped” t-SNAREs ectopically (16, 17). In this assay, a flipped t-SNARE cell is voltage-clamped in the cell-attached configuration. Nanodiscs reconstituted with the neuronal/exocytotic v-SNARE VAMP2 are included in the pipette solution. Fusion of a nanodisc with the cell surface creates a nanometer size pore that connects the cytosol to the exterior, allowing passage of ions under voltage clamp. Direct currents report pore size with sub-millisecond time resolution (16, 17). Fusion pore currents fluctuate and may return to baseline transiently multiple times, evidently reflecting pore flickering (2, 16, 17, 67). Pore conductance is eventually lost (5-20 s on average after initial appearance), evidently reflecting pore closure (2, 16, 17, 67). The mechanism of pore closure is not known, but because pore expansion beyond a maximum size is prevented by the nanodisc scaffold, pore closure is one of the few possible outcomes (16, 68). To ensure single-pore detection, the rate at which pore currents appear (reported in pores/min, also referred to as the “fusion rate”) is made low by recording from a small area of the cell surface and by tuning the nanodisc concentration (2, 16, 17, 67) (See Materials and Methods and SI Appendix for details).

To test whether Syt1 affected fusion pores in this system, we co-reconstituted ~4 copies of recombinant full-length Syt1 together with ~4 copies of VAMP2 (per disc face) into large nanodiscs called nanolipoprotein particles (17, 69) (vsNLPs, ~25 nm in diameter, see SI Appendix, fig. S1). We reasoned that, under these conditions, potential modification of pore properties by Syt1 should be detectable. In the absence of Syt1, we previously found that only ~2 SNARE complexes are sufficient to open a small fusion pore (150-200 pS conductance), but dilation of the pore beyond ~1 nS conductance (~1.7 nm in radius, assuming the pore is a 15 nm long cylinder (70)) required the cooperative action of more than ~10 SNARE complexes (17). The increase in pore size was heterogeneous with increasing SNARE load; most pores remained small (mean conductance ≲ 1 nS), but an increasing fraction had much larger conductances of a few nS. With ~4 v-SNAREs per NLP face, fusion driven by SNAREs alone results in relatively small pores with ~200 pS average conductance, corresponding to a pore radius of ~0.76 nm (17). Larger pores (mean conductance>1 nS) were rare (<5%, (17)). With ~25 nm NLPs, a fusion pore can in principle grow to >10 nm diameter (~9 nS conductance) before the scaffold protein stabilizing the edges of the NLP becomes a limitation for further pore dilation (17). Thus, at this v-SNARE density, there is a large latitude in pore sizes that can be accommodated, if introduction of Syt1 were to lead to any modification.

We tuned NLP size by varying the lipid-to-scaffold protein (ApoE422k) ratio and adjusted copy numbers of VAMP2 and Syt1 until we obtained the target value of ~4 copies of each per NLP face, similar to previous work with SNAREs alone (17, 69). vsNLPs were purified by size exclusion chromatography and characterized by SDS-PAGE and transmission electron microscopy (see SI Appendix, fig. S1B-D). The distribution of NLP diameters was fairly narrow, with mean diameter 25 nm (±5.6 nm SD, see SI Appendix, fig. S1E), and did not change significantly compared to the distribution when v-SNAREs alone were incorporated at ~4 copies per face (mean diameter = 25 ± 4 nm, (see SI Appendix, fig. S1F-H, and (17)).

### Syt1 promotes fusion pore expansion

To probe fusion pores, we voltage-clamped a flipped t-SNARE cell in the cell-attached configuration and included NLPs co-loaded with Syt1 and VAMP2 in the pipette solution as shown in Fig. 1 (100 nM vsNLPs, 120 μM lipid). Even in the presence of 100 μM free calcium, a level that elicits robust release in neurons and chromaffin cells (44, 71–74), pore properties were affected only slightly compared to the case when Syt1 was omitted from the NLPs. For example, pore currents (Fig. 1D) appeared at similar frequency (Fig. 2A) and the mean single-pore conductance, 〈*G*_*po*_〉, was only slightly elevated in the presence of Syt1 (See SI Appendix, fig. S1J, S2, and Supplementary Materials and Methods for definitions and other pore parameters). We wondered whether the lack of acidic lipids in the outer leaflet of the cell membrane could be a limitation for Syt1’s ability to modulate fusion pores. Syt1 is known to interact with acidic lipids, in particular with PI(4,5)P_2_, in both calcium-dependent and independent manners, and these interactions are required for Syt1’s ability to trigger membrane fusion (30, 55, 75–78). However, the outer leaflet of the plasma membrane which is seen by Syt1 in our assay is poor in such lipids. To test for a requirement for PI(4,5)P_2_, we incubated flipped t-SNARE cells with 20 μM diC_8_PI(4,5)P_2_ for 20 min and rinsed excess exogenous lipid. At different time points after rinsing, we probed incorporation of the short-chain PI(4,5)P_2_ into the outer leaflet of the cell membrane by immunofluorescence, using a mouse monoclonal anti-PI(4,5)P_2_ primary antibody, without permeabilizing the cells (Fig 1B). The signal decreased slightly as a function of time but persisted for at least 80 min. To compare the level of short-chain PI(4,5)P_2_ incorporated into the outer leaflet in this manner with endogenous PI(4,5)P_2_ levels in the inner leaflet, we measured immunofluorescence from permeabilized cells that were not incubated with diC_8_PI(4,5)P_2_. Outer leaflet diC_8_PI(4,5)P_2_ levels were within 25% of the endogenous inner-leaflet PI(4,5)P_2_ levels (Fig. 1B).

**Figure 1.**
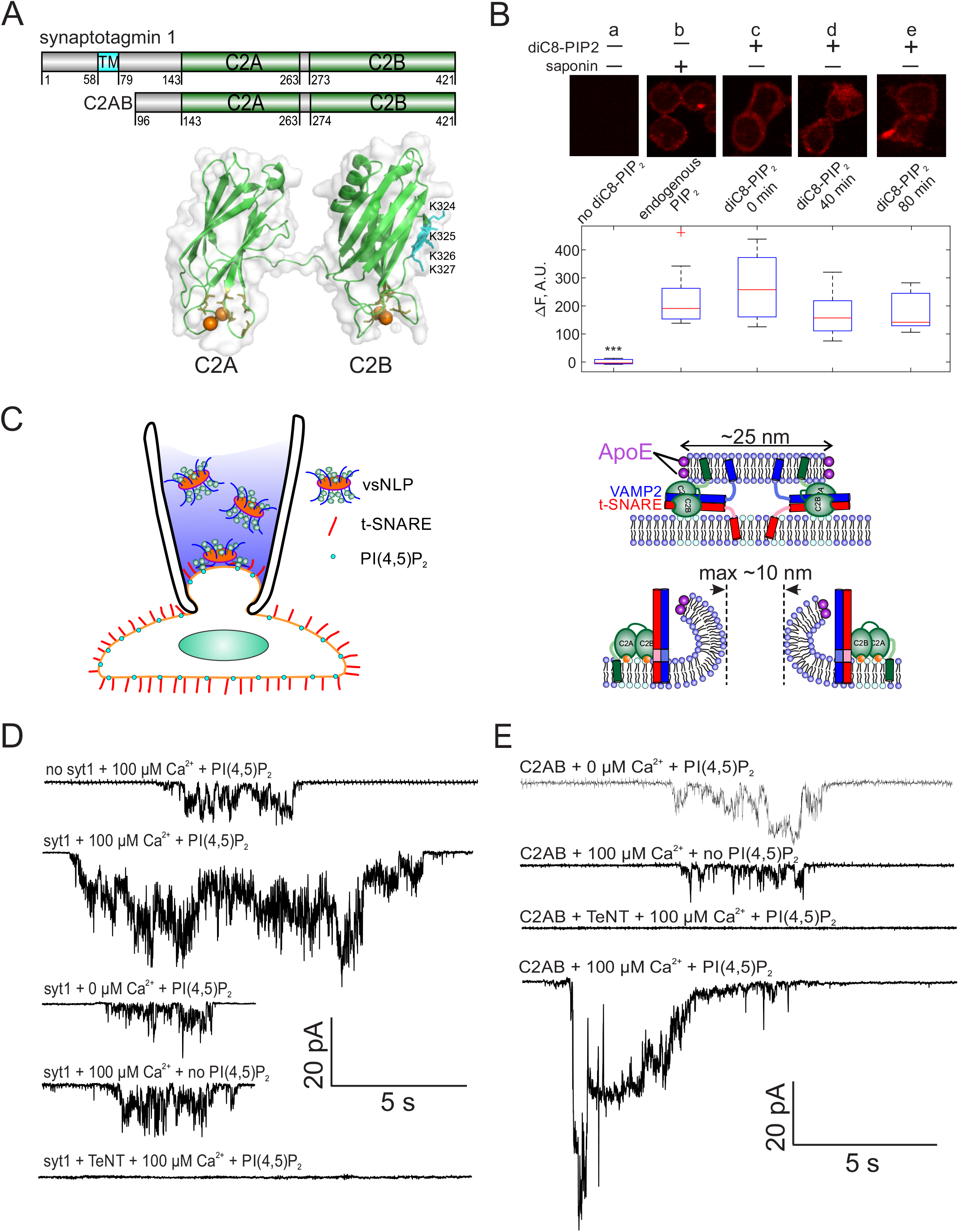
Detection of single fusion pore currents mediated by Syt1 or its C2AB domain. **A**. Domain structures of the constructs used in this study. The structure of the soluble C2AB domains was rendered using PyMol, from PDB: 5kj7 (144). The orientations of the C2A and C2B domains relative to each other are not known in the presence of SNAREs and membranes. Conserved aspartate residues coordinating calcium ions are depicted in orange. Calcium ions are shown as orange spheres. A poly-lysine motif on the side of C2B (K324,K325,K326,K327 in the rat sequence) that preferentially interacts with PI(4,5)P_2_ (91) is highlighted in cyan. **B**. Incorporation of exogenous PI(4,5)P_2_ into the outer leaflet of flipped t-SNARE cells. Top: cells were incubated with diC8-PI(4,5)P_2_ for 20 min, rinsed, and immunolabeled for PI(4,5)P_2_ at the indicated time points. Only control cells that were permeabilized with saponin showed immunostaining, confirming absence of PI(4,5)P_2_ in the outer leaflet, and providing a reference value for inner-leaflet PI(4,5)P_2_ levels (a and b). Cells incubated with diC8-PI(4,5)P_2_ showed immunofluorescence in the absence of permeabilization, indicating successful incorporation of PI(4,5)P_2_ into the outer leaflet of the cell membrane (c-e). The signal was comparable to endogenous inner-leaflet PI(4,5)P_2_ levels, and persisted at least for 80 min (lower panel). Cells processed similarly, but not treated with saponin or diC8-PI(4,5)P_2_ served as negative controls (a). One-way analysis of variance (ANOVA) followed by multiple comparison test was used to compare the signals from the endogenous PI(4,5)P_2_ sample (b) with all others. *, **, *** indicate p<0.05, 0.01, and 0.001, respectively. **C**. Schematic of the single-pore nanodisc-cell fusion assay. A glass micropipette forms a tight seal on a patch of the plasma membrane of a cell expressing “flipped” t-SNARE proteins on its surface. NLPs co-reconstituted with Syt1 and VAMP2 are included in the pipette solution (left). NLP-cell fusion results in a fusion pore connecting the cytosol to the cell’s exterior (right). Under voltage clamp, direct-currents passing through the pore report pore dynamics. With ~25 nm NLPs, the scaffolding ring does not hinder pore expansion up to at least 10 nm diameter. Exogenous PI(4,5)P_2_ can be added to the cell’s outer leaflet as in B, and calcium in the pipette is controlled using calcium buffers. **D**. Representative currents that were recorded during vsNLP-tCell fusion, for the indicated conditions. PI(4,5)P_2_ indicates cells were pre-treated with diC8-PI(4,5)P_2_. Tetanus neurotoxin (TeNT) light chain cleaves VAMP2 and blocks exocytosis. Currents were larger when all components were present (SNAREs, Syt1, exogenous PI(4,5)P_2_ and calcium). **E**. Similar to D, but instead of full-length Syt1, 10 μM soluble Syt1 C2AB domains were used together with NLPs carrying ~4 copies of VAMP2 per face.

**Figure 2.**
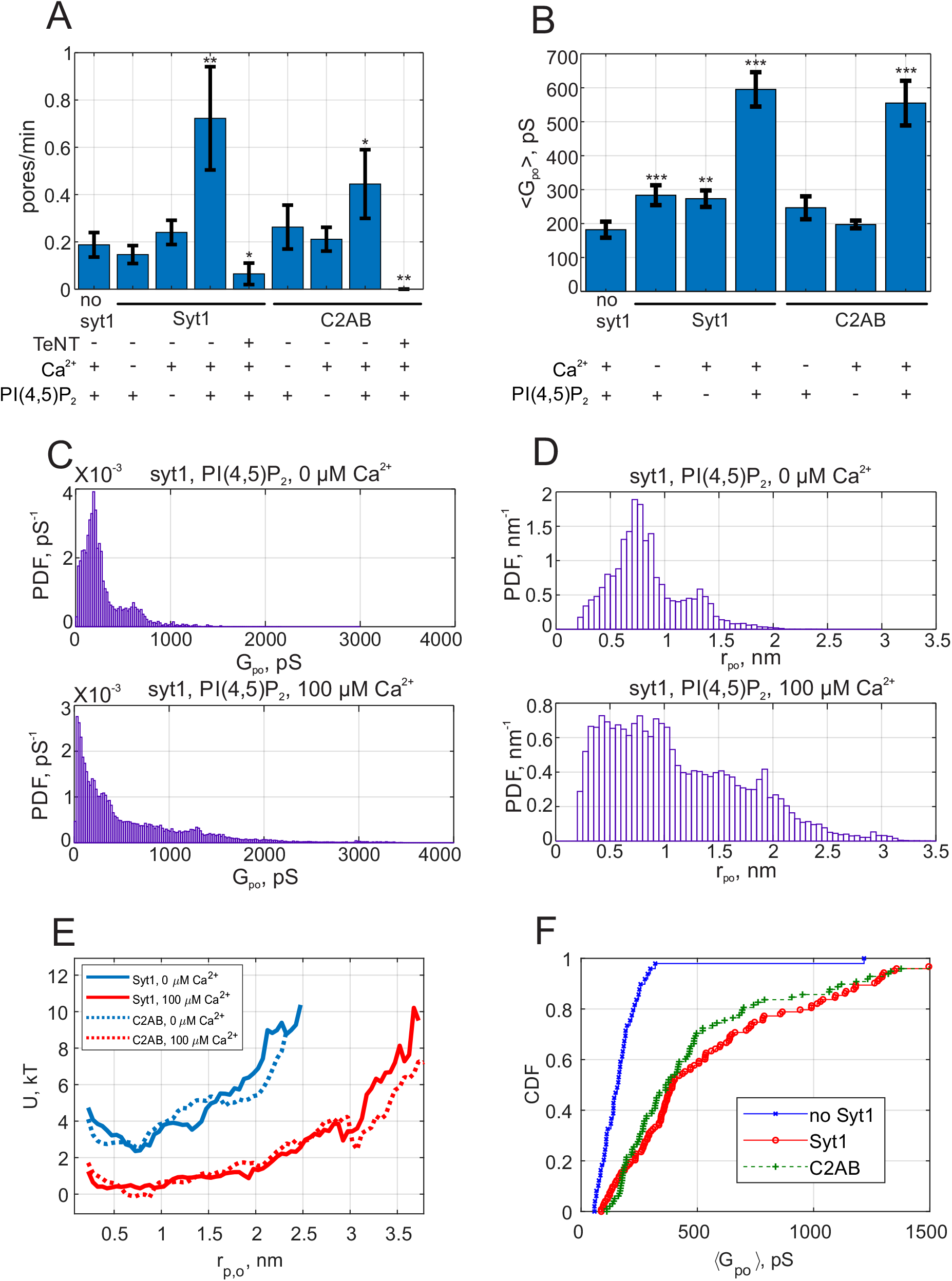
Syt1 promotes fusion and expands fusion pores in a calcium and PI(4,5)P_2_ dependent manner, and soluble Syt1 C2AB recapitulates these effects. **A**. The rate at which current bursts appeared (pore nucleation rate) for the conditions indicated (error bars represent ± S.E.M.). SNARE-induced pores appeared more frequently in the presence of Syt1 or C2AB, when both calcium and PI(4,5)P_2_ were also present. Student’s t-test was used to assess significant differences between the “no Syt1” group and the rest. *, **, *** indicate p<0.05, 0.01, and 0.001, respectively. There is no difference between the Syt1 and C2AB groups in the presence of calcium and exogenous PI(4,5)P_2_ (Student’s t-test: *p* = 0.18). **B**. Mean single fusion pore conductance, 〈*G*_*po*_〉, for different conditions as indicated (± S.E.M.). 〈*G*_*po*_〉 was three-fold larger in the presence of Syt1 or C2AB, when both calcium and PI(4,5)P_2_ were also present. Two-sample Kolmogorov-Smirnov test was used to assess significant differences between the “no Syt1” group and the rest. The same asterisk notation as in A was used. There is no difference between the Syt1 and C2AB groups in the presence of calcium and exogenous PI(4,5)P_2_ (two-sample Kolmogorov-Smirnov test: *p* = 0.29). **C**. Probability density functions (PDFs) for point-by-point open-pore conductances (see Methods) for pores induced in the presence of Syt1, PI(4,5)P_2_ and with 0 or 100 μM calcium. Notice the higher density at larger conductance values in the presence of 100 μM calcium. **D**. Probability density functions for pore radii, calculated from the conductance PDFs in C, assuming a 15-nm long cylindrical pore (70). **E**. Apparent free energy profiles for Syt1 and soluble Syt1 C2AB domains in the absence or presence of calcium. These profiles were calculated from the pore radii PDFs as in D (see text and Methods) (17). The profiles were shifted vertically for clarity. **F**. Cumulative density functions (CDFs) for mean single-pore conductances for the conditions indicated. Soluble C2AB recapitulated effects of full-length Syt1 co-reconstituted into NLPs.

When we repeated vsNLP-flipped t-SNARE cell fusion experiments with cells pre-incubated with diC_8_PI(4,5)P_2_, the rate of fusion in the absence of calcium was unchanged compared to fusion with SNAREs alone, but increased 3-4 fold when 100 μM calcium was present (Fig. 2A). Compared to SNARE-alone fusion, the mean single-pore conductance increased only slightly in the absence of calcium but was three-fold larger in the presence of 100 μM calcium (Fig. 2B). Conductance fluctuations around the mean value were larger and flicker frequency lower when Syt1, calcium and PI(4,5)P_2_ were all present, but no major differences emerged for burst lifetimes, *T*_*o*_, or pore open probability during a burst (the fraction of time the pore was open during a burst), *P*_*o*_ (see SI Appendix, fig. S3). For all cases tested, the distributions of the number of pore flickers (*N*_*flickers*_) and burst durations (*T*_*o*_) were well-described by geometric and exponential distributions, respectively (see SI Appendix, fig. S3), as would be expected for discrete transitions between open, transiently blocked, and closed states (79). Fusion was SNARE-dependent, as treatment with the tetanus neurotoxin TeNT, which cleaves VAMP2 at position 76Q-77F and blocks exocytosis (80), dramatically reduced the fusion rate of vsNLPs even in the presence of calcium and exogenous PI(4,5)P_2_ (Fig 1D and Fig 2A). Thus, Syt1 increases the fusion rate and promotes pore dilation during SNARE-induced fusion, in a calcium- and PI(4,5)P_2_-dependent manner.

We pooled individual current bursts to obtain the distributions for fusion pore conductances and pore radii as shown in Fig. 2C, D, and Figure S4. The distributions were similar for SNAREs alone, whether calcium or PI(4,5)P_2_ were added, and with Syt1 when calcium was omitted (Fig. 2C, D, and see SI Appendix, fig. S4). By contrast, in the presence of 100 μM free calcium and exogenous PI(4,5)P_2_, larger conductance values (and corresponding pore radii) became more likely (Fig. 2C,D).

Even when pores were maximally dilated by Syt1, the mean conductance and pore radius, *G*_*po*_ = 595 pS (S.E.M.= 51 pS), and *r*_*po*_ = 1.13 nm (S.E.M= 0.04 nm) were significantly less than the maximum possible value predicted from NLP dimensions (17). That is, the geometric constraints imposed by the NLP dimensions were not limiting pore expansion. Instead, there is inherent resistance to pore dilation, independent of NLP scaffolding (17) as predicted and observed in other systems (81–83). To quantify the resistance, we computed the apparent pore free energy *U*(*r*_*po*_) from the distribution of pore radii, 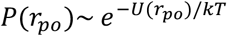 for fusion with both SNAREs alone and with Syt1 under optimal conditions (with exogenous PI(4,5)P_2_ and 100 μM free calcium). With SNAREs alone, or with Syt1 but in the absence of calcium, the free energy profile suggested that ~6-7 kT energy was required to expand the pore from 1 to ~2.5 nm radius, whereas calcium-bound Syt1 reduced this resistance to ~2 kT (Fig. 2E). That is, the force opposing pore expansion decreased from 16-19 pN in the absence of calcium to ~5 pN in the presence of 100 μM calcium.

We tested if the soluble C2AB domains of Syt1 could recapitulate these results. We included 10 μM C2AB together with NLPs reconstituted with ~4 copies per face of VAMP2 in the patch pipette and monitored fusion with flipped t-SNARE cells in the cell attached configuration under voltage clamp. Similar to the results with full-length Syt1, there was little change in fusion rate and mean conductance compared to the SNARE-alone case, unless calcium and exogenous PI(4,5)P_2_ were also present. In the presence of 100 μM calcium and short-chain PI(4,5)P_2_, fusion rate and mean pore conductance were stimulated to the same degree as when full-length Syt1 was used (Fig. 2). The distributions of average single pore conductances (Fig. 2F), conductance fluctuations, and other pore parameters were similar whether full-length Syt1 or soluble C2AB were used, except *P*_0_ was higher for the +Ca^2+^/+PI(4,5)P_2_ case and *T*_1_ lower for +Ca^2+^/-PI(4,5)P_2_ case for C2AB compared to Syt1 (Figs. S3 and S4). The apparent free energy profile calculated from the pore size distribution was indistinguishable from that of full-length Syt1 (Fig. 2E). We conclude that soluble Syt1 C2AB recapitulates the effect of full-length Syt1 on promoting dilation of SNARE-mediated fusion pores. As they were far easier to manipulate, we used soluble Syt1 C2AB domains for the remainder of this work.

In some cases, a peak at ~200 pS is apparent in open-pore conductance distributions, corresponding to a peak at *r*_*po*_ ≈ 0.7 nm in pore size distributions (e.g. see see SI Appendix, fig. S4). This is manifested as a small dip in the energy profiles (Fig. 2E). We do not know the underlying mechanisms, as we have not identified a clear correlation between the peak’s amplitude or location and the parameters we varied, such as calcium concentration.

### Pore dilation by Synaptotagmin-1 C2AB requires binding to calcium, PI(4,5)P_2_, and likely SNAREs

We further tested the requirement for Syt1 C2AB binding to calcium, PI(4,5)P_2_, and SNAREs for promoting pore dilation, using mutagenesis (Fig. 3). Binding of calcium to the second C2 domain of Syt1 is known to be essential for evoked release (41, 84, 85). When calcium binding to the C2B domain was impaired by mutating a highly conserved aspartate to asparagine (Syt1 C2AB D309N (86)), mean single pore conductance returned to the value obtained in the presence of SNAREs alone (Fig. 3C). The rate at which current bursts appeared also returned to the SNARE-alone level (Fig. 3B). Other pore properties were also indistinguishable from the SNARE-alone case (see SI Appendix, fig. S5). We conclude that calcium binding to Syt1 C2B is essential for fusion pore dilation, in addition to its well-known role for triggering the opening of a fusion pore (32).

**Figure 3.**
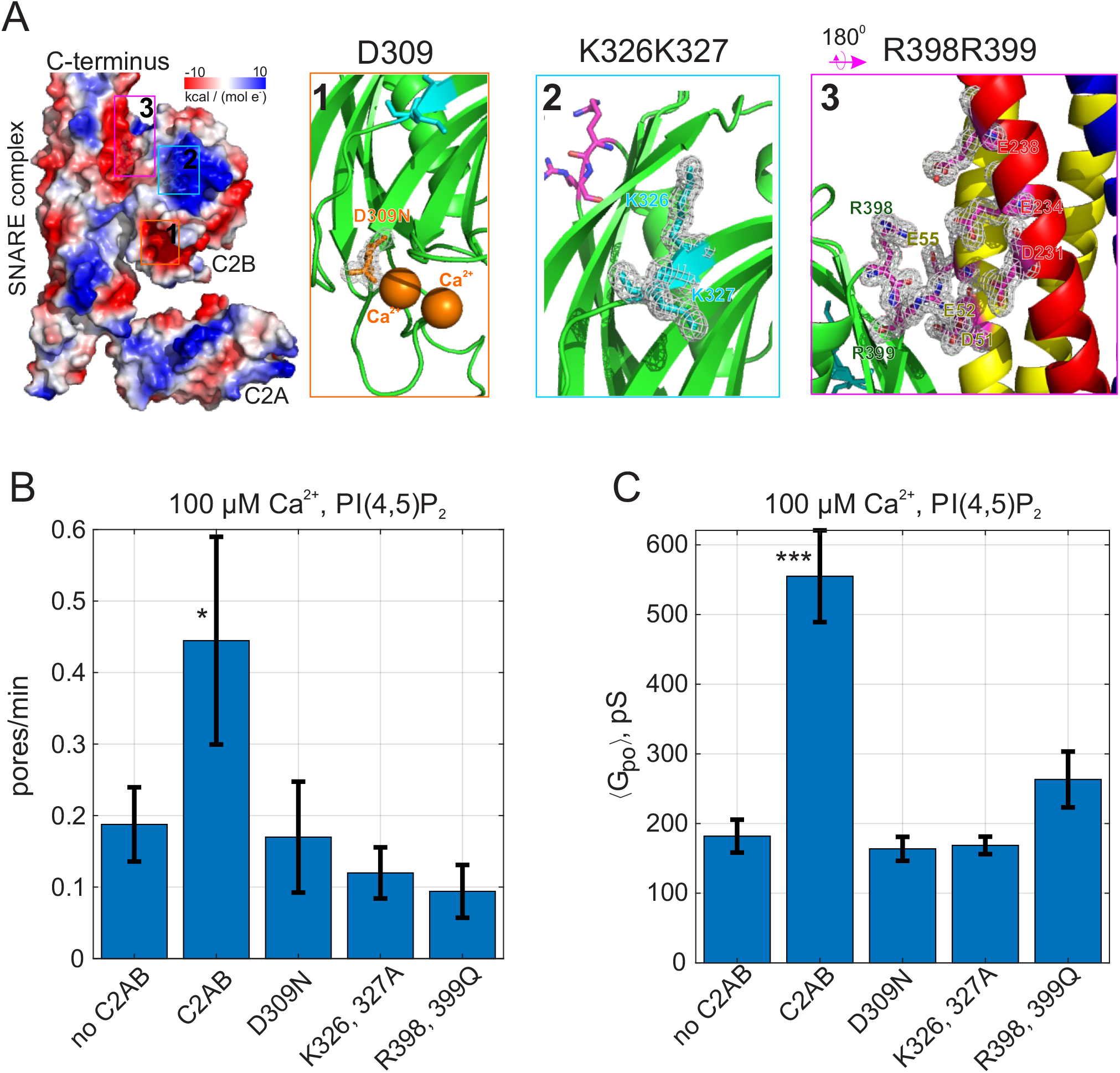
Pore expansion by Syt1 C2AB requires calcium, PI(4,5)P_2_, and putative SNARE binding sites to be intact. **A**. Overview of the Syt1-SNARE complex (144). The electrostatic potential of PDB 5kj7 (144) was rendered using Pymol. The sites mutated in this work are marked by boxes labeled 1-3 on the left and shown in the panels to the right. D309 is a key calcium-binding residue (1), K326, K327 interact with acidic lipids (2), and R398,R399 interact with the t-SNAREs SNAP 25 (E51, E52 and E55) and syntaxin 1A (D231, E234 and E238). VAMP2 is shown in blue, SNAP25 in yellow, and syntaxin 1A in red. **B**. Pore nucleation rates (+/− SEM) for the indicated conditions. All conditions included 100 μM free calcium and pre-incubation of tCells with exogenous PI(4,5)P_2_. Pores appeared 2-3 times less frequently with the mutated proteins compared to wild-type Syt1 C2AB. Student’s t-test was used to assess significant differences between the “no C2AB” group and the rest. **C**. Mean single open-pore conductance values (+/− SEM) for the same conditions as in B. Disrupting binding to calcium (D309N), acidic lipids (K326A, K327A), or the SNARE complex (R398, R399) resulted in ~3-fold smaller mean conductance compared to wild-type C2AB, abrogating the effects of Syt1 C2AB. Two-sample Kolmogorov-Smirnov test was used to assess significant differences between the “no C2AB” group and the rest. *, **, *** indicate p<0.05, 0.01, and 0.001, respectively.

The C2B domain of Syt1 possesses a polybasic patch (K324-327) that interacts with acidic phospholipids (Fig. 3A) and is important for synchronous evoked release (55). Although this interaction occurs in the absence of calcium (41), it contributes to the membrane binding energy of C2AB in the presence of calcium (75), presumably because multivalent interactions increase the bound lifetime of C2AB. Partially neutralizing the polybasic patch in C2B (K326A, K327A) reduced the fusion rate, and resulted in single pore conductances that were indistinguishable from those for SNARE-alone pores (Fig. 3). Similarly, the burst lifetime and the flicker rate were comparable to the SNARE-alone level, but conductance fluctuations were reduced, while there was an increase in the pore open probability during a burst, *P*_*o*_ (see SI Appendix, fig. S5), as would be expected for pores that fluctuate less. Thus, in addition to its established role in evoked release (55, 87), the polybasic patch in Syt1 C2B is also required for fusion pore dilation.

Two recent crystal structures identified a “primary” interaction interface between Syt1 C2B and the four-helical SNARE complex (88, 89) (Fig.3A). Specifically, two arginines (R398 and R399) form salt bridges with glutamates and aspartates in a groove between SNAP25 and Syntaxin-1 (88). Mutation of these arginines to glutamines (R398Q, R399Q) was shown to largely abolish evoked release from hippocampal neurons (55, 88, 90), possibly by disrupting the interaction of Syt1 C2B with SNAREs (55, 88). When we used purified C2AB bearing the same mutations (C2AB^R398Q, R399Q^) both the fusion rate and the mean pore conductance decreased significantly, close to SNARE-alone levels (Fig. 3B, C). Burst lifetimes, conductance fluctuations, and the pore open probability were not significantly different than for pores induced by SNAREs alone, but the flicker rate was lower (see SI Appendix, fig. S5).

Together, these results indicate that binding of Syt1 to calcium, PI(4,5)P_2_, and likely SNAREs, which are all crucial for Syt1’s role in evoked neurotransmitter release (1, 41), are also essential for its function in dilating SNARE-induced fusion pores.

### Calcium-dependence of pore dilation by Syt1 C2AB

To determine whether pore properties are altered by calcium, we varied the free calcium concentration in the pipette solution and repeated the fusion experiments. Mean open-pore conductance 〈*G*_*po*_〉 increased with increasing calcium (Fig. 4A), consistent with a mathematical model (see below). Conductance fluctuations and burst lifetimes also increased, while the flicker rate decreased slightly and the pore open probability during a burst did not change significantly as [Ca^2+^] was increased (see SI Appendix, fig. S6). That is, pores tended to last longer with higher calcium, and the open state conductance increased. The rate at which pore currents appeared also increased with calcium (SI Appendix, fig. S6F).

**Figure 4.**
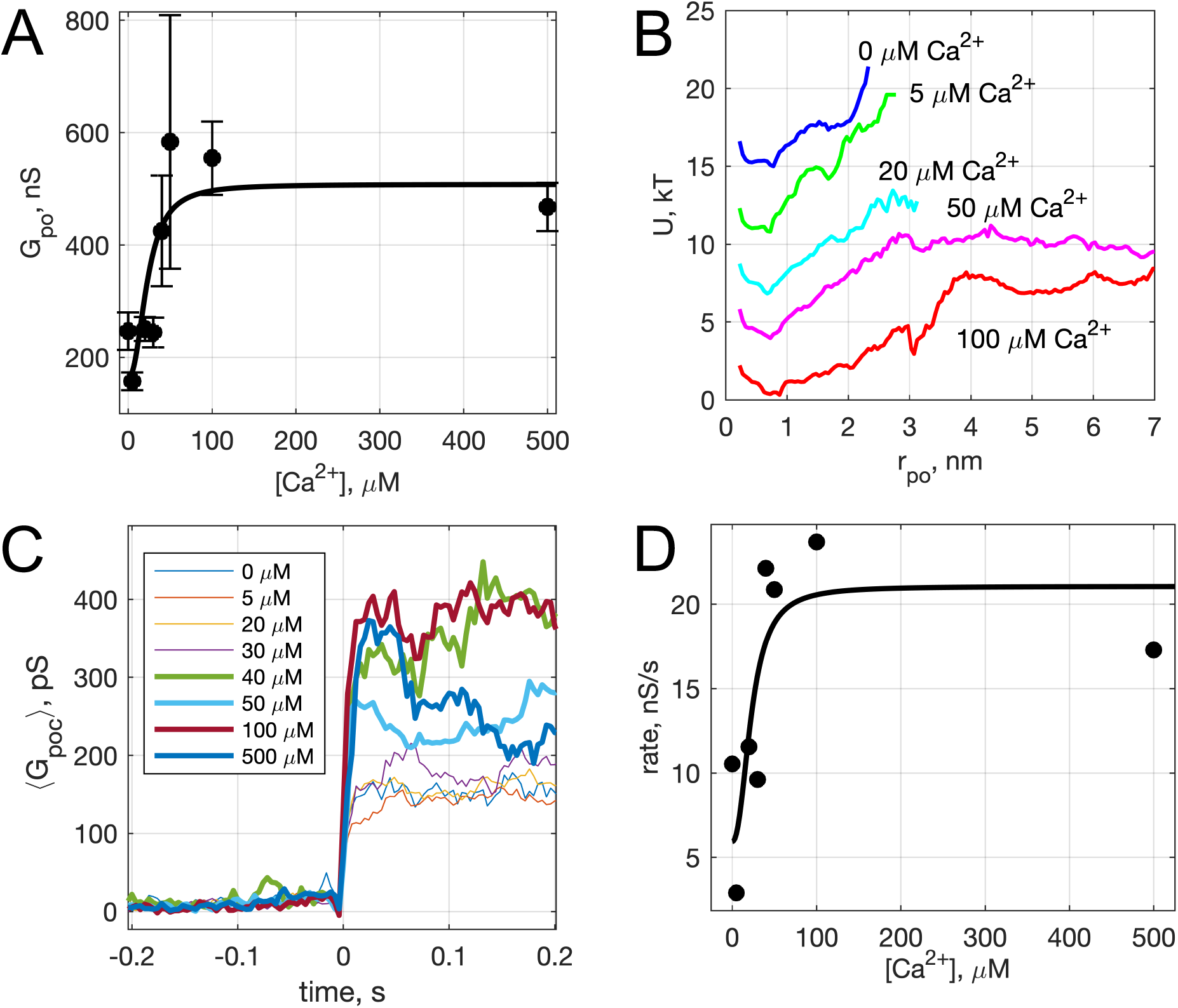
Calcium-dependence of pore properties. **A**. Mean single open-pore conductance, 〈*G*_*po*_〉, as a function of free calcium concentration in the pipette solution. Plotted values are mean± S.E.M. A fit to a Hill equation 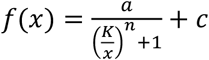 is shown as the black line, where *x* = [Ca^2+^]_*free*_, *n* = 2.3, and *K* = 23 μM (see text). Best fit parameters (with 95% confidence bounds) were *a* = 343.7 (128.5, 558.8), *c* = 164.2 (14.5, 314), and *R*^2^ = 0.72. **B**. Apparent free energy profiles, calculated as in Fig. 2E, for different calcium concentrations. **C.** Kinetics of pore expansion for different [Ca^2+^]_free_ as indicated. Conductance traces were aligned to the first point in a pore and averaged. **D**. Expansion rates of time-aligned and averaged conductances as a function of [Ca^2+^]_free_. Expansion rates were calculated as the 10-90% rise time from the baseline to the level of conductance reached within the first 100 ms after pore opening, divided by the time it took for this rise (see SI Appendix, Supplementary Materials and Methods). A fit to a Hill equation as in A is also shown, using the same *x* and *n* parameter values.

The conductances in the open-state and the corresponding pore radii (*r*_*po*_) were broadly distributed at all calcium concentrations tested, but the distributions did not shift uniformly as calcium increased (see SI Appendix, fig. S6). The apparent free energy profiles, estimated from the pore size distributions, are plotted in Fig. 4B. With increasing calcium, the well around the most likely radius (~0.5-0.7 nm) became wider, and the slopes of the energy profiles for radii above the well’s upper boundary, reflecting the force needed to dilate the pore, decreased as calcium increased. The calcium concentration at which this transition occurs (~20 μM) is consistent with the known calcium binding affinity of Syt1 (75, 76, 78, 91, 92).

We also examined the kinetics of pore dilation as a function of calcium (Fig. 4C,D). To this end, we averaged pore conductances after aligning them to the initial pore opening, in the presence of Syt1 C2AB at different Ca^2+^ levels. The average conductance rapidly increased after initial pore opening for all traces, but reached larger values for larger calcium concentrations (Fig. 4C). We estimated the pore expansion rate as the 10-90% rise time from the baseline to the level of conductance reached within the first 100 ms after pore opening, divided by the time it took for this rise (Fig. 4D). With low amounts of calcium (0-30 μM), the expansion rate is ~3-12 nS/s, which increases rapidly to 20-25 nS/s for 40-100 μM Ca^2+^.

Both the increase in mean open-pore conductance (Fig. 4A) and the pore expansion rate (Fig. 4D) with increasing free calcium were fit to a Hill equation, using parameters describing cooperative binding and loop-insertion of Syt1 C2AB to lipid bilayers containing PI(4,5)P_2_ (78).

### Calcium-dependent membrane-insertion of Syt1 C2AB is necessary for pore dilation

Calcium binds simultaneously to acidic phospholipids and highly conserved aspartate residues in a pocket formed by loops at the top of the beta-sandwich structure of the Syt1 C2 domains (41, 85, 93). Hydrophobic residues at the tips of the loops flanking the calcium-binding residues in Syt1 C2A (M173 and F234) and C2B (V304 and I367) insert into the membrane as a result of these interactions, strengthening membrane binding of C2 domains (41, 75, 94) while causing a reorientation of the C2 domains (95, 96) (Fig. 5A). The membrane insertion of these hydrophobic residues contributes to the triggering of release (37, 52, 53). We wondered whether membrane-insertion of hydrophobic loops also played any role in pore dilation. To test this, we introduced mutations that made the loops insertion-deficient (M173A, F234A, V304A and I367A, the “4A” mutant (37, 52)) or that increased membrane affinity (M173W, F234W, V304W and I367W, the “4W” mutant (37, 52)).

**Figure 5.**
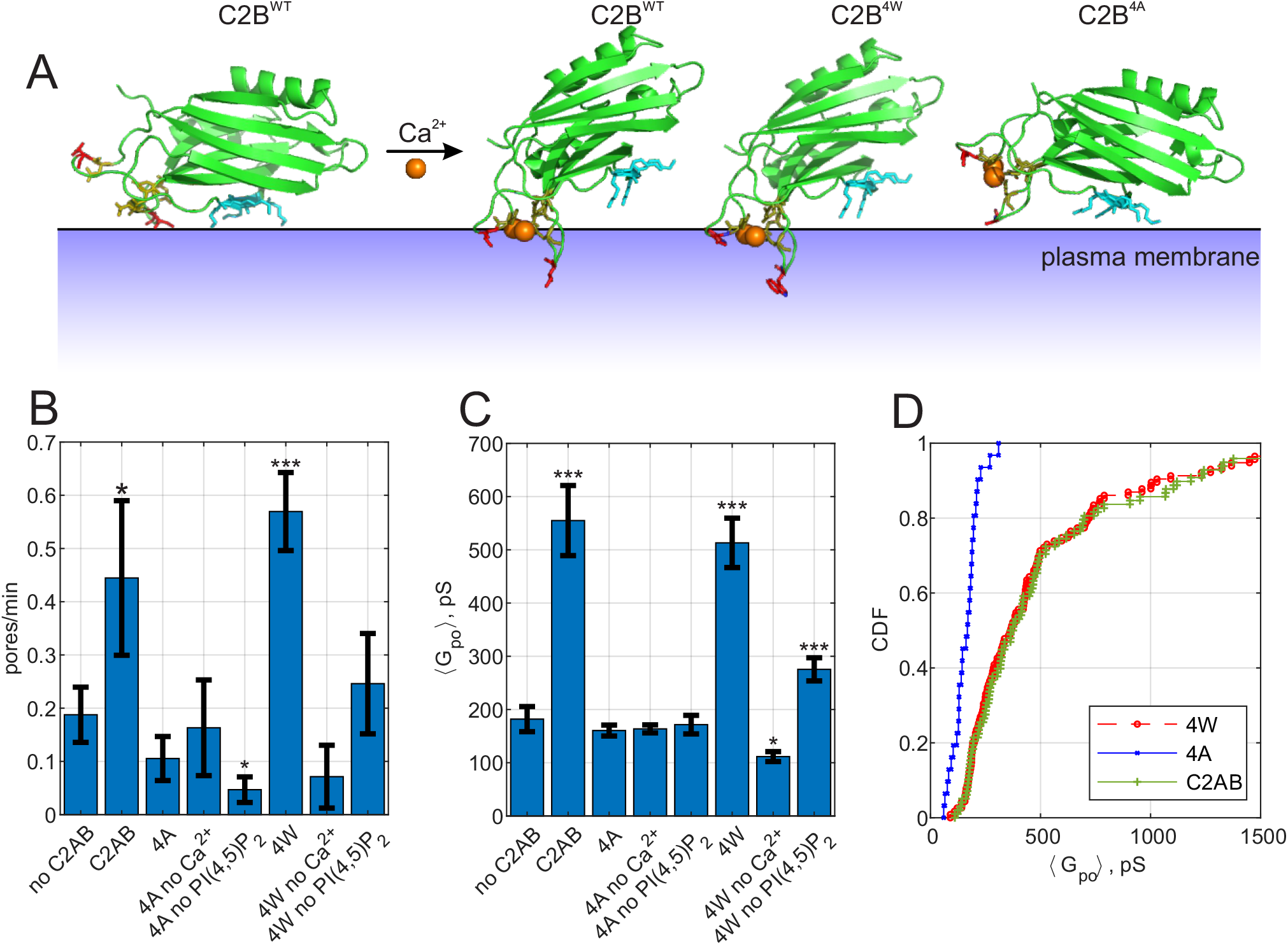
Calcium-induced membrane insertion of Syt1 C2AB hydrophobic loops are critical for both pore nucleation and expansion. **A**. Schematic depiction of Syt1 C2B domain’s calcium-dependent interactions with membranes. Calcium-free C2B interacts with acidic lipids through its poly-lysine motif (highlighted in cyan as in Fig. 1). Upon binding to calcium, hydrophobic residues (V304 and I367 on C2B) insert into the membrane, causing C2B to reorient (41) and inducing membrane curvature (52, 53). In the presence of PI(4,5)P_2_, the calcium-bound C2B assumes a tilted conformation with respect to the membrane (95). M173 and F234 on C2A top loops similarly insert into membranes in a calcium-dependent manner, with similar effect on orientation and curvature generation (41) (not shown). A mutant with the membrane-inserting residues replaced with tryptophans (M173W, F234W, V304W and I367W, “4W”) binds membranes more avidly, resulting in more membrane tubulation activity, whereas alanine substitution of the same residues (“4A”) abolishes membrane penetration and curvature induction (52). **B.** Pore nucleation rate (mean ± S.E.M) in the presence of wildtype, 4W and 4A mutants. Student’s t-test was used to assess significant differences between the “no C2AB” group and the rest. **C**. Mean open-pore conductance (± S.E.M) for the conditions indicated. Two-sample Kolmogorov-Smirnov test was used to assess significant differences between the “no C2AB” group and the rest. **D**. Cumulative density functions for mean open-pore conductances for wild-type Syt1 C2AB, 4W and 4A mutants. In A, calcium-free C2B was rendered from PDB 5w5d (89) and calcium-bound C2B was rendered from 5kj7 (144). *, **, *** indicate p<0.05, 0.01, and 0.001, respectively.

In the nanodisc-cell fusion assay, the membrane penetration deficient 4A mutant was non-functional, having no discernible effect on pore dilation or fusion rate when compared to the assay without Syt1, other than a slight reduction in the fusion rate in the absence of PI(4,5)P_2_ (Fig. 5B-D). By contrast, the 4W mutant which binds the membrane more avidly essentially behaved like the wild-type C2AB, with the exception that the pore dilation ability of the 4W mutant was less dependent on the presence of PI(4,5)P_2_ (Fig 5C and see SI Appendix, fig. S7). Thus, calcium-induced membrane penetration of Syt1 C2 domains is required for pore expansion by Syt1.

### Mathematical modelling suggests that Syt1 and SNARE proteins cooperatively dilate fusion pores in a mechanical lever action

How do Syt1 and SNAREs cooperate to expand the pore in the presence of calcium? To help elucidate the mechanism, we developed a detailed mathematical model of the membrane fusion pore and the ApoE scaffold of the NLP in the presence of SNARE proteins and the C2AB domain of Syt1 (see SI Appendix for model details and parameters). The energetics of the fusion pore membrane are described in the classic Helfrich framework, with contributions from bending energy and membrane tension (97), while the ApoE scaffold is modelled by adapting the theory of elasticity (98)(see SI Appendix, fig. S8A). We obtained the minimum energy shape of the fusion pore with a given height and radius by solving the membrane shape equation (99) (see SI Appendix). To compare directly with the present experiments, we incorporate 4 SNARE complexes, each of which can either be in the *trans* configuration and free to roam the fusion pore, or else fully zippered in the *cis* configuration near the waist, Figure 6A (17). The model accounts for the SNARE zippering energy which favors full zippering (100, 101), and for crowding interactions among zippered SNAREs which favor partial unzippering into the *trans* state, an entropic effect (17, 102).

**Figure 6.**
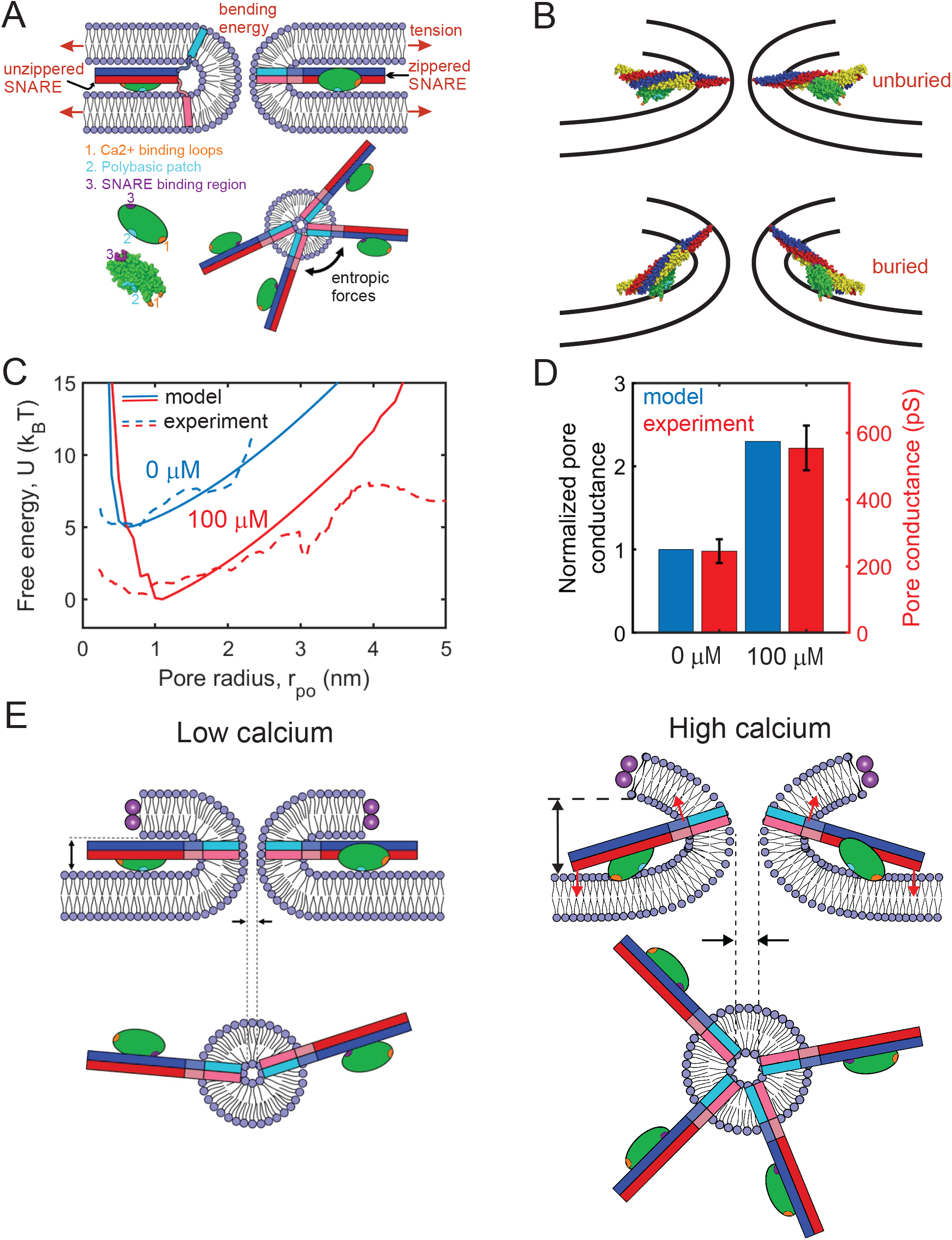
Mathematical model of the fusion pore in the presence of Syt1 and SNAREs suggests a mechanical calcium-triggered pore dilation mechanism. **(A)** Schematic of model. The membrane free energy has contributions from membrane tension and bending energy. SNARE complexes may be unzippered and free to roam laterally, or zippered and confined to the pore waist. Crowding among zippered SNARE complexes generates entropic forces that tend to enlarge the pore (top view, shown lower right). The Syt1 C2B domain (green ellipsoid) has a SNARE-binding region, a polybasic patch and Ca^2+^-binding loops. **(B)** Free energy-minimizing fusion pore shapes determined by solving the membrane shape equation in the presence and absence of constraints applied by the SNARE-C2B complex (see SI appendix). The C2B calcium-binding loops may either be unburied (top panel) or buried (lower panel) in the membrane. In the buried state the SNARE complex tilts upwards, expanding the fusion pore. The membrane shape constraint is evaluated using the SNARE-C2B complex crystal structure in a space filling representation. Both upper and lower panels depict situations in the presence of Ca^2+^. The model predicts the tilted configuration is strongly favored at high [Ca^2+^] following equilibration, while the untilted configuration is relevant to the kinetics that establish this equilibrium, and to experiments using low [Ca^2+^]. VAMP2, syntaxin, SNAP25 and the C2B domain are shown blue, red, yellow and green, respectively. The C2B hydrophobic membrane-inserting residues (V304, I367), polybasic patch (K326, K327) and SNARE binding region (R398, R399) are shown orange, cyan and purple, respectively. The protein structure was generated with PyMOL (145) using the SNARE-C2B crystal structure (PDB ID 5ccg) (88). The TMD of the SNARE complex (PDB ID 3hd7) (146) was incorporated using UCSF chimera software (147). **(C)** Model-predicted free energy and experimental apparent free energy versus pore radius without calcium and in the presence of excess calcium. **(D)** Model-predicted normalized conductances shown with experimentally measured values for comparison. Experimental data taken from Fig. 2B experiments including Ca^2+^ and PI(4,5)P_2_. (**E)** Pore dilation mechanism emerging from the model. Under conditions of low calcium concentration, the C2B domain is unburied, the SNARE complex lies parallel to the membrane and the membrane separation is set by the maximum thickness of the SNARE-C2B complex. At high calcium concentrations, the calcium binding loops penetrate the plasma membrane, rotating the C2B domain and the entire SNARE-C2B complex which exerts force (red arrows) on the upper and lower membranes of the fusion pore in a lever-like action. These forces increase the fusion pore height, which is coupled by membrane energetics to fusion pore dilation.

Syt1 C2B domains are assumed bound to each SNARE complex at the so-called primary interface identified in recent crystal structures (88, 89, 103) (Fig. 3A). For simplicity we first consider only the C2B domain in our model. When Ca^2+^ is bound to the C2B domain loops, the loops may be buried or unburied in the membrane with a relative probability that depends on the calcium concentration according to the Hill equation (36, 91). We use a Hill coefficient of 2.3, and the measured affinity of calcium for Syt1 in the presence of PI(4,5)P_2_-containing membranes (78). Without calcium, the loops are assumed unburied.

Thus, in the presence of calcium, the model permits two configurations of the SNARE-C2B complex, implemented according to the crystal structure (PDB ID 5ccg (88)), Figure 6B. **(1)**With bound Ca^2+^, the C2B complex can be in the buried state, in which the C2B polybasic patch lies ~ 0.5 nm from the membrane (95) and the C2B domain is anchored to the membrane by its calcium-binding loops, reported to penetrate ~1 nm deep (96). With these constraints, the SNAREpin is forced to tilt its C-terminus “upwards,” see Figure 6B; precise implementation of the constraints shows that the C2B anchoring tilts the SNAREpin upwards at ~ 15° to the plasma membrane, imposing a significant constraint on the shape of the fusion pore. We determined whether a given fusion pore geometry satisfied these constraints by directly comparing the structure of the SNARE-C2B complex with the shape of the fusion pore (see SI Appendix). Only fusion pores satisfying the shape constraints were accepted as possible pores. **(2)** With no bound calcium, the C2B polybasic patch (104) and the SNAREpins orient parallel to the plasma membrane. In this configuration, the SNARE-C2B complex imposes no constraints on the shape of the fusion pore. This unanchored state is also accessible when calcium is bound, with a probability that decreases with increasing calcium concentration.

Given the microscopically long pore lifetimes of seconds, we assumed the fusion pore-SNARE-Syt1 system has sufficient time to equilibrate. For a given pore radius, *r*_po_, we calculated the free energy by summing over all allowed SNARE-C2B configurations and all possible numbers of zippered SNAREs. Each state in this sum is weighted by its Boltzmann factor, yielding the free energy *U*(*r*_po_) and pore size distribution ~ exp[−*U*(*r*_po_)/*k*_B_*T*]. We assumed that the pore height is equal to the value that minimizes the free energy at a given radius *r*_po_, since other heights have small probability as the free energy increases rapidly as a function of pore height. The predicted free energy profiles with and without calcium are close to the experimental profiles, as shown in Figure 6C. We compared model and experimental free energies up to a maximum pore size of 4 nm, since sampling for larger pores was limited in the experiments. In agreement with experiment, introduction of calcium is predicted to increase the pore size fluctuations, as reflected by the broader distribution. From these pore size statistics, we calculated mean pore sizes and conductances. In the absence of calcium, the model predicts a mean fusion pore radius ~0.9 nm and a mean height ~9.0 nm, due to entropic crowding effects among *cis* SNARE complexes (17), Figure S8. These crowding effects expand the pore relative to the SNARE-free case, since a bigger pore increases the entropy of *cis*-SNAREs at the waist by providing more space.

When Ca^2+^ is introduced at high saturating concentrations, the model predicts a ~1.4-fold increase of pore radius to ~1.3 nm, or a ~2.3-fold increase in conductance, close to the experimentally measured ~2.2-fold increase (Figure 6D). The pore expansion mechanism is the constraint on the pore shape imposed by the SNARE-C2B complex. At low pore radii, the SNARE-C2B complex acts as a membrane inclusion that increases the height of the fusion pore, forcing the pore to adopt energetically unfavorable shapes, biasing the system towards large pore radii (Figure 6B, E, S8C). Due to membrane bending and tension, the fusion pore resists the lever action tending to increase its height and enlarge the pore.

However, these resistance forces are insufficient to rotate the SNARE-C2B lever complex and undo its pore-enlarging action, since this would require unanchoring of the Ca-binding loops from the membrane or dissociation of the SNARE-C2B domain binding interface. Both of these are sufficiently energetically unfavorable (75, 89) to overcome the fusion pore resistance forces (see SI Appendix for a detailed discussion). Figure 6D shows the predicted increase of normalized pore conductance in elevated Ca concentrations, compared with the experimental values. In summary, our model suggests a mechanism in which the SNARE-C2B complex is a calcium-triggered mechanical lever that enlarges the fusion pore in cooperation with entropic forces generated by SNARE complexes (Figure 6E). On addition of Ca^2+^, the C2B domain rotates and inserts its calcium binding loops into the membrane, tilting the SNARE complex so that it pushes the membrane surfaces further apart in a lever action. Since this increase in pore height would otherwise increase the net membrane bending energy, the pore diameter increases to offset this penalty (see SI Appendix).

### The model suggests the C2A domain is required for rapid pore dilation kinetics

Experiments have shown that the C2A domain plays a role in neurotransmitter release (105, 106), not just the C2B domain which was emphasized in the model above. Thus, we asked if the additional membrane binding energy provided by burying of the Ca^2+^-bound C2A loops is important to the pore enlargement effect. With calcium, the reported binding energy of the C2A domain to membranes is ~4 *k*_B_*T* from single-molecule force spectroscopy (75), and a dissociation constant 550 μM was measured using liposome titration (66). Estimating a ~ 1 nm capture radius, the latter translates to a ~8 *k*_B_*T* binding energy. Thus, C2A-membrane binding energies lie in the range 4 *k*_B_*T* < *E*_C2A_< 8 *k*_B_*T*.

A simple argument shows that this contribution from C2A is not required for the equilibrium enlarged pore state of Figure 6B, as follows. The energetic cost to disrupt the C2B-membrane interaction is ~9-18 k_B_T per Syt molecule, from single-molecule force spectroscopy and surface force apparatus measurements (75, 107). This is much bigger than the work done by SNARE-Syt levers to increase the pore height, ~9 k_B_T from our model (see SI Appendix and Fig. S8E). Thus, C2B anchoring alone is sufficient to oppose the membrane pushing forces, with no need for the additional 4-8 *k*_B_*T* provided by the C2A domain.

Since the steady state pore appears not to require C2A, we asked if C2A was important for the kinetics of pore dilation that establish this enlarged pore, which we measured experimentally (Figure 4D). Thus, we developed a simple model of the kinetics that establish many SNARE-Syt1 levers (Figure 6B, bottom panel) starting from unanchored, untilted levers (Figure 6B, top panel). The model suggests that C2A does indeed play a significant role: in its absence, much reduced pore dilation rates are predicted. Thus, the kinetic model suggests the full C2AB domain is required for normal pore dilation.

For simplicity we consider only the critical kinetic step, insertion of the first lever, i.e. the first fully-zippered SNARE-C2AB complex with C2 domains anchored to the membrane that pushes the pore membranes apart. The model considers three states corresponding to the sequence of events to establish this first lever. (i) the initial state, with Ca^2+^ bound to the C2 domains, in which the SNARE-C2AB complex is oriented parallel to the membranes and the C2 domains are unanchored. (ii) The final state with the lever in place, with calcium-bound C2 domains anchored to the tCell membrane. (iii) A high energy transition state with anchored calcium-bound C2 domains in which the membranes have not been pushed apart, and the SNARE-C2AB complex is under duress.

Relative to the initial state, the energetic cost of the transition state is *E*^⋆^ − *E*_C2X_, where *E*^⋆^ is the energy of the SNARE-Syt complex under duress, and *E*_C2X_ the C2 domain membrane anchoring energy (*X* = *B*, *A* or *AB*). Thus the ratio of the rates at which the transition state is established from the initial state in the presence and absence of the C2A domain is the ratio of Boltzmann factors exp[−(*E*^⋆^ − *E*_C2AB_))/*k*_B_*T*]/exp[−(*E*^⋆^ − *E*_C2B_)]/*k*_B_*T*). Assuming additive energies, *E*_C2A_ = *E*_C2AB_ − *E*_C2B_, this simplifies to exp(*E*_C2A_/*k*_B_*T*). Thus with only the C2B domain the rate to transit from the top panel of Figure 6B (untilted SNARE-Syt1 lever) to the bottom panel (first upwardly tilting SNARE-Syt1 lever, expanding the pore) is reduced by a factor exp(*E*_C2A_/*k*_B_*T*) ≈ 50 – 3000 relative to the rate for the full C2AB domain. This is a good estimate for the reduction factor for the entire dilation process, since insertion of subsequent levers is much easier as the membrane is now raised so there is no high duress high energy transition state. Moreover, the membrane load will now be shared by several levers.

Our experiments using wild-type C2AB measured a pore dilation time ~25 ms (Figs. 4C,D), so the waiting time to observe the fully expanded pore is predicted by the model to increase to ~1 − 90 s in the absence of C2A. Given the experimentally measured pore lifetimes of ~ 15 s (see SI Appendix, fig. S3D), this suggests that fully dilated pores would rarely be observed in the absence of the C2A domain.

## DISCUSSION

Membrane fusion occurs in stages. First, membranes are brought into close apposition to overcome repulsive hydration forces. Second, a small, nascent fusion pore forms, connecting the fusing membranes. Third, the initial small pore expands to allow passage of cargo molecules (2–4). Among different stages of membrane fusion, pore expansion can be energetically one of the costliest (81, 82, 108–110). Consistent with this notion, fusion pores connecting protein-free lipid bilayers fluctuate, flicker open-closed, and eventually reseal unless external energy is supplied in the form of membrane tension (111), while the initial fusion pore during biological membrane fusion is a metastable structure whose dynamics are regulated by cellular processes (3–7, 10, 83, 112–115).

Syt1 is involved in both the pore opening and pore expansion stages during calcium-triggered exocytosis. *Before* membrane fusion, Syt1 was proposed to regulate membrane apposition (54–58), preventing fusion pore opening at low calcium by maintaining the membranes >5-8 nm apart, halting complete SNARE zippering. Upon calcium binding to Syt1, this distance is reduced to <5 nm (55), sufficient for SNAREs to complete their zippering and initiate fusion. Other mechanisms, such as calcium-dependent release of an inhibition of complete SNARE assembly by Syt1 (50), or concerted action of an oligomeric complex containing Syt1, SNAREs, and additional proteins (59, 60), have also been proposed for the pore opening stage. It has also been proposed that during this stage, curvature generation by insertion of Syt1’s hydrophobic loops into the membranes may contribute to pore opening (37, 52, 53).

*After* fusion pore opening, Syt1 contributes to the dilation of the nascent fusion pore (32, 37), but the mechanisms for this regulation have remained even less clear. Several Syt1-independent mechanisms regulating fusion pore dynamics have recently emerged. First, membrane tension promotes fusion pore dilation during exocytosis, often through cytoskeleton-plasma membrane interactions (116–118). Second, neuronal/exocytic SNARE proteins promote fusion pore dilation by providing entropic forces due to molecular crowding at the pore’s waist (17), consistent with the observation that increased SNARE availability results in larger, or faster expanding pores (15, 17, 26–28). Third, during yeast vacuole-vacuole fusion, increased fusogen volume has been suggested as a mechanism that stabilizes fusion pores (83, 119). However, these mechanisms cannot explain fusion pore dilation during exocytosis, because none are calcium-dependent, in contrast to exocytic fusion pore expansion (32, 120–122). Previous reconstituted single-pore measurements by Lai et al. (39) and Das et al. (123) found Syt1 and calcium promoted expansion of SNARE-mediated fusion pores. In the former study, pores were detected indirectly through passage of large probe molecules (39), while the latter study reported that the larger, stable pores formed in the presence of Syt1, calcium and PI(4,5)P_2_ could be closed by dissociation of the SNARE complexes by the ATPase NSF, but not by a soluble cytoplasmic fragment of the v-SNARE VAMP2 (123). However, the mechanism of fusion pore dilation remained unclear.

Here, we found that Syt1 has roles in both fusion pore formation and dilation, consistent with studies in secretory cells (32, 37) and in previous reconstitutions (39, 123), and we focused on pore dilation mechanisms. Syt1 promotes expansion of SNARE-induced fusion pores in a calcium- and acidic lipid-dependent manner. When PI(4,5)P_2_ is present, increasing free Ca^2+^ leads to pores with larger mean open-pore conductance. Fusion pore expansion by Syt1 also likely relies on Syt1’s interactions with the neuronal SNARE complex, because when we used C2AB domains with mutations (R398Q,R399Q) designed to disrupt the “primary” interaction interface with the SNARE complex (88, 89), the pore dilation function of Syt1 C2AB was largely reduced (Fig. 3). The same mutations were previously shown to greatly reduce evoked release from hippocampal neurons (55, 88, 90), possibly by disrupting the interaction of Syt1 C2B with SNAREs (55, 88). The most relevant interactions in which these residues engage is however not completely resolved, so results of mutagenesis of these residues must be interpreted with caution. For example, this mutation did not have a significant effect in the co-IP experiments of Syt1 with SNAREs (88), but it did have substantial effects on the ability of Syt1 C2B to bridge two membranes (90). In addition, in the presence of polyvalent ions such as Mg^2+^ and ATP, Syt1 was found not to bind to SNAREs (124), but ATP did not have any effect in a tethered-liposome fusion assay (88). Later work by Wang et al. (103) examined these interactions in the presence of membranes and SNARE complexes, and suggested that the C2B (R398 R399)–SNARE complex interaction is Ca^2+^ independent (*K*_*d*_ < 1 μM in the presence of PI(4,5)P_2_ in the membranes), stronger than the C2B (R398 R399)–acidic lipid interactions, persists during insertion of the Ca^2+^-binding loops into the membrane, and occurs simultaneously with the calcium-independent interactions of the C2B polybasic patch with PI(4,5)P_2_ containing membranes. Wang et al. showed ATP/Mg^2+^ does not disrupt Syt1-SNARE complex interactions in the absence of Ca^2+^, but not in its presence (103). Thus, although the most likely interpretation is that mutation of R398,R399 disrupts Syt1 C2B-SNARE complex binding through the primary interface, other possibilities cannot be excluded.

A mathematical model suggests the major contribution of Syt1 to pore dilation is through its mechanical modulation of the fusion pore shape. Syt-SNARE complexes introduce non-local constraints on the fusion pore shape, making larger pores more energetically favorable. How does the non-local constraint come about? Previous work showed calcium binding to isolated Syt1 C2 domains leads to insertion of the hydrophobic residues at the tips of both of the the calcium-binding loops into the membrane (41, 96, 104, 125) (however, see ref. (126)). In the presence of PI(4,5)P_2_, calcium-bound C2B assumes a conformation in which its long axis is tilted with respect to the membrane normal, as it interacts with the membrane simultaneously through its calcium binding loops and the polybasic patch (K324-327) bound to PI(4,5)P_2_ (95, 127). When present, C2B also binds the t-SNAREs Stx1 and SNAP25, with its long axis parallel to the SNARE bundle, in a calcium-independent manner (88, 103). In this orientation, the polybasic patch on C2B (K324-327) is free to interact with acidic lipids on the target membrane (88). At low, resting amounts of calcium, the calcium-free SNARE-C2B complex is therefore expected to lie parallel to the membrane, with the C2B domain simultaneously interacting with target membrane acidic lipids and the SNARE complex (88) (Fig. 6). By contrast, in the presence of high calcium, the calcium-bound C2B domain will tend to reorient such that its hydrophobic top loops insert into the target membrane, resulting in a tilting of the SNARE complex of ~15 degrees, which alters the pore shape (Fig. 6). The resultant pore size increase quantitatively accounts for the conductance increase in the presence of Syt1, and its requirements for intact calcium- and SNARE-binding regions on C2B. At intermediate calcium levels, the mean pore radius is expected to have an intermediate value, as the Syt1 molecules would be activated by calcium for a fraction of the time only. Thus, our results may explain why initial fusion pore size and its expansion rate increase as intracellular calcium increases (32, 37, 120–122). In addition, regulation of the fusion pore shape including interbilayer distance may be a general mechanism to stabilize fusion pores against re-closure, as a similar mechanism was observed during yeast vacuole-vacuole fusion (83, 119).

Mutations of the hydrophobic residues at the tips of the calcium-binding loops of the C2 domains (M173, F234, V304 and I367) designed to increase or decrease the affinity of Syt1 for calcium-induced membrane binding were previously interpreted largely in terms of the ability of these mutants to generate membrane curvature. Indeed, the rates of fusion between liposomes (52, 53) and exocytosis (37, 128) correlate well with the curvature-generation ability of the Syt1 mutants. By contrast, here the correlation between the curvature-generation ability of the mutants and pore expansion was not strong, with the 4W mutant with enhanced membrane tubulation activity (37, 52) having a similar effect as wild-type C2AB. Modeling supported the idea that curvature-generation by Syt1 membrane penetration is not needed to explain how Syt1 promotes pore expansion.

We also explored how Syt1 affects pore dilation kinetics as a function of calcium. We found pore expansion rate increases with increasing [Ca^2+^]_free_, with a similar dependence on calcium as the mean open-pore conductance (Fig. 4A,D), from ~3-12 nS/s at low calcium (0-30 μM), to 20-25 nS/s at high calcium (40-100 μM). Modelling suggests the C2A domain of Syt1 is critical for rapid expansion of the fusion pore, by contributing to the total binding energy of Syt1 C2 domains to acidic membranes. By comparison, in secretory cells the pore opens suddenly (129) before continuing to expand at a slower rate. In horse eosinophils stimulated by intracellular application of GTP-γ-S, pores were found to expand, on average, at 19 nS/s, 40 nS/s, and 89 nS/s at low (<10 nM), 1.5 μM, and 10 μM Ca^2+^, respectively (120), consistent with a later study (122). Pore expansion rates were 5-10 nS/s for rat mast cells, with higher rates at high calcium (121), and varied from 15 to 50 nS/s for bovine chromaffin cells (25, 130, 131). Lower rates (~7 nS/s) were observed in excised patch recordings (131). A rate of ~98 nS/s was reported for rat chromaffin cells overexpressing myosinII (132). These pore expansion rates, and the increasing rates with increasing calcium, are remarkably consistent with our findings.

Our findings also recapitulate the observation that during exocytosis, fusion pore fluctuations increase with intracellular calcium (133). A mathematical model suggests that this originates in the cooperative mechanical effects of Syt1 and SNAREs which exert outward expansive forces on the fusion pore. These forces oppose the inward force that results from the intrinsic tendency of the protein-free fusion pore to close down due to membrane bending and tension effects (17). The net inward force is thus lowered, leading to a broader distribution of pore sizes and bigger fluctuations.

In several neuronal preparations, the maximal rate of secretion scales as 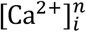 with *n* ≈ 4 (71, 72, 134–137), while in our system the mean open pore conductance or the rate of fusion pore expansion (Fig. 4A,D) are consistent with a Hill relationship with cooperativity ~2 and calcium affinity ~20 μM, taken from studies of purified recombinant Syt1 C2AB binding to lipid bilayers (78). There are several reasons for these differences. Most importantly, the maximal rates of secretion measured in neurons or neuroendocrine chromaffin cells (which possess a similar fusion machinery) is due to the rapid fusion of a pool of docked and primed vesicles called the readily-releasable pool (RRP) (138–140). Vesicles acquire fusion-competence at low, resting calcium levels (≲ 0.1 μM). When the calcium concentration near release sites increases rapidly in response to stimulation, fusion from the RRP ensues within milliseconds. Docking (~30 s) and priming (~10 s) are much slower events (138, 139) and require tethering and priming factors such as Munc13 and Munc18 (1, 141). There is no RRP or its equivalent in our assay: nanodiscs dock and fuse with the target cell membrane under a constant calcium level throughout the measurement and key components of the docking and priming machinery such as Munc13 and Munc18 are absent in our minimalistic reconstitution. Thus, the steep calcium-dependence of the maximal rate of RRP secretion observed in neurons is not directly comparable to the fusion pore opening or expansion kinetics in our assay in which discs fuse with the target membrane under conditions of constant calcium levels, very low fusion rates, and absence of docking and priming factors.

The nanodisc-cell fusion assay is tuned for sensitivity to post-fusion stages. Unfortunately, like other electrical or electrochemical methods that generate a signal only after fusion pore opening, our assay cannot directly detect pre-fusion stages. In particular, the delay between docking and fusion of nanodiscs, and the molecular configurations leading to the opening of the initial fusion pore are currently not known. Until a better understanding of such pre-fusion stages is achieved, our post-fusion studies should be interpreted with care. A detailed discussion of the relevance and limitations of nanodisc-based single pore measurements in relation to exocytotic fusion pores monitored in secretory cells can be found in ref. (2).

In neurons and many neuroendocrine cells, fusion is triggered by a brief calcium transient. The finding that fusion pore dilation is calcium sensitive suggests that the pore size, expansion rate, and duration can be modulated by calcium dynamics. Thus, weak stimulations that result in brief calcium transients would be more likely to lead to small fusion pores and slow release, and strong stimulations would conversely result in larger and faster dilating pores. This behavior is indeed observed in neurons (9), and in neuroendocrine cells (7, 142). In this framework, different Syt isoforms would affect fusion pore dynamics differently, depending on their ability to reorient with respect to the membranes, their interactions with the SNAREs, and their calcium affinities.

## MATERIALS AND METHODS

### Recombinant protein expression and purification

Expression and purification of the constructs used are described in SI Appendix, Supplementary Materials and Methods.

### Reconstitution of synaptotagmin-1 and VAMP2 into nanodiscs

Eight copies of VAMP2 (~four per face) were incorporated into nanolipoprotein particles (vNLP8) as previously described (16, 17, 69). The protocol was modified to produce nanolipoprotein particles co-reconstituted with full-length Syt1 and VAMP2 (vsNLP), as detailed in SI Appendix, Supplementary Materials and Methods.

### Stable flipped SNARE cell lines

Stable “tCell” HeLa cell lines expressing flipped t-SNAREs (rat Syntaxin-1, residues 186-288, and rat SNAP-25, residues 2-206) and the nuclear marker CFP-nls (cyan fluorescent protein fused to nuclear localization signal) were a generous gift from the Rothman laboratory (143) and cultured as previously reported (16, 17). Details are given in SI Appendix, Supplementary Materials and Methods.

### Single fusion pore conductance assay

All recordings were done as previously described (16, 17), and detailed in SI Appendix, Supplementary Materials and Methods. Estimations of fusion rates and pore properties are explained in the SI Appendix, Supplementary Materials and Methods, along with evidence that ATP-dependent channel activity is absent and that cell membrane potential changes are negligible during recordings.

### Statistical analysis

Details are given in SI Appendix, Supplementary Materials and Methods, and in figure legends.

## Data availability

Data and Matlab routines used for analyses are available upon reasonable request.

## ACKNOWLEDGEMENTS

We thank Ekaterina Stroeva and Shyam Krishnakumar (Rothman Laboratory, Yale University) for help with reconstitution of full-length Syt1 into nanodiscs. This work was supported by National Institute of General Medical Sciences and National Institute of Neurological Disorders and Stroke of the National Institutes of Health under award numbers R01NS113236 and R01EY010542 (to EK) and R01GM117046 (to BOS). The content is solely the responsibility of the authors and does not necessarily represent the official views of the National Institutes of Health. We acknowledge computing resources from Columbia University’s Shared Research Computing Facility project. We thank Rui Su and members of the Karatekin lab for helpful discussions.

## AUTHOR CONTRIBUTIONS

ZW and EK conceived the study and wrote the manuscript, with input from other co-authors. ND, ST, ZM, and BO conceived the model, performed calculations, and contributed to the writing.

## ADDITIONAL INFORMATION

SI Appendix

**Correspondence and requests for materials** should be addressed to B.O. or E.K. (lead contact).

## REFERENCES

1. Brunger AT, Choi UB, Lai Y, Leitz J, & Zhou Q (2018) Molecular Mechanisms of Fast Neurotransmitter Release. Annu Rev Biophys 47:469–497.

2. Karatekin E (2018) Toward a unified picture of the exocytotic fusion pore. FEBS Lett 592(21):3563–3585.

3. Sharma S & Lindau M (2018) The fusion pore, 60 years after the first cartoon. FEBS Lett 592(21):3542–3562.

4. Chang CW, Chiang CW, & Jackson MB (2017) Fusion pores and their control of neurotransmitter and hormone release. J Gen Physiol 149(3):301–322.

5. Alabi AA & Tsien RW (2013) Perspectives on kiss-and-run: role in exocytosis, endocytosis, and neurotransmission. Annu Rev Physiol 75:393–422.

6. Collins SC, et al. (2016) Increased Expression of the Diabetes Gene SOX4 Reduces Insulin Secretion by Impaired Fusion Pore Expansion. Diabetes 65(7):1952–1961.

7. Fulop T, Radabaugh S, & Smith C (2005) Activity-dependent differential transmitter release in mouse adrenal chromaffin cells. Journal of Neuroscience 25(32):7324–7332.

8. He LM, Wu XS, Mohan R, & Wu LG (2006) Two modes of fusion pore opening revealed by cell-attached recordings at a synapse. Nature 444(7115):102–105.

9. Pawlu C, DiAntonio A, & Heckmann M (2004) Postfusional control of quantal current shape. Neuron 42(4):607–618.

10. Staal RGW, Mosharov EV, & Sulzer D (2004) Dopamine neurons release transmitter via a flickering fusion pore. Nature Neuroscience 7(4):341–346.

11. Chapochnikov NM, et al. (2014) Uniquantal release through a dynamic fusion pore is a candidate mechanism of hair cell exocytosis. Neuron 83(6):1389–1403.

12. Gandhi SP & Stevens CF (2003) Three modes of synaptic vesicular recycling revealed by single-vesicle imaging. Nature 423(6940):607–613.

13. Lisman JE, Raghavachari S, & Tsien RW (2007) The sequence of events that underlie quantal transmission at central glutamatergic synapses. Nat Rev Neurosci 8(8):597–609.

14. Verstreken P, et al. (2002) Endophilin mutations block clathrin-mediated endocytosis but not neurotransmitter release. Cell 109(1):101–112.

15. Bao H, et al. (2018) Dynamics and number of trans-SNARE complexes determine nascent fusion pore properties. Nature 554(7691):260–263.

16. Wu Z, et al. (2016) Nanodisc-cell fusion: control of fusion pore nucleation and lifetimes by SNARE protein transmembrane domains. Sci Rep 6:27287.

17. Wu Z, et al. (2017) Dilation of fusion pores by crowding of SNARE proteins. Elife 6:e22964.

18. Wu Z, Thiyagarajan S, O’Shaughnessy B, & Karatekin E (2017) Regulation of Exocytotic Fusion Pores by SNARE Protein Transmembrane Domains. Front Mol Neurosci 10:315.

19. Han X, Wang CT, Bai JH, Chapman ER, & Jackson MB (2004) Transmembrane segments of syntaxin line the fusion pore of Ca2+-triggered exocytosis. Science 304(5668):289–292.

20. Bretou M, Anne C, & Darchen F (2008) A fast mode of membrane fusion dependent on tight SNARE zippering. J Neurosci 28(34):8470–8476.

21. Kesavan J, Borisovska M, & Bruns D (2007) v-SNARE actions during Ca(2+)-triggered exocytosis. Cell 131(2):351–363.

22. Dhara M, et al. (2016) v-SNARE transmembrane domains function as catalysts for vesicle fusion. Elife 5:e17571.

23. Ngatchou AN, et al. (2010) Role of the synaptobrevin C terminus in fusion pore formation. Proc Natl Acad Sci U S A 107(43):18463–18468.

24. Weber T, et al. (1998) SNAREpins: Minimal machinery for membrane fusion. Cell 92(6):759–772.

25. Fang QH, et al. (2008) The role of the C terminus of the SNARE protein SNAP-25 in fusion pore opening and a model for fusion pore mechanics. P Natl Acad Sci USA 105(40):15388–15392.

26. Acuna C, et al. (2014) Microsecond dissection of neurotransmitter release: SNARE-complex assembly dictates speed and Ca(2)(+) sensitivity. Neuron 82(5):1088–1100.

27. Gucek A, et al. (2016) Dominant negative SNARE peptides stabilize the fusion pore in a narrow, release-unproductive state. Cell Mol Life Sci 73(19):3719–3731.

28. Zhao Y, et al. (2013) Rapid structural change in synaptosomal-associated protein 25 (SNAP25) precedes the fusion of single vesicles with the plasma membrane in live chromaffin cells. Proc Natl Acad Sci U S A 110(35):14249–14254.

29. Segovia M, et al. (2010) Push-and-pull regulation of the fusion pore by synaptotagmin-7. P Natl Acad Sci USA 107(44):19032–19037.

30. Zhang Z, Hui EF, Chapman ER, & Jackson MB (2010) Regulation of Exocytosis and Fusion Pores by Synaptotagmin-Effector Interactions. Molecular Biology of the Cell 21(16):2821–2831.

31. Zhang ZJ, Zhang Z, & Jackson MB (2010) Synaptotagmin IV Modulation of Vesicle Size and Fusion Pores in PC12 Cells. Biophysical Journal 98(6):968–978.

32. Wang CT, Bai J, Chang PY, Chapman ER, & Jackson MB (2006) Synaptotagmin-Ca2+ triggers two sequential steps in regulated exocytosis in rat PC12 cells: fusion pore opening and fusion pore dilation. J Physiol 570(Pt 2):295–307.

33. Wang CT, et al. (2003) Different domains of synaptotagmin control the choice between kiss-and-run and full fusion. Nature 424(6951):943–947.

34. Wang CT, Lu JC, Chapman ER, Martin TFJ, & Jackson MB (2003) Synaptotagmin IV induces long-duration kiss-and-run exocytosis through small fusion pores. Biophysical Journal 84(2):209a-209a.

35. Wang CT, et al. (2001) Synaptotagmin modulation of fusion pore kinetics in regulated exocytosis of dense-core vesicles. Science 294(5544):1111–1115.

36. Bai J, Wang CT, Richards DA, Jackson MB, & Chapman ER (2004) Fusion pore dynamics are regulated by synaptotagmin*t-SNARE interactions. Neuron 41(6):929–942.

37. Lynch KL, et al. (2008) Synaptotagmin-1 utilizes membrane bending and SNARE binding to drive fusion pore expansion. Mol Biol Cell 19(12):5093–5103.

38. Rao TC, et al. (2014) Distinct fusion properties of synaptotagmin-1 and synaptotagmin-7 bearing dense core granules. Mol Biol Cell 25(16):2416–2427.

39. Lai Y, et al. (2013) Fusion pore formation and expansion induced by Ca2+ and synaptotagmin 1. Proc Natl Acad Sci U S A 110(4):1333–1338.

40. Geppert M, et al. (1994) Synaptotagmin I: a major Ca2+ sensor for transmitter release at a central synapse. Cell 79(4):717–727.

41. Chapman ER (2008) How does synaptotagmin trigger neurotransmitter release? Annu Rev Biochem 77:615–641.

42. Bhalla A, Tucker WC, & Chapman ER (2005) Synaptotagmin isoforms couple distinct ranges of Ca2+, Ba2+, and Sr2+ concentration to SNARE-mediated membrane fusion. Mol Biol Cell 16(10):4755–4764.

43. Bhalla A, Chicka MC, & Chapman ER (2008) Analysis of the synaptotagmin family during reconstituted membrane fusion. Uncovering a class of inhibitory isoforms. J Biol Chem 283(31):21799–21807.

44. Pinheiro PS, Houy S, & Sorensen JB (2016) C2-domain containing calcium sensors in neuroendocrine secretion. J Neurochem 139(6):943–958.

45. Volynski KE & Krishnakumar SS (2018) Synergistic control of neurotransmitter release by different members of the synaptotagmin family. Curr Opin Neurobiol 51:154–162.

46. Craxton M (2010) A manual collection of Syt, Esyt, Rph3a, Rph3al, Doc2, and Dblc2 genes from 46 metazoan genomes--an open access resource for neuroscience and evolutionary biology. BMC Genomics 11:37.

47. Sugita S, Shin OH, Han W, Lao Y, & Sudhof TC (2002) Synaptotagmins form a hierarchy of exocytotic Ca(2+) sensors with distinct Ca(2+) affinities. EMBO J 21(3):270–280.

48. Hui E, et al. (2005) Three distinct kinetic groupings of the synaptotagmin family: candidate sensors for rapid and delayed exocytosis. Proc Natl Acad Sci U S A 102(14):5210–5214.

49. Xu J, Mashimo T, & Sudhof TC (2007) Synaptotagmin-1, -2, and -9: Ca(2+) sensors for fast release that specify distinct presynaptic properties in subsets of neurons. Neuron 54(4):567–581.

50. Brunger AT, Leitz J, Zhou Q, Choi UB, & Lai Y (2018) Ca(2+)-Triggered Synaptic Vesicle Fusion Initiated by Release of Inhibition. Trends Cell Biol 28(8):631–645.

51. Sudhof TC (2013) Neurotransmitter Release: The Last Millisecond in the Life of a Synaptic Vesicle. Neuron 80(3):675–690.

52. Martens S, Kozlov MM, & McMahon HT (2007) How synaptotagmin promotes membrane fusion. Science 316(5828):1205–1208.

53. Hui EF, Johnson CP, Yao J, Dunning FM, & Chapman ER (2009) Synaptotagmin-Mediated Bending of the Target Membrane Is a Critical Step in Ca2+-Regulated Fusion. Cell 138(4):709–721.

54. Rothman JE, Krishnakumar SS, Grushin K, & Pincet F (2017) Hypothesis - buttressed rings assemble, clamp, and release SNAREpins for synaptic transmission. FEBS Lett 591(21):3459–3480.

55. Chang S, Trimbuch T, & Rosenmund C (2018) Synaptotagmin-1 drives synchronous Ca(2+)-triggered fusion by C2B-domain-mediated synaptic-vesicle-membrane attachment. Nat Neurosci 21(1):33–40.

56. van den Bogaart G, et al. (2011) Synaptotagmin-1 may be a distance regulator acting upstream of SNARE nucleation. Nat Struct Mol Biol 18(7):805–812.

57. Seven AB, Brewer KD, Shi L, Jiang QX, & Rizo J (2013) Prevalent mechanism of membrane bridging by synaptotagmin-1. Proc Natl Acad Sci U S A 110(34):E3243–3252.

58. Lin CC, et al. (2014) Control of membrane gaps by synaptotagmin-Ca2+ measured with a novel membrane distance ruler. Nat Commun 5:5859.

59. Bello OD, et al. (2018) Synaptotagmin oligomerization is essential for calcium control of regulated exocytosis. Proc Natl Acad Sci U S A 115(32):E7624–E7631.

60. Tagliatti E, et al. (2020) Synaptotagmin 1 oligomers clamp and regulate different modes of neurotransmitter release. Proc Natl Acad Sci U S A 117(7):3819–3827.

61. Chernomordik LV & Kozlov MM (2008) Mechanics of membrane fusion. Nature Structural & Molecular Biology 15(7):675–683.

62. Kozlov MM, McMahon HT, & Chernomordik LV (2010) Protein-driven membrane stresses in fusion and fission. Trends in Biochemical Sciences 35(12):699–706.

63. Zhang Z, et al. (2011) Release mode of large and small dense-core vesicles specified by different synaptotagmin isoforms in PC12 cells. Mol Biol Cell 22(13):2324–2336.

64. Lynch KL & Martin TF (2007) Synaptotagmins I and IX function redundantly in regulated exocytosis but not endocytosis in PC12 cells. J Cell Sci 120(Pt 4):617–627.

65. Osterberg JR, et al. (2015) Membrane Docking of the Synaptotagmin 7 C2A Domain: Electron Paramagnetic Resonance Measurements Show Contributions from Two Membrane Binding Loops. Biochemistry 54(37):5684–5695.

66. Voleti R, Tomchick DR, Sudhof TC, & Rizo J (2017) Exceptionally tight membrane-binding may explain the key role of the synaptotagmin-7 C2A domain in asynchronous neurotransmitter release. Proc Natl Acad Sci U S A 114(40):E8518–E8527.

67. Dudzinski NR, Wu Z, & Karatekin E (2019) A Nanodisc-Cell Fusion Assay with Single-Pore Sensitivity and Sub-millisecond Time Resolution. Methods Mol Biol 1860:263–275.

68. Shi L, et al. (2012) SNARE Proteins: One to Fuse and Three to Keep the Nascent Fusion Pore Open. Science 335(6074):1355–1359.

69. Bello OD, Auclair SM, Rothman JE, & Krishnakumar SS (2016) Using ApoE Nanolipoprotein Particles To Analyze SNARE-Induced Fusion Pores. Langmuir 32(12):3015–3023.

70. Hille B (2001) Ion channels of excitable membranes (Sinauer, Sunderland, Mass.) 3rd Ed pp xviii, 814 p.

71. Schneggenburger R & Neher E (2000) Intracellular calcium dependence of transmitter release rates at a fast central synapse. Nature 406(6798):889–893.

72. Schneggenburger R & Neher E (2005) Presynaptic calcium and control of vesicle fusion. Curr Opin Neurobiol 15(3):266–274.

73. Voets T (2000) Dissection of three Ca2+-dependent steps leading to secretion in chromaffin cells from mouse adrenal slices. Neuron 28(2):537–545.

74. Chanaday NL & Kavalali ET (2018) Presynaptic origins of distinct modes of neurotransmitter release. Curr Opin Neurobiol 51:119–126.

75. Ma L, et al. (2017) Single-molecule force spectroscopy of protein-membrane interactions. Elife 6:e30493.

76. Perez-Lara A, et al. (2016) PtdInsP2 and PtdSer cooperate to trap synaptotagmin-1 to the plasma membrane in the presence of calcium. Elife 5:e15886.

77. Honigmann A, et al. (2013) Phosphatidylinositol 4,5-bisphosphate clusters act as molecular beacons for vesicle recruitment. Nat Struct Mol Biol 20(6):679–686.

78. Bai J, Tucker WC, & Chapman ER (2004) PIP2 increases the speed of response of synaptotagmin and steers its membrane-penetration activity toward the plasma membrane. Nat Struct Mol Biol 11(1):36–44.

79. Colquhoun D & Hawkes AG (1995) The Principles of the Stochastic Interpretation of Ion-Channel Mechanisms. Single-Channel Recording, eds Sakmann B & Neher E (Plenum Press, New York), 2 Ed, pp 397–482.

80. Schiavo G, Matteoli M, & Montecucco C (2000) Neurotoxins affecting neuroexocytosis. Physiol Rev 80(2):717–766.

81. Jackson MB (2009) Minimum membrane bending energies of fusion pores. J Membr Biol 231(2-3):101–115.

82. Cohen FS & Melikyan GB (2004) The energetics of membrane fusion from binding, through hemifusion, pore formation, and pore enlargement. J Membr Biol 199(1):1–14.

83. D’Agostino M, Risselada HJ, Endter LJ, Comte-Miserez V, & Mayer A (2018) SNARE-mediated membrane fusion arrests at pore expansion to regulate the volume of an organelle. EMBO J 37(19).

84. Mackler JM, Drummond JA, Loewen CA, Robinson IM, & Reist NE (2002) The C(2)B Ca(2+)-binding motif of synaptotagmin is required for synaptic transmission in vivo. Nature 418(6895):340–344.

85. Shin OH, Xu J, Rizo J, & Sudhof TC (2009) Differential but convergent functions of Ca2+ binding to synaptotagmin-1 C2 domains mediate neurotransmitter release. Proc Natl Acad Sci U S A 106(38):16469–16474.

86. Nishiki T & Augustine GJ (2004) Dual roles of the C2B domain of synaptotagmin I in synchronizing Ca2+-dependent neurotransmitter release. J Neurosci 24(39):8542–8550.

87. Borden CR, Stevens CF, Sullivan JM, & Zhu Y (2005) Synaptotagmin mutants Y311N and K326/327A alter the calcium dependence of neurotransmission. Mol Cell Neurosci 29(3):462–470.

88. Zhou Q, et al. (2015) Architecture of the synaptotagmin-SNARE machinery for neuronal exocytosis. Nature 525(7567):62–67.

89. Zhou Q, et al. (2017) The primed SNARE-complexin-synaptotagmin complex for neuronal exocytosis. Nature 548(7668):420–425.

90. Xue M, Ma C, Craig TK, Rosenmund C, & Rizo J (2008) The Janus-faced nature of the C(2)B domain is fundamental for synaptotagmin-1 function. Nat Struct Mol Biol 15(11):1160–1168.

91. Radhakrishnan A, Stein A, Jahn R, & Fasshauer D (2009) The Ca2+ affinity of synaptotagmin 1 is markedly increased by a specific interaction of its C2B domain with phosphatidylinositol 4,5-bisphosphate. J Biol Chem 284(38):25749–25760.

92. Davis AF, et al. (1999) Kinetics of synaptotagmin responses to Ca2+ and assembly with the core SNARE complex onto membranes. Neuron 24(2):363–376.

93. Martens S & McMahon HT (2008) Mechanisms of membrane fusion: disparate players and common principles. Nat Rev Mol Cell Biol 9(7):543–556.

94. Chapman ER & Davis AF (1998) Direct interaction of a Ca2+-binding loop of synaptotagmin with lipid bilayers. J Biol Chem 273(22):13995–14001.

95. Kuo W, Herrick DZ, & Cafiso DS (2011) Phosphatidylinositol 4,5-bisphosphate alters synaptotagmin 1 membrane docking and drives opposing bilayers closer together. Biochemistry 50(13):2633–2641.

96. Herrick DZ, Sterbling S, Rasch KA, Hinderliter A, & Cafiso DS (2006) Position of synaptotagmin I at the membrane interface: cooperative interactions of tandem C2 domains. Biochemistry 45(32):9668–9674.

97. Helfrich W (1973) Elastic properties of lipid bilayers: theory and possible experiments. Z Naturforsch C 28(11):693–703.

98. Landau LD & Lifshitz EM (1986) Theory of Elasticity (Third Edition), eds Lifshitz EM, Kosevich AM, & Pitaevskii LP (Butterworth-Heinemann, Oxford), pp 38–86.

99. Ou-Yang Z & Helfrich W (1989) Bending Energy of Vesicle Membranes - General Expressions for the 1st, 2nd, and 3rd Variation of the Shape Energy and Applications to Spheres and Cylinders. Phys Rev A 39(10):5280–5288.

100. Gao Y, et al. (2012) Single reconstituted neuronal SNARE complexes zipper in three distinct stages. Science 337(6100):1340–1343.

101. Ma L, et al. (2015) Munc18-1-regulated stage-wise SNARE assembly underlying synaptic exocytosis. Elife 4:e09580.

102. Mostafavi H, et al. (2017) Entropic forces drive self-organization and membrane fusion by SNARE proteins. Proc Natl Acad Sci U S A 114(21):5455–5460.

103. Wang S, Li Y, & Ma C (2016) Synaptotagmin-1 C2B domain interacts simultaneously with SNAREs and membranes to promote membrane fusion. Elife 5.

104. Kuo W, Herrick DZ, Ellena JF, & Cafiso DS (2009) The calcium-dependent and calcium-independent membrane binding of synaptotagmin 1: two modes of C2B binding. J Mol Biol 387(2):284–294.

105. Striegel AR, et al. (2012) Calcium binding by synaptotagmin’s C2A domain is an essential element of the electrostatic switch that triggers synchronous synaptic transmission. J Neurosci 32(4):1253–1260.

106. Lee J, Guan Z, Akbergenova Y, & Littleton JT (2013) Genetic analysis of synaptotagmin C2 domain specificity in regulating spontaneous and evoked neurotransmitter release. J Neurosci 33(1):187–200.

107. Gruget C, et al. (2018) Rearrangements under confinement lead to increased binding energy of Synaptotagmin-1 with anionic membranes in Mg(2+) and Ca(2). FEBS Lett 592(9):1497–1506.

108. Chizmadzhev YA, Cohen FS, Shcherbakov A, & Zimmerberg J (1995) Membrane mechanics can account for fusion pore dilation in stages. Biophys J 69(6):2489–2500.

109. Ryham RJ, Ward MA, & Cohen FS (2013) Teardrop shapes minimize bending energy of fusion pores connecting planar bilayers. Phys Rev E Stat Nonlin Soft Matter Phys 88(6):062701.

110. Nanavati C, Markin VS, Oberhauser AF, & Fernandez JM (1992) The Exocytotic Fusion Pore Modeled as a Lipidic Pore. Biophysical Journal 63(4):1118–1132.

111. Chanturiya A, Chernomordik LV, & Zimmerberg J (1997) Flickering fusion pores comparable with initial exocytotic pores occur in protein-free phospholipid bilayers. P Natl Acad Sci USA 94(26):14423–14428.

112. Doreian BW, Fulop TG, Meklemburg RL, & Smith CB (2009) Cortical F-Actin, the Exocytic Mode, and Neuropeptide Release in Mouse Chromaffin Cells Is Regulated by Myristoylated Alanine-rich C-Kinase Substrate and Myosin II. Molecular Biology of the Cell 20(13):3142–3154.

113. Barg S, et al. (2002) Delay between fusion pore opening and peptide release from large dense-core vesicles in neuroendocrine cells. Neuron 33(2):287–299.

114. Hanna ST, et al. (2009) Kiss-and-run exocytosis and fusion pores of secretory vesicles in human beta-cells. Pflugers Archiv-European Journal of Physiology 457(6):1343–1350.

115. MacDonald PE, Braun M, Galvanovskis J, & Rorsman P (2006) Release of small transmitters through kiss-and-run fusion pores in rat pancreatic beta cells. Cell Metab 4(4):283–290.

116. Bretou M, et al. (2014) Cdc42 controls the dilation of the exocytotic fusion pore by regulating membrane tension. Mol Biol Cell 25(20):3195–3209.

117. Kozlov MM & Chernomordik LV (2015) Membrane tension and membrane fusion. Curr Opin Struct Biol 33:61–67.

118. Wen PJ, et al. (2016) Actin dynamics provides membrane tension to merge fusing vesicles into the plasma membrane. Nat Commun 7:12604.

119. D’Agostino M, Risselada HJ, Lurick A, Ungermann C, & Mayer A (2017) A tethering complex drives the terminal stage of SNARE-dependent membrane fusion. Nature 551(7682):634–638.

120. Hartmann J & Lindau M (1995) A novel Ca(2+)-dependent step in exocytosis subsequent to vesicle fusion. FEBS Lett 363(3):217–220.

121. Fernandez-Chacon R & Alvarez de Toledo G (1995) Cytosolic calcium facilitates release of secretory products after exocytotic vesicle fusion. FEBS Lett 363(3):221–225.

122. Scepek S, Coorssen JR, & Lindau M (1998) Fusion pore expansion in horse eosinophils is modulated by Ca2+ and protein kinase C via distinct mechanisms. EMBO J 17(15):4340–4345.

123. Das D, Bao H, Courtney KC, Wu LX, & Chapman ER (2020) Resolving kinetic intermediates during the regulated assembly and disassembly of fusion pores. Nature Communications 11(1).

124. Park Y, et al. (2015) Synaptotagmin-1 binds to PIP(2)-containing membrane but not to SNAREs at physiological ionic strength. Nat Struct Mol Biol 22(10):815–823.

125. Bradberry MM, Bao H, Lou X, & Chapman ER (2019) Phosphatidylinositol 4,5-bisphosphate drives Ca(2+)-independent membrane penetration by the tandem C2 domain proteins synaptotagmin-1 and Doc2beta. J Biol Chem 294(28):10942–10953.

126. Bykhovskaia M (2021) SNARE complex alters the interactions of the Ca(2+) sensor synaptotagmin 1 with lipid bilayers. Biophys J 120(4):642–661.

127. Perez-Lara A, et al. (2016) PtdInsP2 and PtdSer cooperate to trap synaptotagmin-1 to the plasma membrane in the presence of calcium. Elife 5.

128. Rhee JS, et al. (2005) Augmenting neurotransmitter release by enhancing the apparent Ca2+ affinity of synaptotagmin 1. Proc Natl Acad Sci U S A 102(51):18664–18669.

129. Breckenridge LJ & Almers W (1987) Currents through the Fusion Pore That Forms during Exocytosis of a Secretory Vesicle. Nature 328(6133):814–817.

130. Berberian K, Torres AJ, Fang QH, Kisler K, & Lindau M (2009) F-Actin and Myosin II Accelerate Catecholamine Release from Chromaffin Granules. Journal of Neuroscience 29(3):863–870.

131. Dernick G, Gong LW, Tabares L, Alvarez de Toledo G, & Lindau M (2005) Patch amperometry: high-resolution measurements of single-vesicle fusion and release. Nat Methods 2(9):699–708.

132. Neco P, et al. (2008) Myosin II contributes to fusion pore expansion during exocytosis. J Biol Chem 283(16):10949–10957.

133. Zhou Z, Misler S, & Chow RH (1996) Rapid fluctuations in transmitter release from single vesicles in bovine adrenal chromaffin cells. Biophysical Journal 70(3):1543–1552.

134. Dodge FA & Rahamimo.R (1967) Co-Operative Action of Calcium Ions in Transmitter Release at Neuromuscular Junction. J Physiol-London 193(2):419–&.

135. Sun J, et al. (2007) A dual-Ca2+-sensor model for neurotransmitter release in a central synapse. Nature 450(7170):676–682.

136. Kochubey O, Lou X, & Schneggenburger R (2011) Regulation of transmitter release by Ca(2+) and synaptotagmin: insights from a large CNS synapse. Trends Neurosci 34(5):237–246.

137. Heidelberger R, Heinemann C, Neher E, & Matthews G (1994) Calcium dependence of the rate of exocytosis in a synaptic terminal. Nature 371(6497):513–515.

138. Kaeser PS & Regehr WG (2017) The readily releasable pool of synaptic vesicles. Curr Opin Neurobiol 43:63–70.

139. Sorensen JB (2004) Formation, stabilisation and fusion of the readily releasable pool of secretory vesicles. Pflugers Arch 448(4):347–362.

140. Rizzoli SO & Betz WJ (2005) Synaptic vesicle pools. Nat Rev Neurosci 6(1):57–69.

141. Rizo J (2018) Mechanism of neurotransmitter release coming into focus. Protein Sci 27(8):1364–1391.

142. Cardenas AM & Marengo FD (2016) How the stimulus defines the dynamics of vesicle pool recruitment, fusion mode, and vesicle recycling in neuroendocrine cells. J Neurochem 137(6):867–879.

143. Giraudo CG, Eng WS, Melia TJ, & Rothman JE (2006) A clamping mechanism involved in SNARE-dependent exocytosis. Science 313(5787):676–680.

144. Lyubimov AY, et al. (2016) Advances in X-ray free electron laser (XFEL) diffraction data processing applied to the crystal structure of the synaptotagmin-1 / SNARE complex. Elife 5:e18740.

145. Schrodinger, LLC (2015) The PyMOL Molecular Graphics System, Version 1.8.

146. Stein A, Weber G, Wahl MC, & Jahn R (2009) Helical extension of the neuronal SNARE complex into the membrane. Nature 460(7254):525–U105.

147. Pettersen EF, et al. (2004) UCSF Chimera--a visualization system for exploratory research and analysis. J Comput Chem 25(13):1605–1612.

