## Supplementary Information for "The neuronal calcium sensor Synaptotagmin-1 and SNARE proteins cooperate to dilate fusion pores"

### *SI APPENDIX*

#### SUPPLEMENTARY MATERIALS AND METHODS

##### Recombinant protein expression and purification

All SNARE and Synaptotagmin-1 constructs used were generous gifts from James E. Rothman, unless noted otherwise. Plasmid pET32a-Trx-His6X-ApoE422K, used to express the N-terminal 22 kDa fragment of apolipoprotein E4 (residues 1–199, ApoE422K), was kindly provided by Dr Nicholas Fischer, Lawrence Livermore National Laboratory, CA (1, 2). Full-length VAMP2 (residues x1-116 in plasmid pET-SUMO-VAMP2) and ApoE422K were expressed and purified as previously described (3, 4). Rat Syt1 residues 96-421 corresponding to cytoplasmic C2AB domains were expressed from a pET28a-SUMO-synaptotagmin1 vector. $C2AB^{R398,399Q}$, $C2AB^{D309N}$ and $C2AB^{K326,327A}$ were generated from the wild-type sequence using the QuickChange site-directed mutagenesis kit (Stratagene, La Jolla, CA). $C2AB^{4W}$ and $C2AB^{4A}$ were prepared using QuikChange Multi Site-Directed Mutagenesis Kit (Stratagene, La Jolla, CA). Wild type C2AB and all mutated version of C2AB were expressed in BL21 (DE3) and purified as previously reported (5). Full length Syt1 (pET28a-SUMO-synaptotagmin 1, residues 57-421) was expressed in BL2 (DE3) at 37°C to optical density 0.8 (at 600 nm) and induced with 1 mM isopropyl β-D-thiogalactoside (IPTG) for 4 hours. Cells were then lysed by a cell disruptor (Avestin, Ottawa, CA) and lysates were clarified by centrifugation (35,000 rpm at 4°C for 30 min using a Beckman-Coulter Ti45 rotor and 70 ml polycarbonate tubes, corresponding to 142,160 × g). The supernatant was incubated with Ni-NTA agarose (Qiagen, Valencia, CA) overnight at 4°C. Protein bound beads were washed by buffer A (25 mM HEPES, pH 7.4, 400 mM KCl, 0.5 mM tris-2-carboxyethyl phosphine (TCEP)) supplemented with 50 mM imidazole and 1% Octylglucoside (OG). The protein was first separated from beads using buffer A supplemented with 400 mM Imidazole and 4% OG. Then the His-SUMO tag was cleaved by SUMO proteinase at 4°C for 2 hours. The protein was diluted 4 times by dilution buffer (25 mM HEPES, 0.5 mM TCEP and 4% OG) and then immediately loaded into Mono S™ 5/50G column (GE Healthcare Bio-Sciences, Pittsburgh, PA). The full-length Syt1 was washed out by high salt buffer (25 mM HEPEs, 1 M KCl, 0.5 mM TCEP and 1% OG). After concentration determination using the Bradford assay (Bio Rad, Hercules, CA), the samples were aliquoted, flash frozen by plunging into liquid nitrogen, and stored at -80 °C for future use.

##### Co-reconstitution of Synaptotagmin-1 and VAMP2 into nanolipoprotein particles (NLPs)

Eight copies each of VAMP2 and full-length Synaptotagmin-1 (Syt1) (~four per face each) were incorporated into nanolipoprotein particles (vsNLP8) following previous protocols for reconstitution of VAMP2 alone (3, 4, 6). A mixture of palmitoyl-2-oleoylphosphatidylcholine (POPC) and 1,2-dioleoyl phosphatidylserine (DOPS) (85:15 molar ratio) dissolved in a chloroform-methanol mixture (2:1 by volume) was dried under nitrogen flow, then placed under vacuum for 2 hrs. All lipids were purchased from Avanti Polar Lipids (Alabaster, AL). The lipid film was re-suspended in 25 mM HEPES, pH 7.4, 140 mM KCl, 1 mM TCEP buffer with 1% OG supplemented by the desired amount of full length syt1 and VAMP2. The mixture was vortexed for 1 hr at room temperature followed by the addition of ApoE422K and vortexed for another half hour at room temperature and then 3 hours at cold room. The ApoE422K:VAMP2: syt1: lipid ratio for vsNLPs was 1:2:2:180. Excess detergent was removed using SM-2 bio-beads (Bio-Rad) overnight at 4°C with gentle shaking. The assembled vsNLPs were purified using size-exclusion chromatography using a Superose™ 6, 10/300 GL column (GE Healthcare Bio-Sciences, Pittsburgh, PA). Collected samples were concentrated using Amicon Ultra (30 KDa cutoff) centrifugal filter units, and analyzed by SDS-PAGE with Coomassie staining. The size distribution of the NLPs was determined for every batch of production using transmission electron microscopy (JEM-1400, JEOL, MA, USA). This allowed estimating the average number of ApoE copies per disc as before (4, 6), using previously published information about the number of ApoE copies as a function of disc size (2).

The copy numbers of Syt1 and VAMP2 per disc were then estimated from the quantification of Syt1- or VAMP2-to-ApoE ratio using densitometry (ImageJ, NIH).

**Stable flipped SNARE cell lines**

Stable "tCell" HeLa cell lines expressing flipped t-SNAREs (rat Syntaxin-1, residues 186-288, and rat SNAP-25, residues 2-206) and the nuclear marker CFP-nls (cyan fluorescent protein fused to nuclear localization signal) were a generous gift from the Rothman laboratory (7) and cultured as previously reported (3, 4). The flipped SNARE constructs used in the generation of these lines, pBI-flipped Syntaxin-1 (186-288)-flipped SNAP-25-IRES-CFP-nls, are schematically shown in SI Appendix, fig. S1I (7, 8). The pBI expression vector is a bidirectional mammalian expression vector of the Tet-Off gene expression system that allows co-regulation of the synthesis of two gene products in stoichiometric amounts (9). The cells were cultured in DMEM (4500 mg/L glucose, L-glutamine, sodium pyruvate and sodium bicarbonate) and 10% (v/v) fetal calf serum at 37°C.

**PI(4,5)$P_2$ incorporation, and immunostaining**

Where indicated, short-chain diC8-PI(4,5)$P_2$ (Echelon Biosciences Inc., Salt Lake City, UT) (1 mM stock solution, dissolved in water), was added to the cell culture medium to a final concentration of 20 μM and incubated 20 min at 37°C. Cells were then washed three times using extracellular buffer (ECS: 125 mM NaCl, 4 mM KCl, 2 mM $CaCl_2$, 1 mM $MgCl_2$, and 10 mM HEPES, pH adjusted to 7.2 with NaOH and 10 mM glucose added freshly).

For assessing diC8-PI(4,5)$P_2$ incorporation into the outer leaflet of the plasma membrane and lifetime, after 20 min incubation with the lipid, cells were rinsed thoroughly with phosphate buffered saline (PBS) supplemented with 10% goat serum, and kept at 37 ºC with the same solution for different durations. Mouse monoclonal anti-PI(4,5)$P_2$ primary antibodies (Echelon Biosciences Inc., Utah) were added to the cells at time points of 0, 40, and 80 minutes and incubated 1 hr at 37 °C. Then cells were fixed with 4% paraformaldehyde (Electron Microscopy Sciences, PA) for 20 min at room temperature before addition of goat anti-mouse IgM heavy chain secondary antibody conjugated with Alexa Fluor 647. Control cells that were not incubated with diC8-PI(4,5)$P_2$ were treated similarly and fixed by 4% paraformaldehyde. Some cells were then permeabilized by 0.5% saponin (Sigma, MO) to allow access of the antibody to the inner leaflet of the plasma membrane where endogenous PI(4,5)$P_2$ resides. Cells were blocked for 30 min with PBS supplemented with 10% goat serum, followed by incubation with anti-PI(4,5)$P_2$ primary antibody for 1 hour at 37°C. After three successive washes in PBS, cells were incubated with the secondary antibody as above. All groups of cells were washed three times with PBS and mounted on a glass slide with mounting medium (ProLong Gold Antifade Mountant with DAPI, Molecular Probes, OR). Fluorescence images were collected using a spinning disk confocal microscope (model TiE, Nikon, Japan, equipped with a Yokogawa CSU-W1 spinning disc head and CFI Plan Apochromat Lambda 60x/1.4 oil immersion objective). Images were analyzed using ImageJ software. We drew a region of interest (ROI) around cells using the freehand ROI tool and measured the mean pixel intensity in the ROI. We then subtracted the intensity from a nearby region not containing any cells to define the background subtracted pixel intensity to define $\Delta F$ in Fig. 1B. For each condition, 10 regions of interest encompassing cells were analyzed from 3-6 independent preparations.

**Whole-cell conductance of flipped t-SNARE cells**

We measured whole-cell current responses to step changes in membrane potential under voltage-clamp, from HeLa cells stably expressing flipped t-SNAREs (Fig. S11A). Currents were averaged for 27 cells and plotted against voltage (Fig. S11B). Pipettes were filled with intracellular solution (in mM): 134 KCl, 2 $MgCl_2$, 1 $CaCl_2$, 10 HEPEs and 10 EGTA (pH is adjusted to 7.2 by KOH).

**Single fusion pore conductance assay**

All recordings were done as previously described (3, 4). Briefly, a dish with cultured tCells was rinsed using ECS, then mounted on a Thermo-Plate (Tokai Hit, Shizuoka-ken, Japan) pre-set to 37°C. tCells were visualized with an inverted Olympus IX71 microscope (Olympus Corp., Waltham, MA) using a ThorLabs USB3.0 digital camera (UI-3240CP-NIR-GL-TI) controlled by ThorCam software (ThorLabs, Newton, NJ). Recording pipettes (borosilicate glass, BF 150-86-10, Sutter Instruments, Novato, CA) were pulled using a model P-1000 pipette puller (Sutter Instruments, Novato, CA) and polished using a micro-forge (MF-830, Narishige, Tokyo, Japan). The pipette solution (PipSol) contained: 125 mM NaCl, 4 mM KCl, 1 mM MgCl2, 10 mM HEPES, 26 mM TEA-Cl, 2 mM ATP (freshly added), 0.5 mM EGTA, pH adjusted to 7.2 by NaOH and the indicated free calcium (0-500 μM) was adjusted by 0.1 M Calcium Standard Solutions (Thermo Fisher Scientific, Waltham, MA). Free calcium was calculated using MaxChelator (https://somapp.ucdmc.ucdavis.edu/pharmacology/bers/maxchelator/CaMgATPEGTA-TS.htm) taking into account ATP, $Mg^{2+}$, ionic strength, temperature, and pH. The pipette was pre-filled by PipSol and then back filled with PipSol supplemented with nanodiscs with or without additional C2AB. All voltage-clamp recordings were made using a HEKA EPC10 Double USB amplifier (HEKA Elektronik Dr. Schulze GmbH, Lambrecht/Pfalz, Germany), controlled by Patchmaster software (HEKA). Current signals were digitized at 20 kHz and filtered at 3 kHz. The recording traces were exported to MatLab (MathWorks, Natick, MA) and analyzed as previously described in detail (3, 4).

**Detection of fusion pore currents**

As described previously (3, 4, 10), the pipette tip was initially filled with ~1 μl of disc-free buffer and back-filled with NLPs suspended in the same buffer (final [NLP]≈100 nM, 120 μM lipids). This allowed establishing a tight seal ($R_{seal} > 10$ GOhm) with high success rate and recording a stable baseline before the NLPs diffused to the membrane patch and started fusing with it a few to several min later. All cell-attached recordings were performed using a holding potential of $V_p = -40$ mV relative to bath. With a cell resting membrane potential of $-56 \pm 7$ mV (mean ± S.D., n=36 (3)), this provided 16 mV driving force across the patch membrane. The pipette solution had resistivity 0.60 Ohm.m, measured using a conductivity cell (DuraProbe, Orion Versa Star, Thermo Scientific).

After a good seal was established on a cell, currents were recorded under voltage-clamp for 800 s, in 40 s sweeps, with a sampling rate of 20 kHz using a HEKA EPC10 Double USB amplifier (HEKA Elektronik), controlled by Patchmaster software (HEKA). The analysis pipeline started with initial off-line inspection of the traces in PatchMaster. Traces with activity were exported to Matlab (Mathworks, Natick, MA) where they were analyzed in more detail using an interactive graphical user interface we developed to help identify, crop and process single fusion pore currents (3, 4, 10). Traces with excessive noise or unstable baseline were excluded from analysis. Exported traces were low-pass filtered (280 Hz cutoff) and frequencies due to line voltage were removed using notch filtering. Zero phase shift digital filtering algorithms (Matlab Signal Processing Toolbox function filtfilt) were employed to prevent signal distortion. Filtered traces were averaged in blocks of 80 points (125 Hz final bandwidth) to achieve rms baseline noise $\lesssim 0.2$ pA. Currents $I$ for which $|I| > 2.0$ pA for at least 250 ms were accepted as fusion pore current bursts. During a burst, rapidly fluctuating currents often returned to baseline multiple times, i.e. pores flickered. To quantify pore flickering, we defined currents $< -0.25$ pA and lasting $\geq$ 60 ms (15 points) as open pores and currents not meeting these criteria as closed. For a given burst, the number of open periods was equal to the number of flickers, $N_{flickers}$. The burst lifetime is defined as the time from the initial to the final point detected using the criteria above. Current bursts spaced >5 s apart by a quiet baseline were assigned to separate bursts, since the typical lifetime of well-isolated bursts is 5-10 s. An example of a current burst is shown in Fig. S1J with the threshold current, detected open sub-periods, and the burst lifetime indicated. Examples of entire 800 s recordings are shown in Fig. S2. The MatLab programs used in analysis and the data are available upon request.

**Estimation of fusion rate**

To estimate the fusion rate for each recording (i.e. the rate at which current bursts appeared), we counted the number of current bursts that fit the set criteria (current amplitude >2 pA for at least 250 ms) and divided this number by the duration of the recording. Examples are shown in Fig. S2A,B. These per-cell rates were averaged over all cells to estimate the average rate of fusion ("pores/min") and its standard deviation for a given condition. Standard error of the mean was calculated as the standard deviation divided by the square root of the number of cells. Periods during which the baseline was not stable were excluded from this analysis. For individual cells, the number of well-isolated pores varied from 0 to 22. Many recordings ended with what seemed to be currents from overlapping fusion pores (Fig. S2B). Such end-of-record currents were also excluded, since they could also be attributed to a loose seal. Thus, the fusion rates we report may underestimate the true rates, especially for conditions where fusion activity was high.

We checked that increasing or decreasing the concentration of v-SNARE NLPs in the pipette solution increased or decreased the fusion rate, respectively. Indeed, we found there is good linear correlation between the v-SNARE NLP concentration and the fusion rate, as shown in Fig. S2C.

As an alternative estimate of the fusion rate, for every condition, we summed all detected pores, $N_{tot}$, and the analysis time $\tau_{tot}$ over all cells (excluding portions with noisy/unstable baseline), and calculated the total number of pores divided by the total analysis time, $\dot{F}_{tot} = N_{tot}/\tau_{tot}$. The results of this estimate were close to the ones described above, as shown in Fig. S2D.

**Estimation of fusion pore parameters**

The number of flickers, $N_{flickers}$, and the burst lifetime, $T_o$, were defined as explained above. The flicker rate was defined as the number of flickers divided by the burst lifetime for individual pores. The pore open probability, $P_o$, is defined as the total time the pore was in the "open" state divided by the burst lifetime, $T_o$, for individual pores. We converted current to conductance by dividing every point in a current trace by the transmembrane voltage $V_m = V_{cell} - V_p = -16$ mV, where $V_p$ is the pipette potential (-40 mV) and $V_{cell} = -56$ mV as indicated above. To calculate the open-pore conductance, $G_{po}$, and its statistics, we used pore open-state values, denoted by the subscript “po”. Similarly, we used pore open-state values to calculate the distributions of open-pore conductance values and radii. For the distributions in Figure 2C, D, S4, S6, and S7, we first computed the probability density functions (PDFs) for individual pores using a fixed bin width for all, then averaged these to give equal weight to all pores. All distribution fits (e.g. Fig. S3E, F) were performed using Matlab Statistics Toolbox functions fitdist or mle, using maximum likelihood estimation. Open-pore conductance values were used point-by-point to estimate the open-pore radii, by approximating the pore as a cylinder and using the expression (11) $r_{po} = \left(\rho\lambda G_{po}/\pi\right)^{1/2}$, where $\rho$ is the resistivity of the solution, $\lambda = 15$ nm is the length of the cylinder, and $G_{po}$ is the open-pore conductance.

For assessing statistical significance when comparing sample means, we used the student's t-test when the parameters were normally distributed, or the nonparametric two-sample Kolmogorov-Smirnov test otherwise (ttest2 or kstest2, Matlab Statistics Toolbox), as indicated in figure legends. We considered each single-pore measurement a biological replicate.

**Estimation of fusion pore expansion rates**

For aligning and averaging conductance traces in Fig. 4C, we shifted the time axis such that $t = 0$ corresponded to the first data point in a burst. We estimated pore expansion rate as the 10-90% rise time from the baseline to the level of conductance reached within the first 100 ms after pore opening, divided by the time it took for this rise using the Matlab function "slewrate".

As an alternative, we also fit a straight line to each of the aligned and averaged conductance rise, for the initial 16 ms of the rise, and used the slope of the line as an estimate of the pore expansion rate. Pore expansion rates estimated from these slopes as a function of [$Ca^{2+}$] resulted is a plot very similar to the one in Fig. 4D obtained using the slew rate estimate above. The differences in the slopes can be largely explained by differences in the conductance level reached within ~16 ms. After filtering and block averaging, the spacing between successive points is 4 ms in individual traces, corresponding to a Nyquist frequency of 125 Hz. That is, we should be able to faithfully reconstruct signals varying on a time scale of 8 ms or slower. However, slopes calculated over a 16 ms span are still likely to be limited by our resolution to some degree, because we cannot detect finer kinetic details during this period. Thus, the pore expansion rates we averaged over 16 ms may be underestimates of the true rates and finer details of the kinetics cannot be resolved.

**No evidence for ATP-dependent channel activity in flipped t-SNARE cells**

For cell-attached single-pore measurements, ATP was included in the pipette solutions. HeLa cells were reported to express ATP-dependent P2 receptors (12). To test whether ATP-dependent channel activation is present in the flipped t-SNARE cells, we recorded currents from cell-attached, voltage-clamped patches from these cells in the absence and presence of ATP (nanodiscs were absent). Pipette solutions were the same as for single fusion pore measurements with 100 µM free calcium, except for ATP as noted. Both in the absence and presence of ATP (2 mM), we occasionally had patches that displayed channel-like activity (Fig. S9). We conclude that the activity of these channels is not regulated by ATP, consistent with an earlier report (12).

In addition, we note that the vast majority of channel-like currents as in Fig. S9 are excluded from our analysis of fusion pore currents, because their lifetime is too short ($< 250$ ms), their amplitude is too low ($< -2$ pA), or both, and therefore do not significantly affect our results.

**Cell membrane potential changes do not significantly distort cell-attached fusion pore recordings**

It has been reported that cell membrane potential may change under some conditions during cell-attached recordings (e.g. see Fenwick et al. (13)). In such recordings, the single-channel conductance $g$ is underestimated (compared to its true value $G$), unless $G_{cell} \gg G_{patch}$, where $G_{cell}$ is the cell membrane conductance, and $G_{patch}$ is the patch conductance (14). Given that the ratio of the cell area to patch area is typically $A_{cell}/A_{patch} > 100$ or $1000$, and that membrane capacitance is proportional to membrane area, one would expect the requirement for $G_{cell} \gg G_{patch}$ is easily satisfied. However, for some small cells, sometimes it is found that ions passing through single channels can change the cell membrane potential, hence the potential across the patch (13, 15). The effect was found only occasionally for some cells from the same preparation, and for small cells. Fenwick et al. (13) suggested that some local damage to the membrane patch during the formation of the gigaseal may occur in some cases.

Several lines of evidence suggest cell membrane potential changes do not significantly distort our cell-attached recordings:

1) If the cell membrane potential changed due to currents passing through fusion pores, such currents would depolarize the cell membrane and reduce the transmembrane voltage across the patch ($V_m = V_{cell} - V_p$). Indeed a 15-20 mV depolarization of the cell membrane from its starting value of -56 mV would bring it close to $V_p$ and largely abolish the driving force $V_m$ for current flow across the patch. This would result in larger currents at the beginning of a pore event compared to its end, and this effect would be strongest for the condition producing the largest pores, i.e. in the presence of full-length Syt1 (with calcium and PI(4,5)$P_2$). To test this idea, we aligned pore currents to the beginning or end of events, and averaged them, as shown in Fig. S10. We do not find large differences between averaged traces aligned either way.

2) The hallmark of cell membrane potential changes in single-channel recordings is "relaxation" of single-channel currents when the channels open and close (13). In our recordings, we do not see such relaxation, even after aligning pore current to their moment of closure and averaging them as shown in Fig. S10.

3) In whole-cell voltage-clamp recordings, we found $G_{cell} = 5 - 6$ nS for flipped t-SNARE cells (Fig. S11). Thus, the condition $G_{cell} \gg G_{patch}$ is satisfied in most of our recordings. Even in the presence of Syt1, $G_m$ is nearly 10 times larger than the average conductance ($G_{patch} \approx 600$ pS). The range of mean open-pore currents and transmembrane voltages comprising 95% of the data values for C2AB in the presence of PI(4,5)$P_2$ and 100 μM calcium are indicated as a red-colored box on Fig. S11B.

**Statistical analysis**

For fusion rates and other parameters that followed a normal distribution, the two-sample t-test was used. For open-pore conductance, or other parameter distributions which do not follow a normal distribution, the two-sample Kolmogorov-Smirnov test was used for pair-wise comparisons. In Fig. 1B we used one-way ANOVA, followed by a multiple comparison test (using the Tukey-Kramer criterion). For all statistical analyses, we used Matlab Statistics and Machine Learning Toolbox (MathWorks). Details are provided in figure legends.

**Mathematical model of the fusion pore with SNAREs and Synaptotagmin-1**

The shape of the fusion pore between the nanolipoprotein particle (NLP) and the tCell membrane is determined by minimizing the Helfrich energy (16); this is achieved by numerically solving the membrane shape equation with constraints fixing the pore radius $r_{\mathrm{po}}$ and height $h$ , defined to be the separation between the NLP and tCell membrane (see subsection *"Numerical method for solving the membrane shape equation"*). We assume that each side of the NLP contains $N$ v-SNAREs and that all are available to associate with the t-SNAREs in the tCell membrane and contribute to pore expansion. Out of $N$ SNAREs, $N_{\mathrm{Z}}$ denotes the number of fully zippered SNAREs. For a given set of values $(r_{\mathrm{po}}, h, N, N_{\mathrm{Z}})$ the total free energy of the fusion pore is

$$U_{\mathrm{tot}}(r_{\mathrm{po}}, h, N, N_{\mathrm{Z}}) = U_{\mathrm{mb}} + U_{\mathrm{hyd}} + U_{\mathrm{SNARE}} + U_{\mathrm{scaffold}}, \qquad (1)$$

where $U_{\mathrm{mb}}$, $U_{\mathrm{hyd}}$, $U_{\mathrm{SNARE}}$, and $U_{\mathrm{scaffold}}$ stand for the membrane energy of the pore, the energy due to hydration forces between the NLP and tCell membranes, the free energy associated with the SNAREpins, and the free energy of the deformed NLP scaffold, respectively.

Each SNARE is bound to a Syt1 C2AB domain at the primary interface between the SNARE and the C2B domain (17). The C2AB domain calcium binding loops can be unburied or buried in the membrane with a probability that depends on calcium concentration (see *"Calcium dependent pore conductance"* below). In the unburied state, the C2B polybasic patch is facing the tCell membrane and parallel to it (18). In this orientation the C2B-attached SNARE is also roughly parallel to the membrane. In the buried state, the C2B domain anchors to the membrane by insertion of its calcium binding loops ~1 nm into the membrane (19), and the polybasic patch is distanced ~0.5 nm from the membrane (20). With respect to the unburied state, this configuration has a rotated C2B domain, which is attached to the SNARE complex at the primary interface, such that the SNARE complex is somewhat raised above the membrane and is concomitantly tilted by ~ 15° with respect to the membrane plane (Figure 6B, main text). This tilt angle is measured by taking the inverse sine of the ratio between the SNARE motif length projected on the vertical axis of the pore and the length of the SNARE motif. Thus, the C2B domain acts as a fulcrum about which the SNARE lever pivots. This configuration imposes a geometric constraint on the pore, leading to increased pore radius and height.

We calculate the pore conductance in the absence of calcium where C2B domains are unburied and in saturating calcium levels when all C2B domain are buried, and use these values to predict the mean pore conductance as described in the subsubsection *Model- predicted pore conductance* below. The expressions for the different energy terms in eq. (1) are described below.

**<u>Membrane free energy</u>**

The NLP and tCell membranes are modelled as a planar bilayer with diameter $D$ and an infinite planar bilayer, respectively, and both are at a constant surface tension. The membrane free energy of the fusion pore is given by (16)

$$U_{\mathrm{mb}}(r_{\mathrm{po}}, h) = \min\left\{\int_{A_{mb}} \left[\frac{\kappa}{2}(2C)^2 + \gamma\right] dA\right\} \tag{2}$$

where $\kappa$ is the membrane's bending modulus, $\gamma$ is the membrane tension, $C$ is the local mean curvature, and the integration is taken over the area of the membrane mid plane of the pore, $A_{\mathrm{mb}}$. The shape of the fusion pore is determined by solving a set of differential equations whose solutions minimize the membrane energy subject to the constraints of a fixed pore height $h$ and pore radius $r_{\mathrm{po}}$ (see SI subection "*Numerical method of solving the membrane shape equation*"). A term associated with the Gaussian curvature is omitted throughout our analysis because it depends only on the membrane topology. We set the bending modulus to a typical value of $\kappa = 20\ k_BT$. We set the value of $\gamma$ to $0.1$ pN nm$^{-1}$, which was obtained as a best fit parameter by comparing model-predicted pore energies with results from a similar experimental setup where fusion between the tCell and the NLP was induced by SNAREs alone (4), Figure S8B.

**<u>Numerical method of solving the membrane shape equation</u>**

We used the MATLAB® differential equation solver "bvp4c" with an absolute tolerance of $10^{-6}$ and a relative tolerance of $10^{-4}$ to solve the membrane shape equation (MathWorks, Natick, MA). This method requires that the equation be rendered as a set of first order ordinary differential equations. The process by which we determined these differential equations is described here.

For simplicity, the fusion pore is assumed to be axisymmetric in our calculations, i.e. symmetric under rotations about the $z$-axis. Given this assumption, the membrane energy can be written

$$U_{\mathrm{mb}} = \int_0^{2\pi} \mathrm{d}\theta \int_0^L \mathrm{d}s\, r(s) \left[\frac{\kappa}{2}(2\,C)^2 + \gamma\right] = 2\pi \int_0^L \mathrm{d}s\, r(s) \left[\frac{\kappa}{2}(2\,C)^2 + \gamma\right], \tag{3}$$

where $\theta$ is the azimuthal angle, $s$ measures the arclength along a meridian of the fusion pore (i.e., a curve of constant $\theta$), $r(s)$ is the distance of a given point on the fusion pore from the $z$-axis, and $L$ is the total arclength of the meridian. We add to this expression two Lagrange multiplier terms which fix the definition of $s$ as the arclength,

$$\begin{aligned} U_{\mathrm{mb}} = \; 2\pi \int_0^L \mathrm{d}s \Big\{ r(s) \left[\frac{\kappa}{2}(2\,C)^2 + \gamma\right] + f_r(s)\big(r'(s) - \cos(\phi(s))\big) \\ + f_z(s)\big(z'(s) - \sin(\phi(s))\big)\Big\}, \end{aligned} \tag{4}$$

where $\phi(s)$ gives the angle between the local tangent vector to the meridian and the radially outward direction. The Lagrange multipliers $f_r$ and $f_z$ can be interpreted as the radial and vertical components, respectively, of the force exerted on a curve of constant $z$ due to membrane stress. With this parametrization, the mean curvature can be written

$$C = \frac{1}{2}\left(\phi'(s) + \frac{\sin(\phi(s))}{r(s)}\right). \tag{5}$$

Inserting this expression for the mean curvature into the membrane energy and taking the functional derivative of the membrane energy with respect to $r(s), z(s), \phi(s), f_r(s)$, and $f_z(s)$, we find a set of differential equations characterizing fusion pore shapes that minimize the membrane energy,

$$\begin{gathered} r'(s) = \cos(\phi(s)) \\ z'(s) = \sin(\phi(s)) \\ \phi'(s) = 2C - \frac{\sin(\phi(s))}{r(s)} \\ C'(s) = \frac{f_r(s)\sin(\phi(s)) - f_z(s)\cos(\phi(s))}{2\kappa r(s)} \\ f_r'(s) = \gamma + 2\kappa C\left(C - \frac{\sin(\phi(s))}{r(s)}\right) \\ f_z'(s) = 0. \end{gathered} \tag{6}$$

These are the Hamilton's equations corresponding to the Lagrangian given by $U_{\mathrm{mb}}$, and are equivalent to the membrane shape equation. The last equation indicates that $f_z$ is a constant; this is associated with the assumed symmetry of the fusion pore. We therefore have five first order differential equations, and two unknown parameters ($f_z$ and $L$), requiring seven boundary conditions.

At the end of the meridian corresponding to the perimeter of the NLP (defined as $s = 0$), the boundary conditions are

$$\begin{gathered} r(0) = R_{\mathrm{NLP}}, \\ 2\kappa C(0) = \; k \sin 2\phi(0), \end{gathered} \tag{7}$$

where $R_{\mathrm{NLP}}$ is the radius of the NLP, and $k$ is the twisting stiffness of the ApoE scaffold. The latter equation guarantees torque equilibrium between the membrane and the NLP scaffold, described in the SI subsection *"Mathematical Model of the ApoE scaffold"*. At the opposite end of the meridian, where the fusion pore meets the tCell membrane, the boundary conditions are

$$
\begin{aligned}
r(L) &= R_\infty, \\
z(L) &= 0, \\
\phi(L) &= 0.
\end{aligned} \tag{8}
$$

where $R_\infty = 30$ nm is chosen to be much larger than the length scale of the fusion pore. The choice $z(L) = 0$ is arbitrary, and is only used to set a reference point for the $z$ coordinate.
We fix the vertical force on the fusion pore $f_z$ in order to vary the height of the pore. The membrane energy is minimized when $f_z = 0$. However, forces created by the Syt-SNARE complex can alter the height of the fusion pore. We therefore scan the parameter $f_z$ from $-6$ pN to $+6$ pN to find fusion pores that satisfy the geometric constraint imposed by the Syt-SNARE complex. We assume that the selected height is that which minimizes the free energy subject to the constraints of the Syt-SNARE complex. Lastly, the contour length of the meridian $L$ is unknown. The selected contour length is the one that minimizes the membrane energy. The condition that $L$ minimizes the membrane energy is given by (21)

$$
0 = \cos\big(\phi(L)\big)\, f_r(L) + \sin\big(\phi(L)\big)\, f_z + r(L)\left(2\kappa\, C(L)\left[C(L) - \frac{\sin\big(\phi(L)\big)}{r(L)}\right] - \gamma\right). \tag{9}
$$

In order to find the shape of a fusion pore with a given radius $R_{\text{pore}}$, we additionally impose boundary conditions

$$
\begin{aligned}
r(x) &= R_{\text{pore}} \\
\phi(x) &= \frac{\pi}{2}
\end{aligned} \tag{10}
$$

at the waist of the fusion pore, between the NLP and the tCell membrane. Note that this is measured from the membrane midplane, and related to the pore radius by $R_{\text{pore}} = r_{\text{po}} + \delta/2$, where $\delta$ is the membrane thickness. Since the location of the waist is not known in advance, this procedure is mathematically equivalent to solving the shape equation twice, once from the NLP to the waist, and once from the waist to the point where the fusion pore joins the tCell membrane.

**Mathematical Model of the ApoE scaffold**

We modelled the ApoE scaffold of the NLPs as an elastic rod with rectangular cross-section, a simple representation of the two parallel alpha helices comprising the scaffold. The elastic bending energy of the rod is

$$
U_{\text{scaffold}} = \int dL\left(\frac{K_{\text{soft}}}{2}\, C_{\text{soft}}^2 + \frac{K_{\text{hard}}}{2} C_{\text{hard}}^2\right), \tag{11}
$$

where $K_{\text{soft}}$ and $K_{\text{hard}}$ are the bending moduli of the rod in the material directions across its narrow and wide faces, respectively, $C_{\text{soft}}$ and $C_{\text{hard}}$ are the respective material curvatures, and $L$ measures the arclength around the scaffold. The material curvatures represent the curvature of the scaffold projected onto the basis vectors of the material cross section. Because of the rectangular shape of the cross section, the rod is more difficult to bend across its wide face than across its narrower face. For a homogeneous scaffold with a rectangular cross-section of width $w$ (across the wider face) and thickness $t$ (across the narrower face), this is quantified by a classical result from elasticity theory, which states $K_{\text{soft}} = E\, t^3\, w/12,\ K_{\text{hard}} = E\, t\, w^3/12$, where $E$ is the Young's modulus of the scaffold (22). From these scaling relations, we infer that the scaffold should have twice the bending modulus of a single alpha helix in the soft direction, and 8 times the bending modulus of a single alpha helix in the hard direction. Given the typical persistence length of an alpha helix of ~100 nm and the well-known relation between bending modulus and persistence length $K = k_{\mathrm{B}}T\, L_{\mathrm{p}}$ (23), we conclude $K_{\mathrm{soft}} \approx 200\ kT$ nm and $K_{\mathrm{hard}} \approx 800\ kT$ nm.

Suppose the cross section of the scaffold is rotated such that the long axis of the cross section makes an angle $\phi$ with the vertical direction (as in Fig. S8A), while maintaining the shape of the scaffold as a ring of radius $R_{\mathrm{NLP}} = 12$ nm. Then, the material curvatures are

$$C_1 = \frac{\cos\phi}{R_{\mathrm{NLP}}} \quad C_2 = \frac{\sin\phi}{R_{\mathrm{NLP}}}, \tag{12}$$

This gives an elastic bending energy

$$U_{\mathrm{scaffold}} = 2\pi R_{\mathrm{NLP}}\left(\frac{K_{\mathrm{soft}}}{2}\frac{\cos^2\phi}{R_{\mathrm{NLP}}^2} + \frac{K_{\mathrm{hard}}}{2}\frac{\sin^2\phi}{R_{\mathrm{NLP}}^2}\right). \tag{13}$$

The torque per unit length to twist the scaffold through an angle $\phi$ is thus

$$\tau = \frac{1}{2\pi R_{\mathrm{NLP}}}\frac{\partial F}{\partial\phi} = \frac{K_{\mathrm{hard}} - K_{\mathrm{soft}}}{2R_{\mathrm{NLP}}^2}\sin 2\phi = k\ \sin 2\phi, \tag{14}$$

where $k$ is the apparent twisting rigidity of the scaffold. Using the parameter values above, we find $k = 4.2\ k_BT/\mathrm{nm}$. The model predicts that a torque per unit length ~2 pN twists the rod ~30 degrees.

We incorporated these effects in our calculation of the fusion pore shape by imposing a torque equilibrium condition at the NLP edge, so that membrane torque is resisted by the ND scaffold. The membrane bending torque per unit length about the local tangent vector to the NLP boundary is given by $\tau = 2\kappa C$, where $C$ is the local mean curvature, neglecting terms associated with the Gaussian curvature modulus (24). Thus, equilibrium is attained when

$$2\kappa C = k\sin 2\phi, \tag{15}$$

at the boundary of the NLP.

**Geometric constraints imposed by the Syt-SNARE complex**

In order to determine whether a fusion pore determined by solving the membrane shape equation satisfied the constraints imposed by the Syt-SNARE complex, we directly compared the geometry of the fusion pore with that of the Syt-SNARE complex. First, we measured the dimensions of the Syt-SNARE complex and the orientation of the SNARE complex relative to the membrane using PyMol. The long axis of the SNARE formed a ~15° angle with the membrane when docked via the primary interface to the Syt C2B domain in the orientation measured using electron paramagnetic resonance (17, 20). The point where the SNARE complex would contact the membrane (determined by the Syntaxin and VAMP TMDs) was located $d = 8.0$ nm from the point where the C2B domain inserted into the PM, and the line connecting the two contact points made a $\psi = 39°$ angle with the PM.

To determine if the fusion pore satisfied the relevant constraints, we represented the Syt-SNARE complex as a wedge, with one segment of length 10 nm approximately perpendicular to the membrane, representing the long axis of the SNARE complex. Another segment of length 8 nm at a 17.7° angle to the first represented the line between the two points where the Syt-SNARE complex inserts into the membrane. We scanned this wedge along the meridian of the fusion pore seeking two points $\boldsymbol{r}_1$ and $\boldsymbol{r}_2$ representing the membrane contact points of the Syt-SNARE complex satisfying the following criteria:

1. The distance between $\boldsymbol{r}_1$ and $\boldsymbol{r}_2$ is within 10% of the measured distance between the membrane contact points, $\frac{|r_1 - r_2|}{d} < 0.1$. This 10% error range is roughly comparable with the depth of insertion of the C2B domain Ca-binding loops beyond the phosphate plane (18, 20, 25).
2. The line connecting $\boldsymbol{r}_1$ and $\boldsymbol{r}_2$ makes an angle with the membrane tangent vector at $\boldsymbol{r}_2$ within 0.1 radians of the measured value, $\psi - 0.1 < \arccos\left(\frac{r_1 - r_2}{|r_1 - r_2|} \cdot \boldsymbol{t}_2\right) < \psi + 0.1$, where $\boldsymbol{t}_2$ is the

membrane tangent vector (pointing along the meridian) at $\boldsymbol{r}_2$, representing the point where Syt contacts the membrane.

3. The long axis of the SNARE complex makes an angle with the membrane normal vector at $\boldsymbol{r}_1$ of less than 0.1 radians.

If all three conditions were satisfied, the pore was assumed to satisfy the constraint imposed by the Syt-SNARE complex.

**Short-ranged steric hydration free energy**

The pressure due to short-ranged hydration forces between membranes with separation $d$ follows the form $P_0 \exp(-d/\lambda)$, where $\lambda$ is the characteristic length scale over which the hydration forces decay and $P_0$ is a pressure pre-factor (26). The steric hydration free energy was evaluated in our previous work by calculating the work done by the hydration pressure to increase the pore size of a toroidal pore (4). The expression for the hydration free energy is given by

$$U_{\text{hyd}} = P_0 \lambda \pi l \exp\left(-\frac{2r_{\text{po}}}{\lambda}\right)\left(r_{\text{po}} + \frac{\lambda}{2}\right), \quad (16)$$

where $l = \sqrt{2\lambda(h + 2\delta)}$ is the effective pore height that substantially contributes to the steric hydration interaction. For purposes of determining the hydration energy, we used this expression, approximating the fusion pore as a toroid. Another term giving the work done by the hydration forces to bring two distant planar membranes to a separation $h$ was omitted because it contributed negligibly. The second term is the work done to separate the membranes to form a pore of radius $r_{\text{po}}$. Values for $P_0$ and $\lambda$ are obtained from previous studies and are set to $P_0 = 5 \times 10^{11}$ dyn/cm$^2$ and $\lambda = 0.1$ nm (see Table S1).

**Free energy of SNAREs**

We assume that each side of the NLP contains $N$ v-SNAREs that are all available to associate with the t-SNAREs in the tCell membrane and contribute to pore expansion. SNAREs can be fully zippered, where their TMDs are circularly arranged near the fusion pore waist. Alternatively, they can adopt a partially zippered configuration, where the TMD and linker domain are unzippered, and the v-SNARE and t-SNARE TMDs are located on the NLP and tCell membranes, respectively, but on the same side of the fusion pore. We denote the number of fully and partially zippered as $N_{\text{Z}}$ and $N_{\text{UZ}}$, respectively. The SNARE free energy in the fully zippered state reads

$$U_{\text{Z}}(r_{\text{po}}, N_{\text{Z}}) = -N_{\text{Z}} k_{\text{B}} T \left[\ln \frac{2\pi r_{\text{po}} - N_{\text{Z}} b}{N_{\text{Z}} b} + 1\right] - N_{\text{Z}} k_{\text{B}} T \ln \Omega_{\text{Z}} - N_{\text{Z}} \epsilon_{\text{Z}}, \quad (17)$$

where $k_{\text{B}}$ is the Boltzmann constant, $T$ is the temperature and $b = 2$ nm is the thickness of a single SNARE (4). The first term in eq. (**Error! Reference source not found.**7) is the positional entropy of the zippered SNAREs TMDs. The second term is the orientational entropy associated with the zippered SNAREs. We assume that these are stiff rods that can explore a small solid angle of $\Omega_{\text{Z}} = 0.05$ sr, based on molecular dynamics studies of t-SNARE TMDs showing that these domains explore angles of 10° around their equilibrium position (27). We assume that for the zippered SNAREs the equilibrium orientation is the local normal to the fusion pore membrane.

The last term is the total energy released when the TMDs and the adjacent linker regions of $N_{\text{Z}}$ SNAREs are fully zippered, where $\epsilon_{\text{Z}}$ is the zippering energy per SNARE. This zippering energy was obtained as a best-fit parameter in a previous study where fusion was induced between NLPs and tCells with only SNAREs (4). The best-fit value of 9.6 $k_{\text{B}}T$ (Table S1) is higher than the ~5 $k_{\text{B}}T$ that we estimated from a previous study as the zippering energy of the linker domains (28), as explained in the following paragraph.

The linker domain (LD) has ~10 residues (29), and is thus ~ 3 nm in length, assuming an unfolded contour length 0.3 nm per residue. Previous measurements show the free energy to unzip the SNAREs has slope ~1.5 $k_BT$ per nm when the LDs are being unzippered (28). Thus, we estimate the LD unzippering energy from Gao et al. is ~ 5 $k_BT$.

The SNARE free energy in the partially zippered state reads

$$U_{UZ}(r_{po}, N_{UZ}) = -N_{UZ} k_B T \ln \frac{2\pi D}{b} - N_{UZ} k_B T \ln \Omega_{UZ}. \tag{18}$$

The first term in eq. (**Error! Reference source not found.**8) is the positional entropy of the TMDs, while the second term is the orientational entropy associated with a solid angle $\Omega_{UZ}$ explored by the SNAREs. In the partially zippered state the SNARE linker domains are assumed to be unstructured, which allows them to adopt all orientations where they are not intersecting with the membranes. Since this orientational freedom is available when the SNAREs are away from the fusion pore, we restrict their position to the edge of the pore and set $\Omega = \Omega_{UZ} = \pi$.

We assume that, in elevated Ca concentrations, the C2B domains will bind the membrane via their Ca-binding loops. This lifts the C-terminal end of the SNARE complex ~5 nm above the tCell membrane; this has been proposed to drive dissociation of SNARE complexes from Syt in the presence of Ca (30, 31). As this dissociation would cost $10 - 12\ k_B T$ (see subsection *Pushing forces from the membranes are insufficient to disrupt the SNARE-Syt primary-interface interaction*), we omit this possibility and assume that SNAREs are unable to explore the fusion pore in elevated Ca concentrations. In this case, the positional entropy of SNAREs is unaltered by unzippering, and the free energy difference between the zippered and unzippered state is therefore given by $\epsilon_Z$ per zippered SNARE.

**Total free energy as a function of pore size, minimum membrane separation, and total number of SNAREs**

To obtain the free energy of a fusion pore with a radius $r_{po}$, we numerically summed all the Boltzmann factors of all possible states according to

$$\exp\left(-\frac{U_{tot}(r_{po}, h, N)}{k_B T}\right) = \sum_{N_Z=0}^{N} \exp\left(-\frac{U_{tot}(r_{po}, h, N, N_Z)}{k_B T}\right) \tag{19}$$

$$U_{tot}(r_{po}, N) = \min_{h}\{U_{tot}(r_{po}, h, N)\}$$

where we set the number of SNAREs to $N = 4$ to match experiment.

**Model-predicted pore conductance**

Consider an axially symmetric fusion pore whose inner surface is described by a function $r(z)$ which gives the distance from the axis to the luminal surface of the membrane. The resistance of the pore is given by

$$R_{po} = \rho \int_0^L \frac{dz}{\pi\, r(z)^2} \tag{20}$$

where ρ is the resistivity of the solution in the pore lumen, and the height of the pore lumen $L = h + 2\delta$, where δ is the thickness of the bilayer (32). We numerically evaluated the above integral for fusion pores determined by solving the membrane shape equation. The total resistance of the pore also has a contribution from access resistance given by (32)

$$\mathrm{R}_{\mathrm{acc}} = \frac{\rho}{2r_{\mathrm{po}} + L}. \tag{21}$$

The pore conductance is then given by $G_{\mathrm{po}} = \left(\mathrm{R}_{\mathrm{po}} + \mathrm{R}_{\mathrm{acc}}\right)^{-1}$.

**Pushing forces from the membranes are insufficient to disrupt the SNARE-Syt primary-interface interaction**

We show here that forces needed to separate the membranes and expand the fusion pore are not large enough to disrupt the SNARE-Syt primary interface.

To calculate the pushing forces shouldered by the SNARE-Syt complexes, we calculated increase in the pore free energy $U_{\mathrm{mb}}$ caused by the expansion driven by the SNARE-Syt complexes as a function of pore radius (Fig. S8D). From Fig. S8D, we see that change in height costs ~0-10 $k_BT$. The forces increasing the height of the pore are $\partial U_{\mathrm{mb}} / \partial h \approx 10$ pN. This force is shared across 4 SNARE-Syt complexes. Thus, an estimate for the force shouldered per complex during pore expansion is ~2.5 pN.

We estimate that 2.5 pN is far less than the force needed to break the primary C2B-SNARE complex interface. Reported dissociation constants for the SNARE-C2B complex are 0.86 μM(35) and 14 μM(36). These correspond to binding energies $\Delta G$ of 12 $k_BT$ and 10 $k_BT$, respectively, after estimating a microscopic capture radius of 2 nm (equivalently a reference concentration 0.21M) and using $\Delta G = -k_B T \ln(K_d/0.21\mathrm{M})$. Even if we conservatively use a large "unbinding distance" $d$ ~ 3 nm, breaking the interface thus requires a force of order $\Delta G/d$ ~14-16 pN, much larger than the ~4 pN force exerted on a lever complex by the fusion pore. Thus, we expect the SNARE-Syt complexes will remain intact.

### SUPPLEMENTARY TABLE 1

| Symbol | Meaning | Value | Legend |
|---|---|---|---|
| $\boldsymbol{\delta}$ | Membrane thickness | 5 nm | (A) |
| $\boldsymbol{\epsilon_Z}$ | Zippering energy of SNARE's linker and TMD domains | 9.6 $k_B T$ | (B) |
| $\boldsymbol{D}$ | NLP diameter | 24 nm | (C) |
| $\boldsymbol{\lambda}$ | Hydration interactions decay length | 0.1 nm | (B) |
| $\boldsymbol{P_0}$ | Pressure pre-factor for steric hydration interaction | $5 \times 10^{11}$ dyn/cm$^2$ | (D) |
| $\boldsymbol{\kappa}$ | Membrane bending modulus | 20 $k_{\mathrm{B}} T$ | (E) |
| $\boldsymbol{\gamma}$ | Membrane tension | 0.1 pN $\cdot$ nm$^{-1}$ | (F) |
| $\boldsymbol{\Omega_Z}$ | Solid angle explored by fully zippered SNAREs | 0.05 sr | (G) |
| $\boldsymbol{k}$ | Twisting rigidity of ApoE scaffold | 600 $k_{\mathrm{B}} T \cdot$ nm | (H) |
| $[\mathbf{Ca^{2+}}]_{1/2}$ | Apparent affinity of Syt1 to calcium in the presence of PIP2 containing membranes | 23 μM | (I) |
| $\boldsymbol{n}$ | Hill coefficient | 2.3 | (I) |

**Table S1**. Parameters used in the mathematical model with coarse-grained membranes, SNAREs and Synaptotagmin-1 C2B domains. (A) Measured in (37). (B) Estimated previously as a best fit model parameter to experiments where NLP-tCell fusion pore was induced only by SNAREs (4). (C) Consistent with NLP diameter measurements in this study. (D) Calculated as the weighted average of the hydration pressures of palmitoyl-2-oleoyl phosphatidylcholine (POPC) and 1,2-dioleoyl phosphatidylserine (DOPS) adopted from (26), using a (85:15) molar ratio of POPC:DOPS as present in the NLP in the current study. (E) Values for the bending modulus range between $10 - 50\ k_{\mathrm{B}} T$ (38-41). We used a value of $\kappa = 20\ k_{\mathrm{B}} T$. (F) Obtained by fitting the membrane energy as a function of pore radius to measurements from a previous study using a similar method to measure the pore free energy in the absence of SNAREs, see Fig. S8B (42). (G) Calculated based on a $10°$ angle explored by t-SNARE TMDs around the equilibrium configuration, as measured in molecular dynamics simulations (27). (H) Estimated here from the $\sim$100 nm persistence length of typical alpha helices and cross-sectional dimensions of the ApoE scaffold (see SI subsection *"Mathematical Model of the ApoE scaffold"*). (I) Measured in Syt1-liposome binding assays (33, 43)

### SUPPLEMENTARY FIGURE LEGENDS

**Figure S1.** Co-reconstitution of Synaptotagmin-1 and VAMP2 into nanolipoprotein particles (vsNLPs). **A**. Schematic of an NLP reconstituted with 4 copies per face each with Syt1 and VAMP2. Syt1 C2AB domains are shown in green; VAMP2 is shown in blue and the scaffold protein ApoE protein is shown in cyan. Domain structures of Syt1 and VAMP2 are indicated. **B**. Typical size-exclusion chromatography elution profiles of vNLP (NLPs loaded with VAMP2 alone) and vsNLP (NLPs loaded with VAMP2 and Syt1) samples using a Superose 6 Increase 10/300 GL column. Proteins were detected using absorbance at 280 nm. Collected eluted volumes are indicated by horizontal bars above each profile. **C**. SDS-PAGE stained with Coomassie Brilliant Blue shows the purified NLPs carried Syt1 and VAMP2 proteins. **D**. A representative transmission electron microscopy (TEM) image of a vsNLP sample after purification. Nanodiscs indicated by white stars have their lipid bilayer plane positioned perpendicularly to the imaging plane. **E**. Distribution of vsNLP sizes from TEM images. More dilute samples (5-10x) than the example shown in D were used for size quantification, such that most NLPs were lying flat on the grid. A Gaussian fit to the distribution is shown as the red solid line (fitted mean diameter = 25±5.6 nm (± SD), n

= 200 NLPs). **F-H.** Characterization of nanolipoprotein particles reconstituted with VAMP2 alone. **F**. SDS-PAGE of purified vNLPs stained with Coomassie Brilliant Blue indicting the NLPs incorporated ApoE and VAMP2 proteins. **G**. Negative stain transmission electron microscopy image of a representative vNLP sample. Nanodiscs indicated by white stars are oriented with their disc plane perpendicular to the imaging plane. **H**. Size distribution of vNLPs reconstituted with a total of 8 copies of VAMP2 per disc. A Gaussian fit is shown as the red solid line (best fit diameter =25 ± 4.6 nm (mean ± SD), 171 discs were analyzed). **I**. Domain structure of flipped t-SNARE constructs used to generate HeLa cells stably expressing flipped t-SNAREs (7, 8). **J**. Example of a fusion pore current burst and definition of analysis parameters. To be included in the analysis, a current burst must have amplitude >2 pA and last at least 250 ms. Open sub-periods during a burst cross a threshold (-0.25 pA, red dotted line) for at least 60 ms (indicated as the thick colored bars above the current trace). The number of open sub-periods during a burst is equal to the number of pore flickers, $N_{flickers}$. The duration of the burst is the time from the first detected open-pore point until the last one and is denoted $T_o$. The flicker rate is $N_{flickers}/T_o$. The pore open probability is the sum of the open-pore sub-periods (the colored bars) divided by the burst lifetime, $T_o$.

**Figure S2**. Fusion rates. **A**. An example of an entire 800 s recording that started shortly after establishing a tight seal. The colored regions correspond to currents that are counted as fusion pore currents because they fit the criteria explained in SI Appendix *Detection of pore currents* and *Estimation of fusion rate* (also see Fig. S1J). These regions are shown with expanded axes in insets. In this example, 3 pores were counted in 800 s of recording. **B**. Another example of an 800 s recording. In this case, large currents appeared starting ~140 s. The baseline did not recover before the end of the recording and the red-colored portion was excluded from analysis. Thus, two current bursts contributed to the fusion rate from this trace (starting ~90 and 130 s, colored in green and teal), from 140 s of recording. **C**. Fusion rate increases with increasing NLP concentration. Fusion rates were calculated as described in SI Appendix *Estimation of fusion rate* and plotted against the concentration of NLPs reconstituted with v-SNAREs (8 copies total). Pores from 12-37 cells were recorded for every condition. Error bars indicate S.E.M. The best fit straight line is shown (slope $= 2.3 \times 10^{-3}$ pores/(min $\cdot$ nM), $R^2 = 0.86$). **D.** Comparison of per cell and overall fusion rates. As an alternative estimate of the fusion rate, we summed all detected pores, $N_{tot}$, and the analysis time $\tau_{tot}$ over all cells (excluding portions with noisy/unstable baseline), and calculated the total number of pores divided by the total analysis time, $\dot{F}_{tot} = N_{tot}/\tau_{tot}$ for the indicated conditions. This estimate (blue) is compared with the per cell estimate used throughout (red).

**Figure S3.** Additional properties of single fusion pores in the presence of full-length Syt1 or soluble C2AB. Open-pore conductance fluctuations relative to mean (**A**), average flicker rate during a burst (**B**), average open-pore probability, $P_o$, during a current burst (fraction of time pore is in the open state during a burst) (**C**), and average burst lifetime, $T_o$, (**D**) for the indicated conditions. **E**. Distributions of the number of flickers per burst, $N_{flickers}$, for the indicated conditions. Fits to geometric distributions are shown in red, $y = p(1-p)^{n-1},\ n = 1, 2, 3, \ldots$ . Best fit parameters (with $\pm$ 95% confidence intervals) are $p =$ 0.072 (0.053, 0.092) (no Syt1, 100 μM $Ca^{2+}$, averaged over 49 individual fusion pores from 10 cells, mean $N_{flickers} = 12.8$), 0.083 (0.051, 0115) (Syt1, 0 μM $Ca^{2+}$; averaged over 24 individual fusion pores from 11 cells, mean $N_{flickers} = 11.0$), 0.053 (0.044, 0.063) (Syt1, 100 μM $Ca^{2+}$; averaged over 123 individual fusion pores from 20 cells, mean $N_{flickers} = 17.7$). **F**. Distribution of burst lifetimes, $T_o$ for the indicated conditions. Best fits to single exponentials are shown as red curves, with means (and 95% confidence intervals) as follows. No Syt1, 100 μM $Ca^{2+}$: 6.1 s (4.7 to 8.3 s, 49 fusion pores from 10 cells), Syt1, 0 μM $Ca^{2+}$: 6.5 s (4.5 to 10.1 s, 24 fusion pores from 11 cells), Syt1, 100 μM $Ca^{2+}$: 16 s (13.5 to 19.3 s, 123 fusion pores from 20 cells). In A-D, the two-sample Kolmogorov-Smirnov test was used to assess significant differences between the "no C2AB" group and the rest. *, **, *** indicate p<0.05, 0.01, and 0.001, respectively. Comparison between Syt1 and C2AB in the presence of $Ca^{2+}$ and $PI(4,5)P_2$ are also indicated (using the two-sample Kolmogorov-Smirnov test).

**Figure S4.** Syt1 C2AB dilates fusion pores in a calcium and $PI(4,5)P_2$ dependent manner. **A-D**. Probability density function (PDF) for point-by-point open-pore conductance values for the indicated conditions. Substantial density is present for $G_{po} \gtrsim 500$ pS only when C2AB, calcium, and $PI(4,5)P_2$ were all present. **F-I**. PDFs for open-pore radii corresponding to the conductance distributions in A-D, assuming pores are 15 nm long cylinders. Data were from 49 fusion pores/10 cells (SNARE only), 44 fusion pores/12 cells (0 μM $Ca^{2+}$), 84 fusion pores/19 cells (no $PI(4,5)P_2$) and 98 fusion pores/17 cells (100 μM $Ca^{2+}$ plus $PI(4,5)P_2$).

**Figure S5.** Additional fusion pore properties for Syt1 C2AB domains carrying mutations in D309, K326-327 and R398-399. **A**. Open-pore conductance fluctuations relative to mean. Compared with the SNARE-alone (no C2AB) group, fluctuations were larger for wild-type C2AB, and lower for $C2AB^{K326A,327A}$. **B**. Average flicker rate for the same conditions as in A. Compared with the SNARE alone group (no C2AB), C2AB and $C2AB^{R398Q,R399Q}$ decreased the flicker rate. **C**. Average pore open probability during a burst, $P_o$, for the indicated conditions. Compared with the SNARE alone group (no C2AB), C2AB and $C2AB^{K326A,K327A}$ had larger pore open probabilities. **D**. Average burst lifetimes for the same conditions. Error bars are +/- S.E.M. Data were from no C2AB: 49 pores/10 cells, C2AB: 98 pores/17 cells, $C2AB^{K326A,327A}$: 42 pores/14 cells, $C2AB^{D309N}$ (18 pores/7 cells), $C2AB^{R398Q,R399Q}$ (42 pores/18 cells). For A-D, two-sample Kolmogorov-Smirnov test was used to assess significant differences between the "no C2AB" group and the rest. *, **, *** indicate p<0.05, 0.01, and 0.001, respectively.

**Figure S6.** Additional fusion pore properties as a function of free calcium concentration, $[Ca^{2+}]_{free}$. **A-D**. Average single-pore conductance fluctuations relative to mean (**A**), average burst lifetime (**B**), average flicker rate (**C**), and the pore open probability $P_o$ (**D**) as a function of $[Ca^{2+}]_{free}$. Error bars are +/- S.E.M. **E**. Probability density functions (PDFs) for point-by-point open-pore conductance values at different $[Ca^{2+}]_{free}$. The probability density for $G_{po} \gtrsim 500$ pS increases as a function of calcium. (0 μM $Ca^{2+}$: 44 pores/12 cells; 5 μM $Ca^{2+}$: 54 pores/20 cells; 20 μM $Ca^{2+}$: 114 pores/18 cells; 50 μM $Ca^{2+}$: 88 pores/26 cells; 100 μM $Ca^{2+}$: 98 pores/17 cells). **F.** The fusion rate increases as a function of $[Ca^{2+}]_{free}$. For A-D, two-sample Kolmogorov-Smirnov test was used to assess significant differences between the "no C2AB" group and the rest. *, **, *** indicate p<0.05, 0.01, and 0.001, respectively.

**Figure S7.** Additional fusion pore properties for Syt1 C2AB membrane penetration mutants. **A-D**. Average single-pore conductance fluctuations relative to mean (**A**), burst lifetime (**B**), pore open probability during a burst, $P_o$ (**C**), and flicker rate (**D**) for SNAREs alone (no C2AB), wild-type Syt1 C2AB (C2AB), the 4W mutant with enhanced membrane-penetration ability (M173W, F234W, V304W and I367W), and the 4A mutant which cannot penetrate membranes in response to calcium (M173A, F234A, V304A and I367A). Error bars are +/- S.E.M. **E**. Probability density functions (PDFs) for point-by-point open-pore conductance values for wild-type C2AB, and the membrane penetration mutants 4A and 4W. (4W: 115 pores/20 cells, 4A: 31 pores/9 cells). For A-D, two-sample Kolmogorov-Smirnov test was used to assess significant differences between the "no C2AB" group and the rest. *, **, *** indicate p<0.05, 0.01, and 0.001, respectively

**Figure S8**. Results of the mathematical model of the fusion in the presence of SNAREs and Syt1 C2AB domains. Data in (C) and (D) was smoothed using a moving average with width 0.5 nm. (**A**) Schematic illustrating a fusion pore with buried SNARE-Syt1 levers. ϕ represents the angle of twisting of the ApoE proteins. (**B**) Pore free energy as a function of radius predicted by the model and measured in a previous study (42); the membrane tension was tuned to reproduce the experimental curve here. **(C)** Pore height, defined as the maximal separation between the NLP and the tCell membranes, as a function of pore radius with and without $Ca^{2+}$. (**D**) Free energy difference between pores in the expanded state and those in the unexpanded state. Syt-SNARE complex-driven pore expansion costs a maximum of $\sim 10 - 12\ k_{\mathrm{B}}T$ at low pore radii.

**Figure S9**. Lack of ATP-regulated channel activity in flipped t-SNARE cells. **Top:** Schematic of the cell-attached recordings to test for ATP-regulated channel activity. Pipette solutions were the same as for single-pore measurements with 100 µM free calcium, but adjusted to contain either 0 or 2 mM ATP. Pipette potential was -40 mV. **Middle**: Current recordings under voltage clamp from three different patches (out of 21 total) in the absence of ATP. **Bottom**: Current recordings as in the middle panels, but in the presence of 2 mM freshly prepared ATP (26 cells were recorded). Occasionally, channel activity is recorded both with and without ATP.

**Figure S10.** Average pore conductance as a function of time, after aligning pores to the moment of opening (blue) or closure (red). Data for full-length Syt1, in the presence of 100 µM free $Ca^{2+}$ and $PI(4,5)P_2$. Other conditions also failed to yield large differences between pore opening or closure.

**Figure S11.** Whole-cell conductance of flipped t-SNARE HeLa cells. **A**. Whole-cell current responses to step changes in membrane potential under voltage-clamp, from a HeLa cell line expressing flipped t-SNAREs. **B**. Current-voltage relationship. Currents were averaged for 27 cells. The average slope is $G_{cell} = 5.04 \pm 0.32$ nS (95% confidence interval), excluding the five highest voltages (red crosses). If all points are included, $G_{cell} = 6.20 \pm 0.51$ nS (95 % confidence interval). The range of mean open-pore currents and transmembrane voltages comprising 95% of the data values for C2AB in the presence of $PI(4,5)P_2$ and 100 µM calcium are indicated as a red-colored box.

#### REFERENCES


1. J. A. Morrow, K. S. Arnold, K. H. Weisgraber, Functional characterization of apolipoprotein E isoforms overexpressed in Escherichia coli. *Protein Expr Purif* **16**, 224-230 (1999).
2. C. D. Blanchette *et al.*, Quantifying size distributions of nanolipoprotein particles with single-particle analysis and molecular dynamic simulations. *Journal of lipid research* **49**, 1420-1430 (2008).
3. Z. Wu *et al.*, Nanodisc-cell fusion: control of fusion pore nucleation and lifetimes by SNARE protein transmembrane domains. *Sci Rep* **6**, 27287 (2016).
4. Z. Wu *et al.*, Dilation of fusion pores by crowding of SNARE proteins. *Elife* **6**, e22964 (2017).
5. L. Ma *et al.*, Single-molecule force spectroscopy of protein-membrane interactions. *Elife* **6**, e30493 (2017).
6. O. D. Bello, S. M. Auclair, J. E. Rothman, S. S. Krishnakumar, Using ApoE Nanolipoprotein Particles To Analyze SNARE-Induced Fusion Pores. *Langmuir* **32**, 3015-3023 (2016).
7. C. G. Giraudo, W. S. Eng, T. J. Melia, J. E. Rothman, A clamping mechanism involved in SNARE-dependent exocytosis. *Science* **313**, 676-680 (2006).
8. C. G. Giraudo *et al.*, SNAREs can promote complete fusion and hemifusion as alternative outcomes. *J Cell Biol* **170**, 249-260 (2005).
9. U. Baron, S. Freundlieb, M. Gossen, H. Bujard, Co-regulation of two gene activities by tetracycline via a bidirectional promoter. *Nucleic Acids Res* **23**, 3605-3606 (1995).
10. N. R. Dudzinski, Z. Wu, E. Karatekin, A Nanodisc-Cell Fusion Assay with Single-Pore Sensitivity and Sub-millisecond Time Resolution. *Methods Mol Biol* **1860**, 263-275 (2019).
11. B. Hille, *Ion channels of excitable membranes* (Sinauer, Sunderland, Mass., ed. 3rd, 2001), pp. xviii, 814 p.
12. L. Welter-Stahl *et al.*, Expression of purinergic receptors and modulation of P2X7 function by the inflammatory cytokine IFNgamma in human epithelial cells. *Biochim Biophys Acta* **1788**, 1176-1187 (2009).
13. E. M. Fenwick, A. Marty, E. Neher, A patch-clamp study of bovine chromaffin cells and of their sensitivity to acetylcholine. *J Physiol* **331**, 577-597 (1982).
14. R. Fischmeister, R. K. Ayer, Jr., R. L. DeHaan, Some limitations of the cell-attached patch clamp technique: a two-electrode analysis. *Pflugers Arch* **406**, 73-82 (1986).
15. O. P. Hamill, "Potassium and Chloride Channels in Red Blood Cells" in Single-Channel Recording, B. Sakmann, E. Neher, Eds. (Plenum Press, New York, 1983), pp. 451-472.
16. W. Helfrich, Elastic properties of lipid bilayers: theory and possible experiments. *Z Naturforsch C* **28**, 693-703 (1973).
17. Q. Zhou *et al.*, Architecture of the synaptotagmin-SNARE machinery for neuronal exocytosis. *Nature* **525**, 62-67 (2015).
18. W. Kuo, D. Z. Herrick, J. F. Ellena, D. S. Cafiso, The calcium-dependent and calcium-independent membrane binding of synaptotagmin 1: two modes of C2B binding. *J Mol Biol* **387**, 284-294 (2009).
19. D. Z. Herrick, S. Sterbling, K. A. Rasch, A. Hinderliter, D. S. Cafiso, Position of synaptotagmin I at the membrane interface: cooperative interactions of tandem C2 domains. *Biochemistry* **45**, 9668-9674 (2006).
20. W. Kuo, D. Z. Herrick, D. S. Cafiso, Phosphatidylinositol 4,5-bisphosphate alters synaptotagmin 1 membrane docking and drives opposing bilayers closer together. *Biochemistry* **50**, 2633-2641 (2011).
21. F. Julicher, U. Seifert, Shape equations for axisymmetric vesicles: A clarification. *Phys Rev E Stat Phys Plasmas Fluids Relat Interdiscip Topics* **49**, 4728-4731 (1994).
22. L. D. Landau, E. M. Lifshitz, in Theory of Elasticity (Third Edition), E. M. Lifshitz, A. M. Kosevich, L. P. Pitaevskii, Eds. (Butterworth-Heinemann, Oxford, 1986),

https://doi.org/10.1016/B978-0-08-057069-3.50009-7 chap. CHAPTER II - THE EQUILIBRIUM OF RODS AND PLATES, pp. 38-86.
23. S. Choe, S. X. Sun, The elasticity of alpha-helices. *J Chem Phys* **122**, 244912 (2005).
24. M. Deserno, Fluid lipid membranes: from differential geometry to curvature stresses. *Chem Phys Lipids* **185**, 11-45 (2015).
25. A. Perez-Lara *et al.*, PtdInsP2 and PtdSer cooperate to trap synaptotagmin-1 to the plasma membrane in the presence of calcium. *Elife* **5** (2016).
26. R. Rand, V. Parsegian, Hydration forces between phospholipid bilayers. *Biochimica et Biophysica Acta -Reviews on Biomembranes* **988**, 351-376 (1989).
27. V. Knecht, H. Grubmuller, Mechanical coupling via the membrane fusion SNARE protein syntaxin 1A: a molecular dynamics study. *Biophys J* **84**, 1527-1547 (2003).
28. Y. Gao *et al.*, Single reconstituted neuronal SNARE complexes zipper in three distinct stages. *Science* **337**, 1340-1343 (2012).
29. A. Stein, G. Weber, M. C. Wahl, R. Jahn, Helical extension of the neuronal SNARE complex into the membrane. *Nature* **460**, 525-U105 (2009).
30. R. Voleti, K. Jaczynska, J. Rizo, Ca(2+)-dependent release of Synaptotagmin-1 from the SNARE complex on phosphatidylinositol 4,5-bisphosphate-containing membranes. *Elife* **9** (2020).
31. K. Grushin *et al.*, Structural basis for the clamping and Ca(2+) activation of SNARE-mediated fusion by synaptotagmin. *Nat Commun* **10**, 2413 (2019).
32. C. Nanavati, V. S. Markin, A. F. Oberhauser, J. M. Fernandez, The Exocytotic Fusion Pore Modeled as a Lipidic Pore. *Biophysical Journal* **63**, 1118-1132 (1992).
33. J. Bai, W. C. Tucker, E. R. Chapman, PIP2 increases the speed of response of synaptotagmin and steers its membrane-penetration activity toward the plasma membrane. *Nat Struct Mol Biol* **11**, 36-44 (2004).
34. A. F. Davis *et al.*, Kinetics of synaptotagmin responses to Ca2+ and assembly with the core SNARE complex onto membranes. *Neuron* **24**, 363-376 (1999).
35. S. Wang, Y. Li, C. Ma, Synaptotagmin-1 C2B domain interacts simultaneously with SNAREs and membranes to promote membrane fusion. *Elife* **5** (2016).
36. Q. Zhou *et al.*, The primed SNARE-complexin-synaptotagmin complex for neuronal exocytosis. *Nature* **548**, 420-425 (2017).
37. K. Mitra, I. Ubarretxena-Belandia, T. Taguchi, G. Warren, D. M. Engelman, Modulation of the bilayer thickness of exocytic pathway membranes by membrane proteins rather than cholesterol. *Proc. Natl. Acad. Sci. U. S. A.* **101**, 4083-4088 (2004).
38. F. Brochard, J. J. J. d. P. Lennon, Frequency spectrum of the flicker phenomenon in erythrocytes. **36**, 1035-1047 (1975).
39. G. Khelashvili, B. Kollmitzer, P. Heftberger, G. Pabst, D. Harries, Calculating the Bending Modulus for Multicomponent Lipid Membranes in Different Thermodynamic Phases. *J. Chem. Theory Comput.* **9**, 3866-3871 (2013).
40. D. Marsh, Elastic curvature constants of lipid monolayers and bilayers. *Chem. Phys. Lipids* **144**, 146-159 (2006).
41. F. S. Cohen, G. B. Melikyan, The energetics of membrane fusion from binding, through hemifusion, pore formation, and pore enlargement. *J Membr Biol* **199**, 1-14 (2004).
42. Z. Wu *et al.*, Dilation of fusion pores by crowding of SNARE proteins. *Elife* **6** (2017).
43. J. Bai, C. T. Wang, D. A. Richards, M. B. Jackson, E. R. Chapman, Fusion pore dynamics are regulated by synaptotagmin*t-SNARE interactions. *Neuron* **41**, 929-942 (2004).

### Figure S1

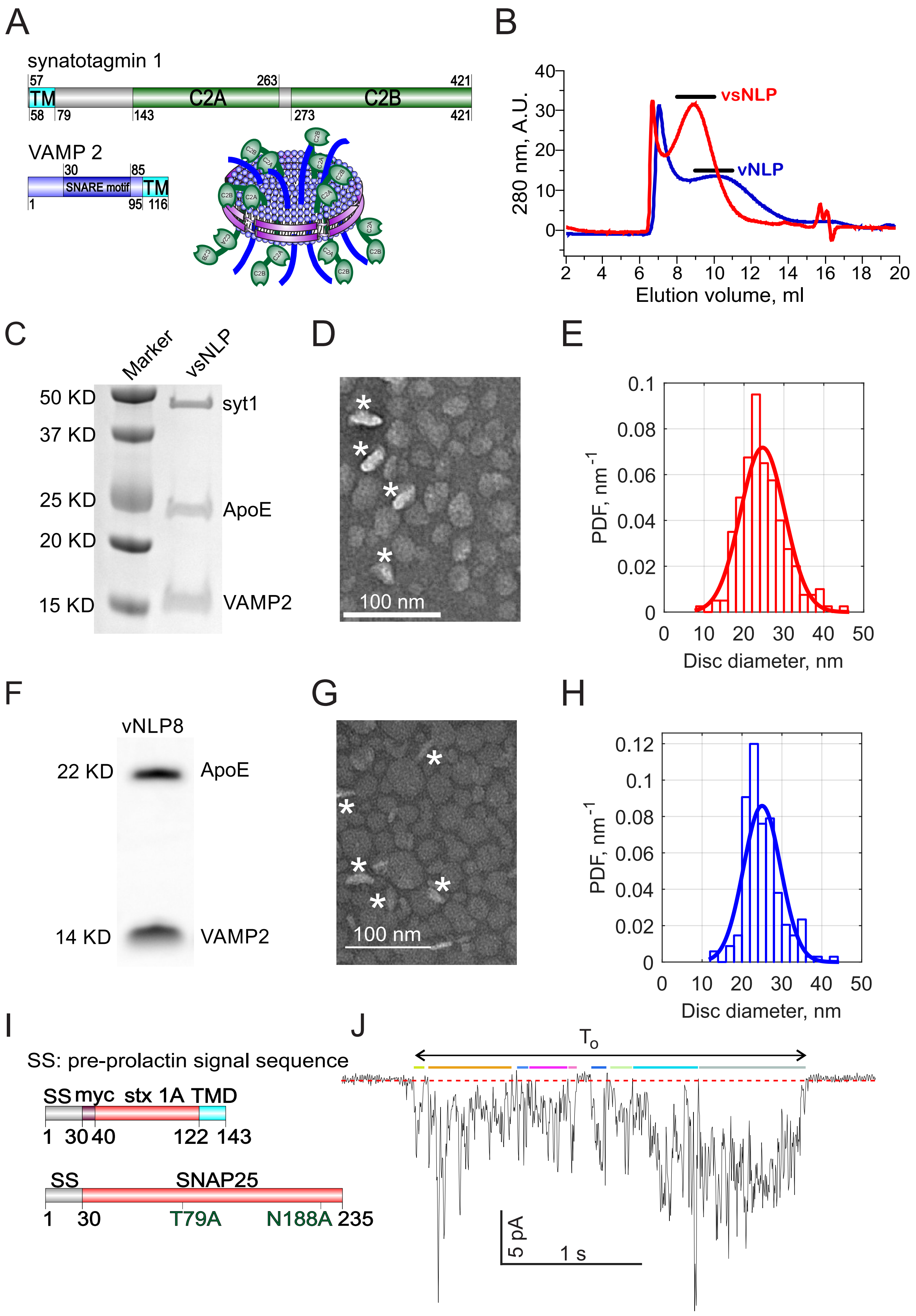
A
synatotagmin 1
57
263
421
TM
C2A
C2B
58
79
143
273
421
VAMP 2
30
85
SNARE motif
TM
1
95
116
B
280 nm, A.U.
vsNLP
vNLP
Elution volume, ml
C
Marker
vsNLP
50 KD
37 KD
25 KD
20 KD
15 KD
syt1
ApoE
VAMP2
D
100 nm
E
PDF, nm-1
Disc diameter, nm
F
vNLP8
22 KD
14 KD
ApoE
VAMP2
G
100 nm
H
PDF, nm-1
Disc diameter, nm
I
SS: pre-prolactin signal sequence
SS myc stx 1A TMD
1 30 40 122 143
SS SNAP25
1 30 T79A N188A 235
J
To
5 pA
1 s

### Figure S2

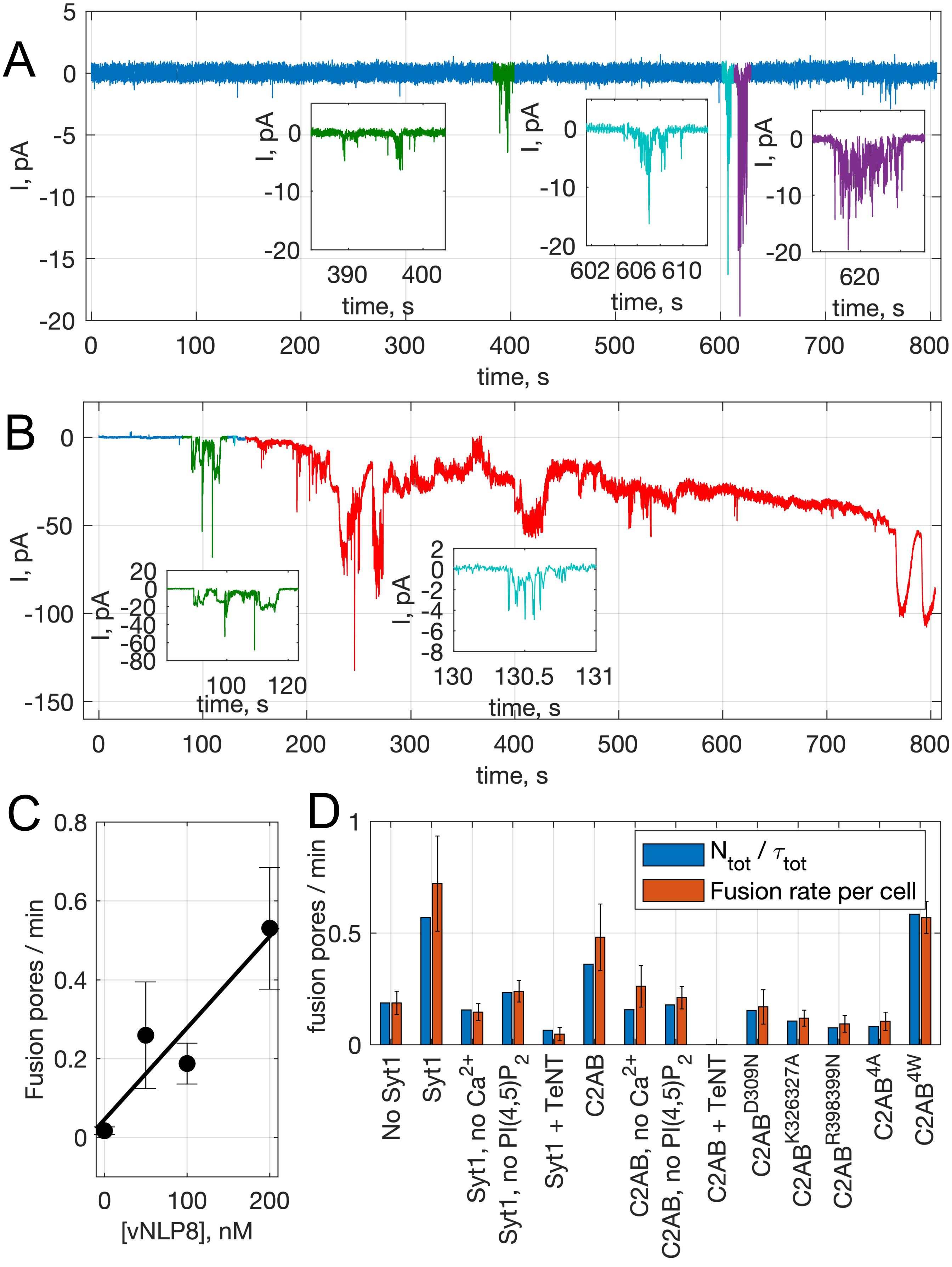

A
I, pA
time, s
B
I, pA
time, s
C
Fusion pores / min
[vNLP8], nM
D
fusion pores / min
N tot / τ tot
Fusion rate per cell
No Syt1
Syt1
Syt1, no Ca2+
Syt1, no PI(4,5)P2
Syt1 + TeNT
C2AB
C2AB, no Ca2+
C2AB, no PI(4,5)P2
C2AB + TeNT
C2AB D309N
C2AB K326327A
C2AB R398399N
C2AB 4A
C2AB 4W

### Figure S3

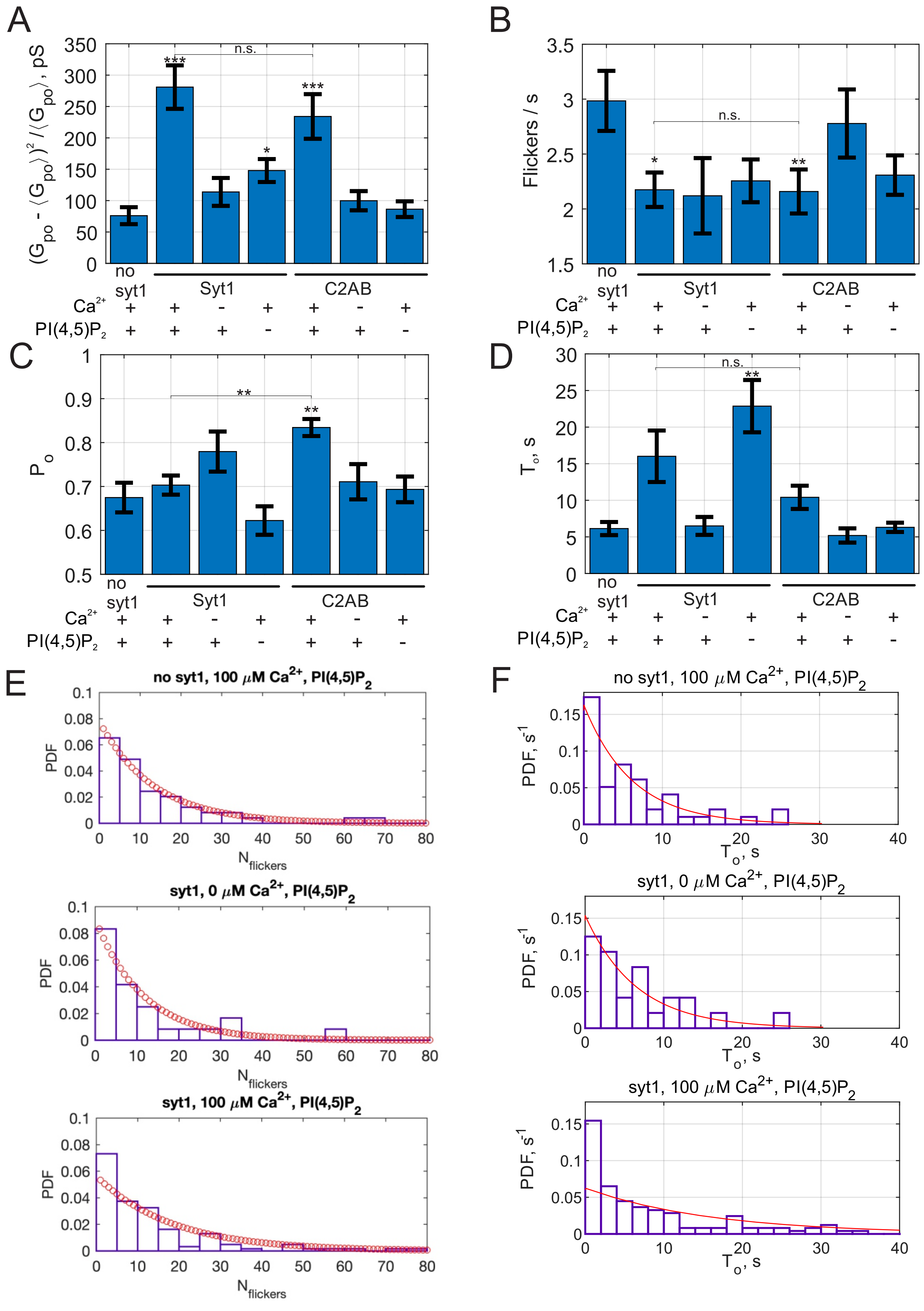
A
(G_po - ⟨G_po⟩)^2 / ⟨G_po⟩, pS
n.s.
***
*
no syt1
Syt1
C2AB
Ca2+
PI(4,5)P2
B
Flickers / s
n.s.
C
P_o
**
D
T_o, s
E
no syt1, 100 μM Ca2+, PI(4,5)P2
syt1, 0 μM Ca2+, PI(4,5)P2
syt1, 100 μM Ca2+, PI(4,5)P2
PDF
N_flickers
F
PDF, s-1
T_o, s

### Figure S4

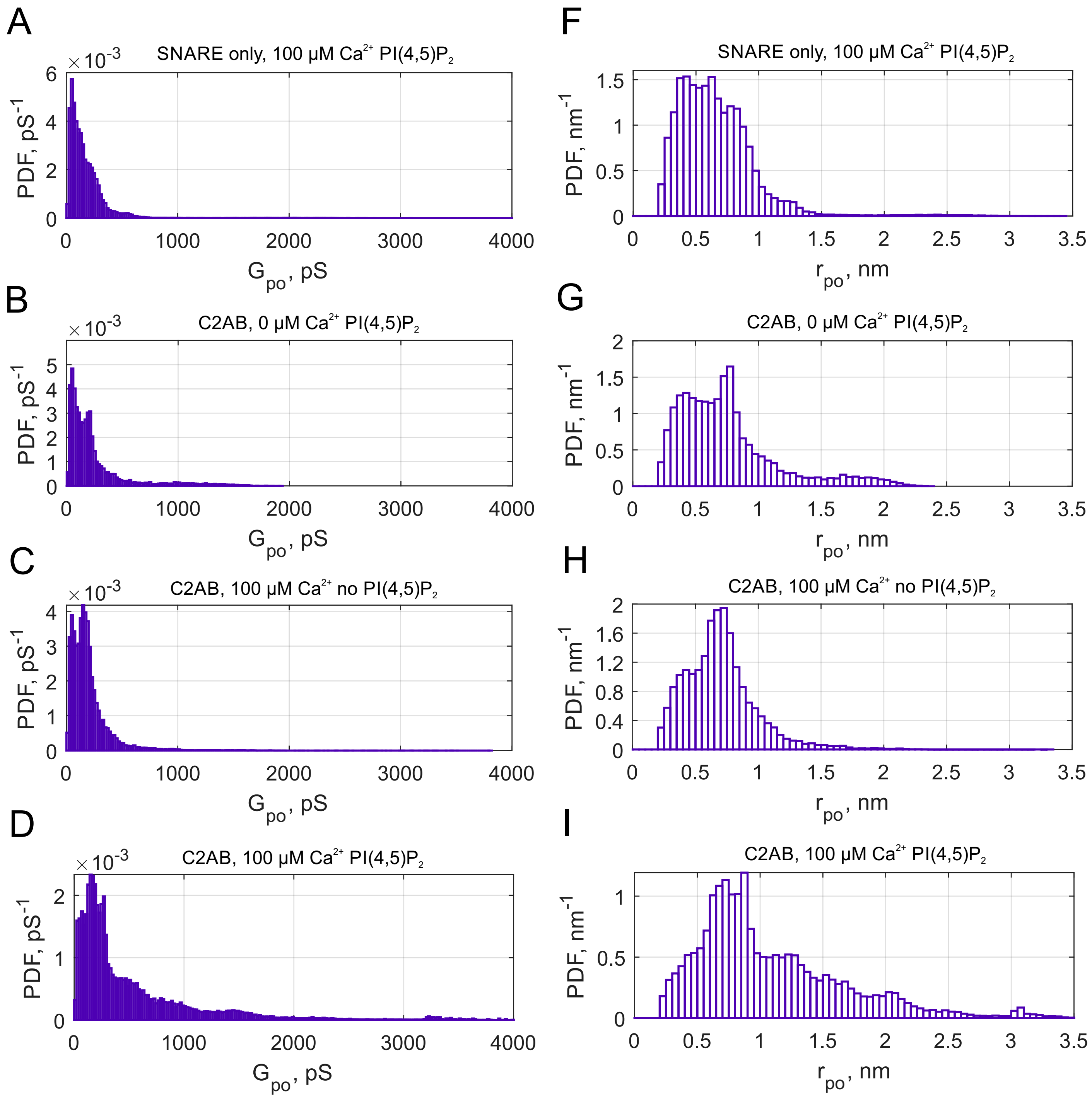

A
SNARE only, 100 μM Ca2+ PI(4,5)P2
PDF, pS-1
Gpo, pS
B
C2AB, 0 μM Ca2+ PI(4,5)P2
PDF, pS-1
Gpo, pS
C
C2AB, 100 μM Ca2+ no PI(4,5)P2
PDF, pS-1
Gpo, pS
D
C2AB, 100 μM Ca2+ PI(4,5)P2
PDF, pS-1
Gpo, pS
F
SNARE only, 100 μM Ca2+ PI(4,5)P2
PDF, nm-1
rpo, nm
G
C2AB, 0 μM Ca2+ PI(4,5)P2
PDF, nm-1
rpo, nm
H
C2AB, 100 μM Ca2+ no PI(4,5)P2
PDF, nm-1
rpo, nm
I
C2AB, 100 μM Ca2+ PI(4,5)P2
PDF, nm-1
rpo, nm

### Figure S5

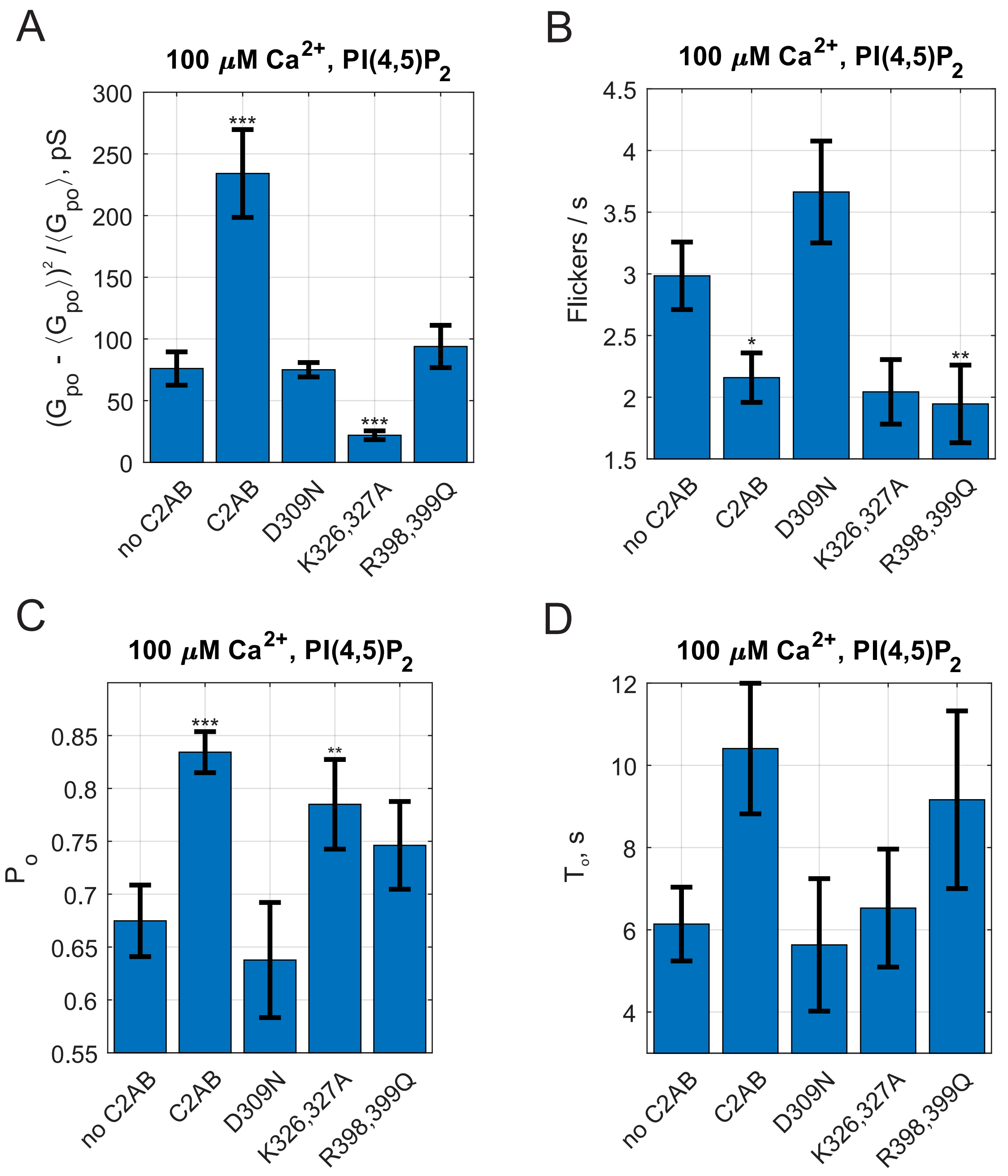
A
100 μM Ca2+, PI(4,5)P2
(Gpo - ⟨Gpo⟩)2 / ⟨Gpo⟩, pS
300
250
200
150
100
50
0
***
***
no C2AB
C2AB
D309N
K326,327A
R398,399Q
B
100 μM Ca2+, PI(4,5)P2
Flickers / s
4.5
4
3.5
3
2.5
2
1.5
*
**
no C2AB
C2AB
D309N
K326,327A
R398,399Q
C
100 μM Ca2+, PI(4,5)P2
Po
0.85
0.8
0.75
0.7
0.65
0.6
0.55
***
**
no C2AB
C2AB
D309N
K326,327A
R398,399Q
D
100 μM Ca2+, PI(4,5)P2
To, s
12
10
8
6
4
no C2AB
C2AB
D309N
K326,327A
R398,399Q

### Figure S6

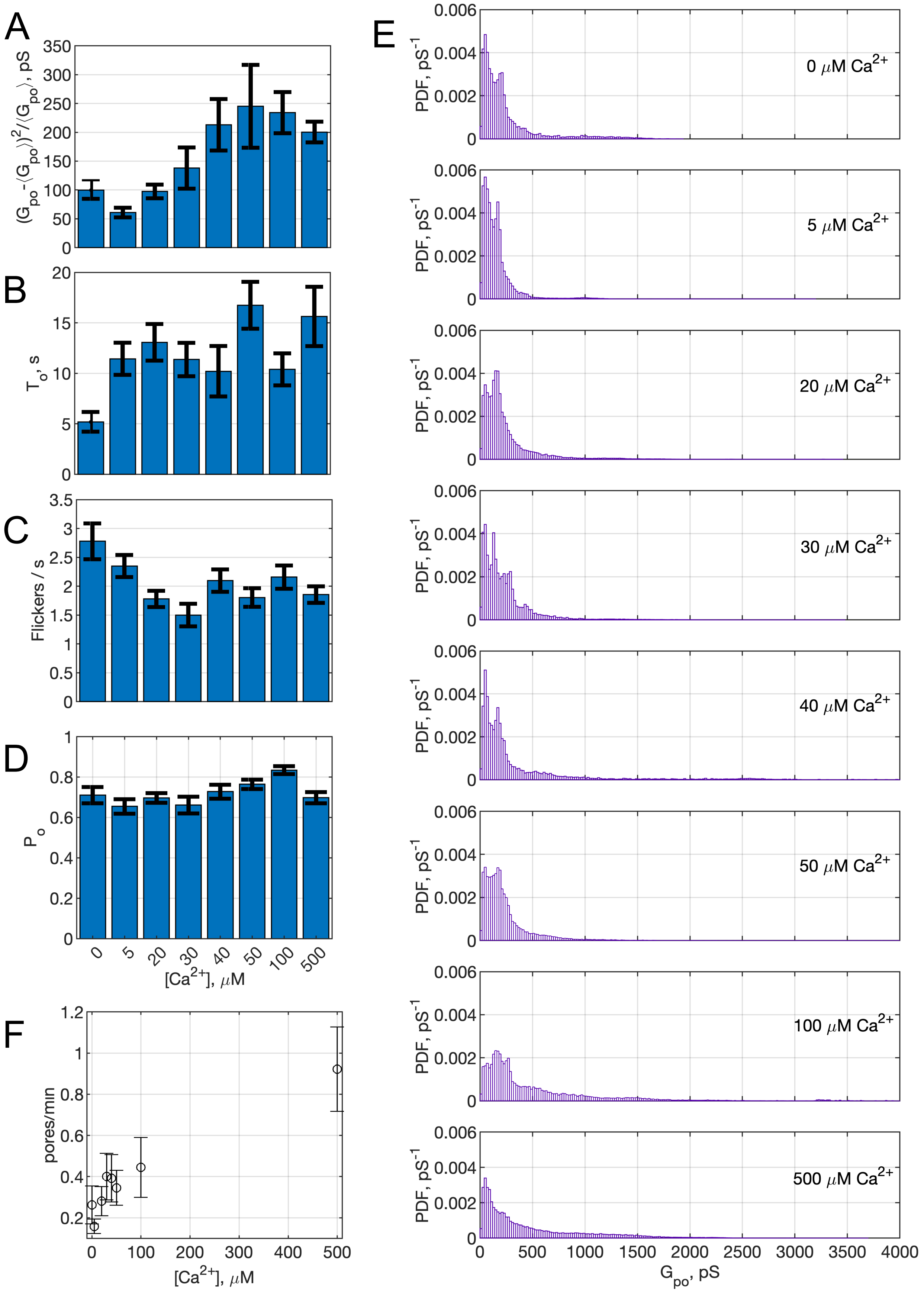

A
$(G_{po}-\langle G_{po}\rangle)^2/\langle G_{po}\rangle$, pS
B
$T_o$, s
C
Flickers / s
D
$P_o$
$[Ca^{2+}]$, $\mu$M
0
5
20
30
40
50
100
500
F
pores/min
$[Ca^{2+}]$, $\mu$M
E
PDF, $pS^{-1}$
0 $\mu$M $Ca^{2+}$
5 $\mu$M $Ca^{2+}$
20 $\mu$M $Ca^{2+}$
30 $\mu$M $Ca^{2+}$
40 $\mu$M $Ca^{2+}$
50 $\mu$M $Ca^{2+}$
100 $\mu$M $Ca^{2+}$
500 $\mu$M $Ca^{2+}$
$G_{po}$, pS

### Figure S7

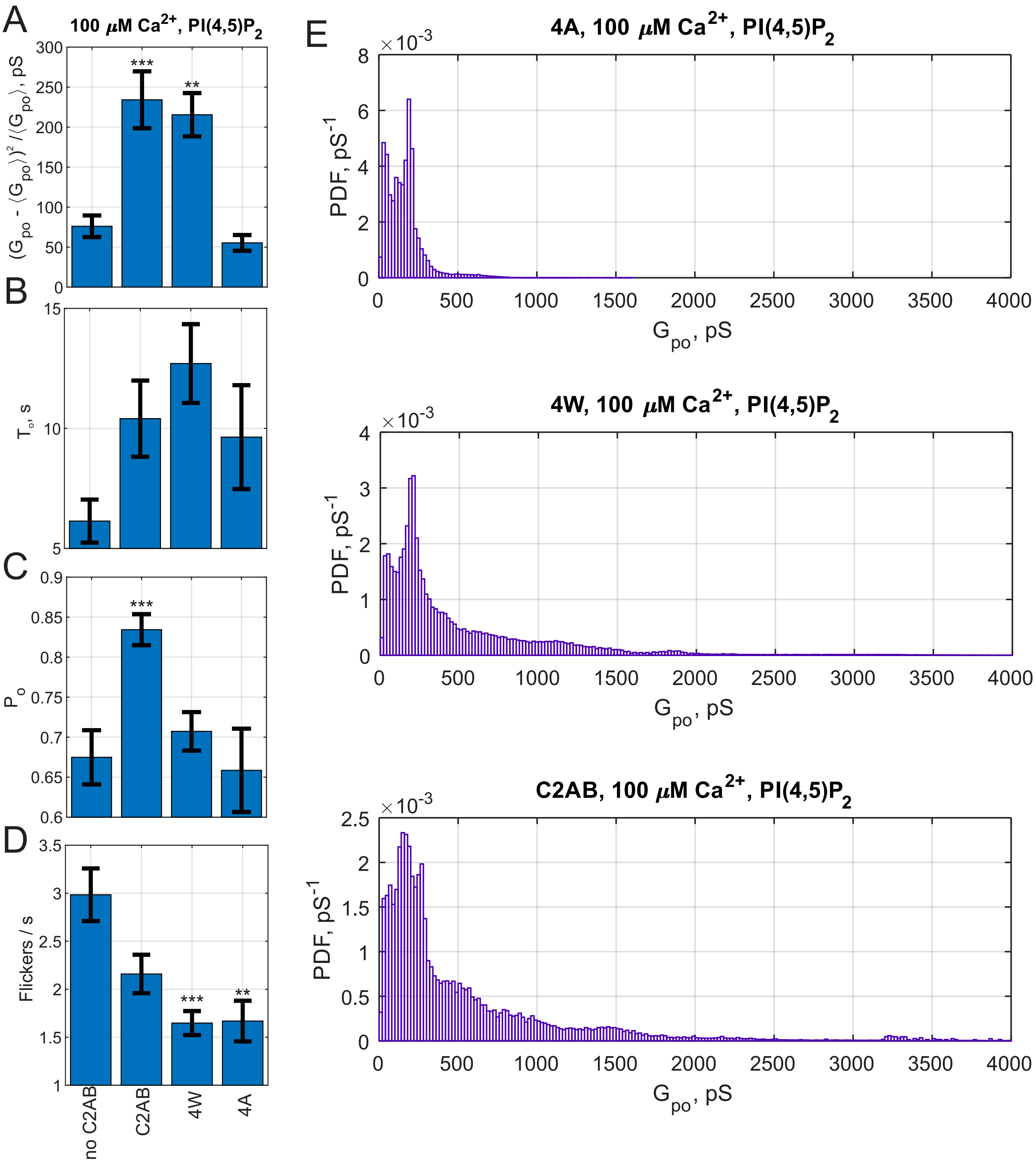
A
100 μM Ca2+, PI(4,5)P2
(Gpo - ⟨Gpo⟩)² / ⟨Gpo⟩, pS
***
**
B
To, s
C
Po
***
D
Flickers / s
***
**
no C2AB
C2AB
4W
4A
E
4A, 100 μM Ca2+, PI(4,5)P2
×10-3
PDF, pS-1
Gpo, pS
4W, 100 μM Ca2+, PI(4,5)P2
×10-3
PDF, pS-1
Gpo, pS
C2AB, 100 μM Ca2+, PI(4,5)P2
×10-3
PDF, pS-1
Gpo, pS

### Figure S8

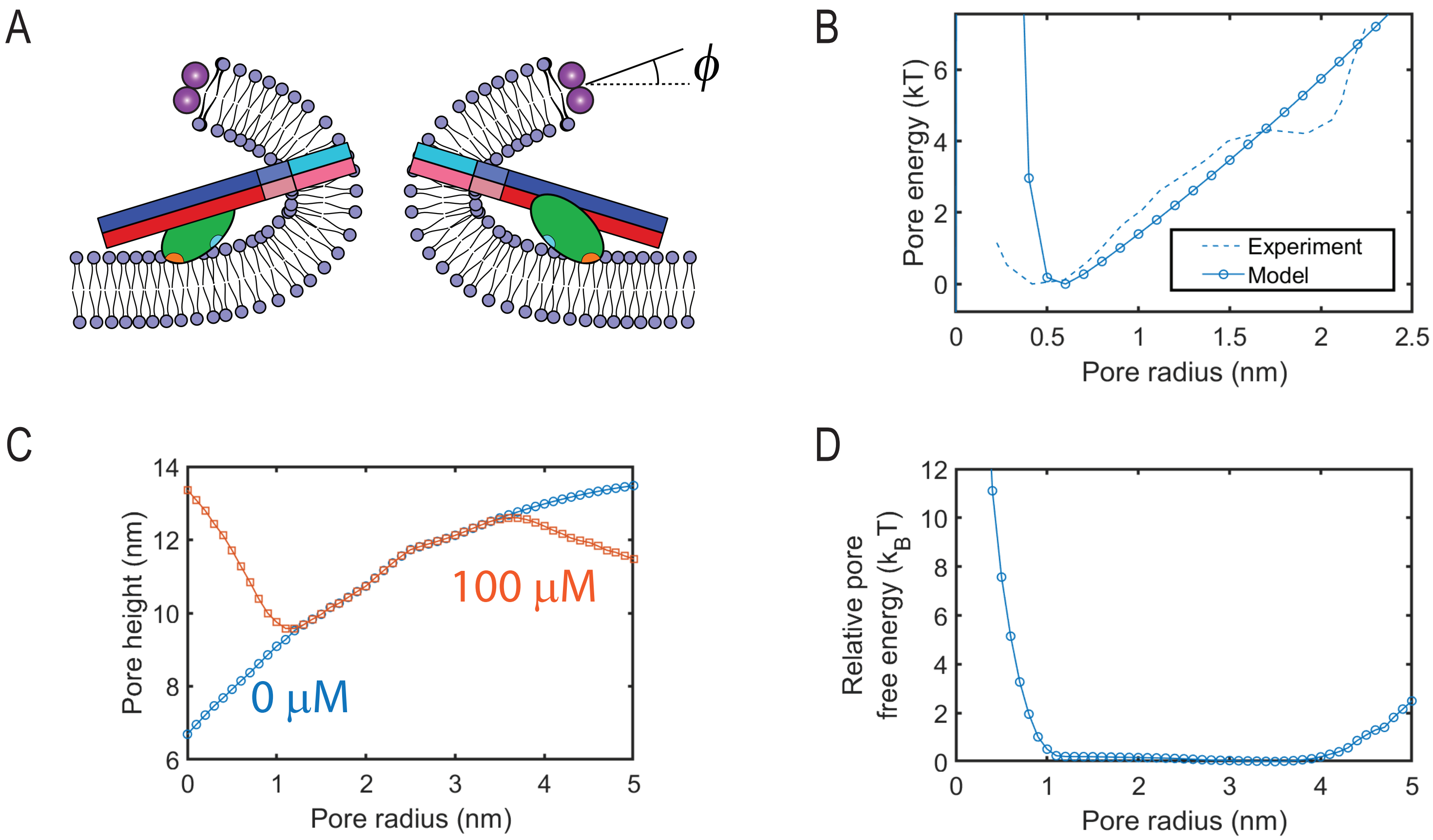

A
$\phi$
B
Pore energy (kT)
Pore radius (nm)
Experiment
Model
C
Pore height (nm)
Pore radius (nm)
100 µM
0 µM
D
Relative pore free energy ($k_BT$)
Pore radius (nm)

### Figure S9

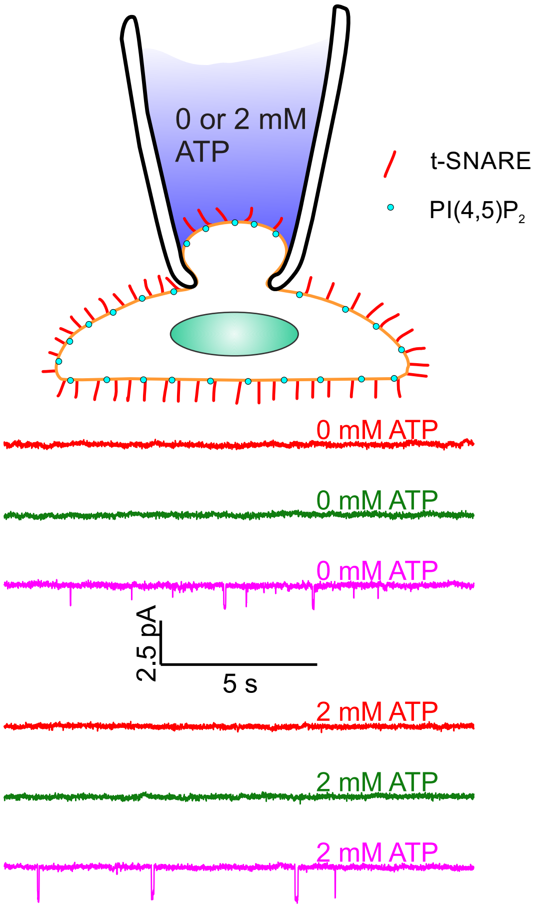
0 or 2 mM
ATP
t-SNARE
PI(4,5)P2
0 mM ATP
0 mM ATP
0 mM ATP
2.5 pA
5 s
2 mM ATP
2 mM ATP
2 mM ATP

### Figure S10

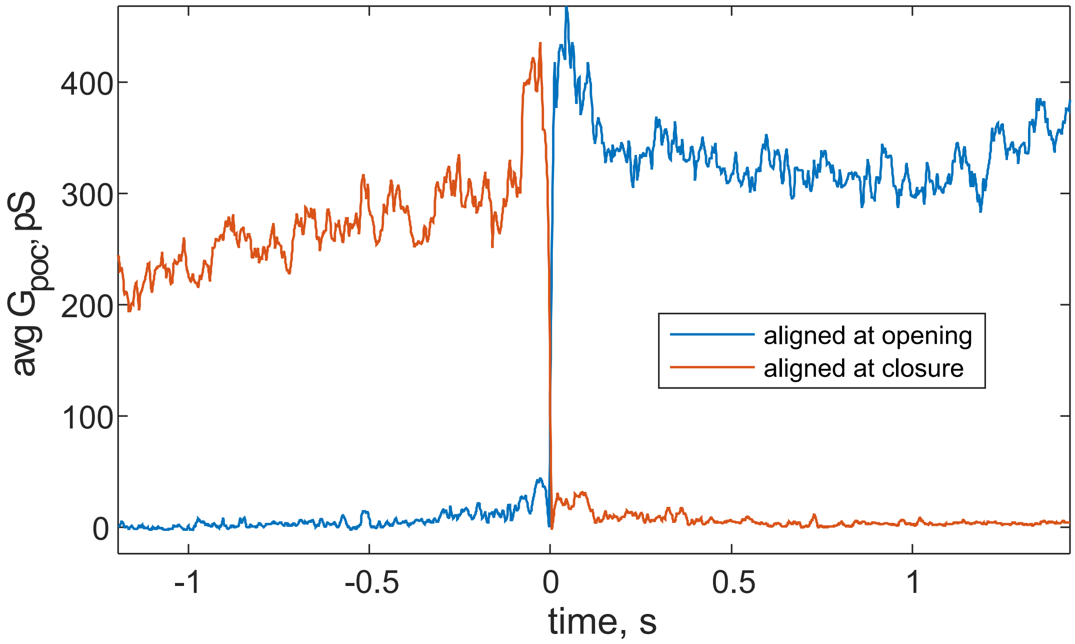
avg $G_{poc}$, pS
400
300
200
100
0
-1
-0.5
0
0.5
1
time, s
aligned at opening
aligned at closure

### Figure S11

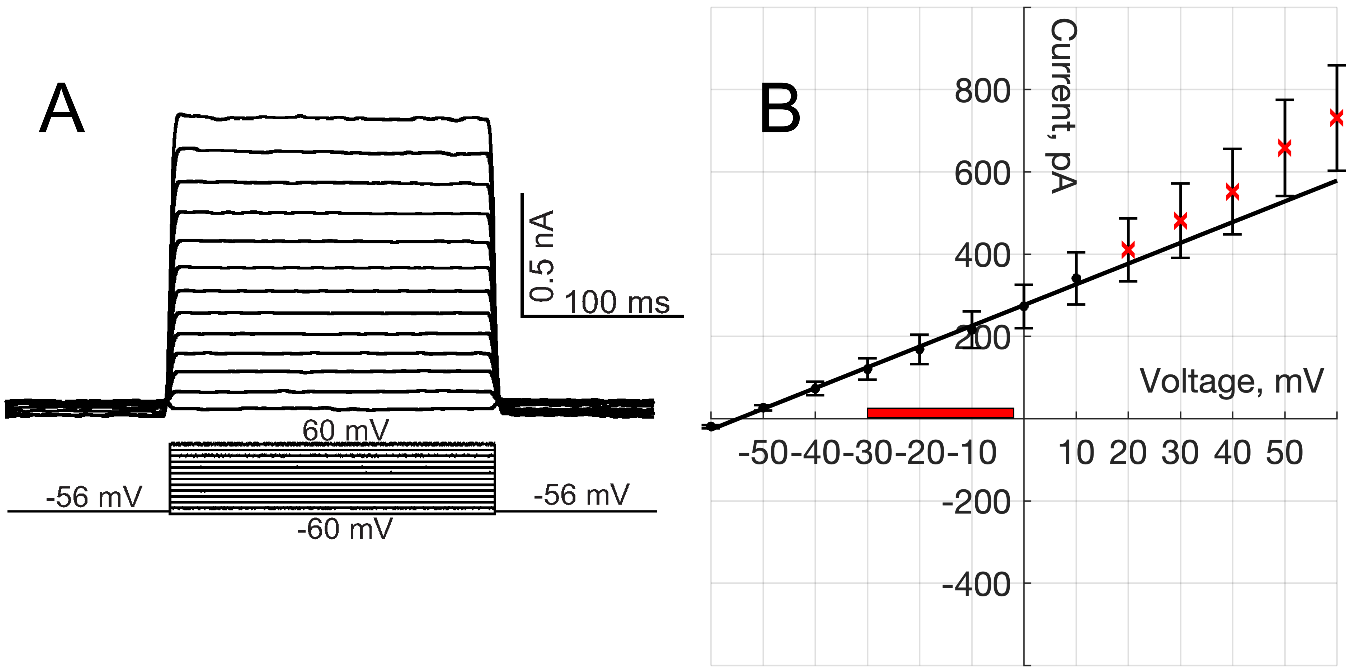

A
0.5 nA
100 ms
60 mV
-56 mV
-56 mV
-60 mV
B
Current, pA
800
600
400
200
-200
-400
Voltage, mV
-50-40-30-20-10
10 20 30 40 50
